# The E3 ubiquitin ligase mechanism specifying target-directed microRNA degradation

**DOI:** 10.64898/2026.01.05.697729

**Authors:** Jakob Farnung, Elena Slobodyanyuk, Peter Y. Wang, Lianne W. Blodgett, Daniel H. Lin, Susanne von Gronau, Brenda A. Schulman, David P. Bartel

**Affiliations:** Department of Molecular Machines and Signaling, Max Planck Institute of Biochemistry, Martinsried 82152, Germany; Howard Hughes Medical Institute, Cambridge, MA 02142, USA; Whitehead Institute for Biomedical Research, Cambridge, MA 02142, USA; Department of Biology, Massachusetts Institute of Technology, Cambridge, MA 02139, USA

## Abstract

MicroRNAs (miRNAs) associate with Argonaute (AGO) proteins to form complexes that down-regulate target RNAs, including mRNAs from most human genes^1–3^. Within each complex, the miRNA pairs to target mRNAs to specify their repression, and AGO provides effector function while also protecting the miRNA from cellular nucleases^2–5^. Although much has been learned about this mode of posttranscriptional gene regulation, less is known about how the miRNAs themselves are regulated. In one such regulatory pathway, unusual miRNA targets called “trigger” RNAs reverse the canonical regulatory logic and instead down-regulate microRNAs^6–21^. This target-<u>d</u>irected <u>m</u>iRNA <u>d</u>egradation (TDMD) is thought to require a <u>c</u>ullin–<u>R</u>ING E3 ligase (CRL) because it depends on the cullin protein CUL3 and other ubiquitylation components, including the BC-box protein ZSWIM8 (ref. 22,23). ZSWIM8 is required for murine perinatal viability and for destabilization of most short-lived miRNAs, but is otherwise poorly understood^23–25^. Here, we demonstrate that a human AGO–miRNA– trigger complex selectively binds ZSWIM8 for CUL3-mediated polyubiquitylation of the AGO protein within this complex. Cryogenic electron-microscopy (cryo-EM) analyses show how ZSWIM8 recognizes the distinct AGO2 and miRNA–trigger conformations shaped by pairing of the miRNA to the trigger. For example, this pairing extracts the miRNA from a binding pocket within AGO2, allowing the pocket to be captured by ZSWIM8, and it directs the trigger RNA along a distinct trajectory to be also recognized by ZSWIM8. These results biochemically establish AGO binding and polyubiquitylation as the key regulatory step of TDMD, define a unique CRL class, and reveal generalizable RNA–RNA, RNA–protein, and protein–protein interactions that specify the ubiquitin-mediated degradation of AGO with exquisite selectivity. The substrate features recognized by the E3 ubiquitin ligase do not conform to a conventional degron^26–28^, but rather establish a two-RNA-factor authentication mechanism specifying a protein ubiquitylation substrate.

## Introduction

Metazoan miRNAs recognize their mRNA targets primarily through pairing between the miRNA “seed” region (miRNA nucleotides 2–8) and sites within the mRNA 3′ untranslated regions (UTRs)^29^. In contrast, the unusual transcripts that trigger miRNA degradation not only pair to the miRNA seed region but, following an internal loop involving the miRNA central region, also pair extensively to the miRNA 3′ region^30–34^. Although these trigger sites presumably afford greater affinity to the AGO–miRNA complex due to pairing to the miRNA 3′ region^35^, this increased affinity is insufficient to explain the selectivity of the TDMD machinery for the vastly outnumbered trigger-bound complexes. Indeed, for a miRNA undergoing TDMD, trigger sites in the cellular transcriptome typically number ∼100 (ref. 12,23), which, when compared to the ∼100,000 seed-matched sites residing within mRNA 3′ UTRs^36^, present a formidable stoichiometric challenge for molecular recognition.

To this point, our understanding of TDMD has come primarily from genetic analyses, which have revealed the key factors involved in TDMD and the molecular and physiological consequences of losing those factors. This pathway appears to affect many metazoan miRNAs. Over 50 ZSWIM8- sensitive miRNAs have been identified in human cell lines, over 50 have been identified in mouse tissues, 21 have been identified in Drosophila embryos and cells, and 22 have been identified in nematodes^17,20,21,23–25,37^. Additionally, some herpesviruses express trigger RNAs that direct the decay of host miRNAs^7–10^. The breadth and magnitude of ZSWIM8-mediated miRNA regulation presumably underlie the lethality of ZSWIM8 knockout in both flies and mice^17,24,25^, although other diverse functions have also been proposed for ZSWIM8^38–46^.

In the current model for TDMD, the binding of a trigger RNA to the AGO–miRNA complex causes a conformational change that is recognized by a ZSWIM8–CUL3 CRL through an unknown mechanism (Figure 1a)^14,22,23^. This CRL, together with the ARIH1 RBR-type E3 ligase and cognate E2s^22^, presumably polyubiquitylates AGO^47–49^, causing its degradation by the 26S proteasome and ultimately leaving the miRNA susceptible to cellular nucleases^22,23^. The ZSWIM8 CRL belongs to a mix-and-match system of substrate-binding receptors with common ubiquitylation components that assemble in various combinations to form hundreds of different E3 ligases^50,51^. The CRL that achieves TDMD has some unprecedented features. Genetic data implicate ZSWIM8 and its obligate partner proteins ELOB and ELOC (hereafter referred to collectively as ZSWIM8, except when referring to the *ZSWIM8* gene) as the substrate-binding receptor of a CUL3-based CRL^22,23,38^. However, how these proteins partner within a CRL is not clear; BC-box proteins such as ZSWIM8 are only known to function with CUL2 or CUL5^52,53^, whereas CUL3 is thought to partner exclusively with the BTB family of substrate-binding receptors^54–57^. Moreover, for TDMD, this mix-and-match system of proteins is further elaborated with a mix-and-match system of RNAs—the miRNAs and trigger RNAs—that promote polyubiquitylation of their common protein partner. Here we show how these RNAs provide a unique two-RNA-factor authentication mechanism specifying a ubiquitylation substrate.

**Figure 1:**
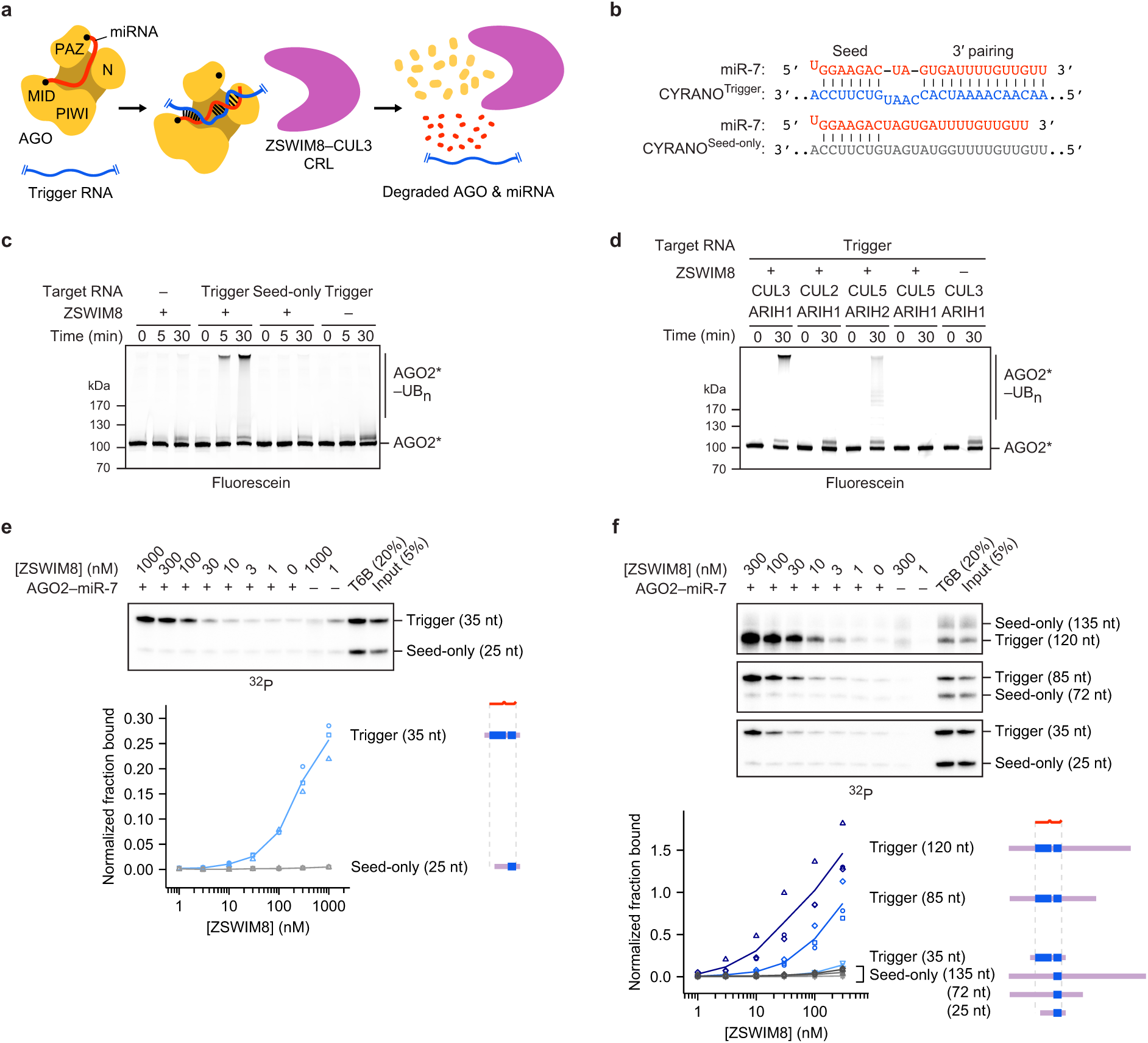
Trigger RNA specifies AGO ubiquitylation by the ZSWIM8–CUL3 E3 ligase through selective ZSWIM8 binding. **a,** Schematic of the proposed mechanism for TDMD. See main text for description. **b,** Diagrams showing pairing between miR-7 and the miRNA-binding regions of CYRANO^Trigger^ and CYRANO^Seed-only^ target RNAs used for *in vitro* ubiquitylation and co-IP assays. Vertical lines represent Watson–Crick–Franklin (W–C–F) pairing. **c,** Biochemical reconstitution of trigger- and ZSWIM8-dependent ubiquitylation of AGO2 using the miR-7–CYRANO miRNA–trigger pair. CYRANO trigger (120 nt) and CYRANO seed-only (135 nt) target RNAs were used (Supplementary Table 1). Polyubiquitylated products (AGO2*–UB_n_) were resolved by SDS-PAGE and detected by in-gel visualization of fluorescently labeled AGO2 (AGO2*). Shown is a representative experiment; *n* = 2 technical replicates. Gels stained with Coomassie blue are shown in Supplementary Figure 1. **d,** *In vitro* ubiquitylation of AGO2, comparing the activity of different cullin–ubiquitin-carrying-enzyme pairs. CUL3 and CUL2 canonically function with ARIH1, whereas CUL5 functions with ARIH2, a close paralog of ARIH1. AGO2 was complexed with the miR-7–CYRANO miRNA–trigger pair. Otherwise, this panel is as in **c**. Shown is a representative experiment; *n* = 2 technical replicates. Gels stained with Coomassie blue are shown in Supplementary Figure 1. **e,** Biochemical reconstitution of ZSWIM8 binding to AGO2–miR-7–CYRANO^Trigger^ complexes. Top: *In vitro* co-IP assay (Extended Data Figure 1a). AGO2–miR-7 was preincubated with radiolabeled target RNAs derived from the CYRANO trigger and then incubated with an excess of purified epitope-tagged ZSWIM8. These target RNAs harbored either trigger or seed-only pairing to miR-7 (35- and 25-nt RNAs, respectively) (Supplementary Table 1). RNAs that co-IPed with ZSWIM8 were resolved on a denaturing gel, and radioactivity was visualized by phosphor imaging. Bottom: Quantification of radiolabeled co-IPed RNA. Values were background-subtracted using values for the 0 nM ZSWIM8 sample and then normalized to those of the T6B sample. The symbols show data from independent measurements; lines pass through mean values. *n* = 3 technical replicates. At the right are schematics of the RNAs, indicating the degree of miRNA pairing (blue) and relative lengths of flanking regions (purple). This quantification revealed up to 70-fold preference for co-IP of target RNAs with trigger pairing over seed-only pairing. **f,** Effect of RNA flanking the miRNA-binding site on ZSWIM8 binding to AGO2–miR-7–CYRANO. Top: *In vitro* co-IP assay as in **e**, but target RNAs had additional CYRANO sequence flanking the miRNA-binding site, and heparin (1 µg/mL) was included to reduce non-specific binding of longer RNAs to the beads. Bottom: Quantification of radiolabeled co-IPed RNA, as in **e**. *n* = 3 technical replicates.

## Results

### Trigger RNA specifies AGO ubiquitylation

Despite genetic and molecular evidence supporting an E3 model for TDMD^22,23^, and despite the centrality of AGO polyubiquitylation in this model (Figure 1a), ubiquitylation of TDMD-competent AGO had not been reported. We purified the genetically defined factors implicated in TDMD and assayed their collective ability to ubiquitylate AGO2. Fluorescent AGO2 preloaded with miR-7 was polyubiquitylated in a ZSWIM8-, CUL3-, and ARIH1-dependent manner only if also incubated with a 120-nt fragment of CYRANO, the trigger RNA of human miR-7 (ref. 12) (Figure 1b,c, Supplementary Table 1). A mutant (“seed-only”) version of the CYRANO fragment, in which pairing to the 3′ region of miR-7 was disrupted, failed to specify polyubiquitylation. The ZSWIM8–CUL3–ARIH1 E3–E3 assembly was required, as polyubiquitylation was not achieved with other canonical CRL–RBR E3–E3 partnerings (ZSWIM8–CUL2–ARIH1 or ZSWIM8–CUL5–ARIH2) (Figure 1d)^48,49,58^. Thus, an AGO2 associated with a miRNA is a direct substrate for ZSWIM8– CUL3-mediated polyubiquitylation, but only when bound to a trigger RNA.

### ZSWIM8 binds AGO–miRNA–trigger ternary complex

Our *in vitro* ubiquitylation assay showed that components present in our purified system were sufficient for substrate selectivity. One mechanism for achieving this selectivity would be through preferential binding of ZSWIM8 to the AGO2–miRNA–trigger ternary complex (Figure 1a). We investigated this possibility with an *in vitro* co-immunoprecipitation (co-IP) assay designed to detect binding of the ternary complex. An excess of purified epitope-tagged ZSWIM8 was incubated with AGO2–miR-7 that had been mixed with two radiolabeled target-RNA species, which harbored either the native CYRANO site or a seed-only site. These RNAs were of different sizes (35 and 25 nt, respectively), such that RNA species co-purifying with ZSWIM8 could be resolved on a denaturing gel and quantified (Extended Data Figure 1a). In parallel, co-IPs were performed with a peptide bait derived from TNRC6, a downstream effector of AGO^2^. This peptide (called “T6B”) binds AGO–miRNA complexes irrespective of their bound target RNAs^59–61^, and it was used to normalize for any differences in AGO2–miR-7 binding to different target RNAs (Extended Data Figure 1a).

ZSWIM8 preferentially co-IPed with the AGO2–miR-7–trigger complex, with up to 70-fold enrichment of the trigger-bound complex over its seed-only counterpart (Figure 1e). As another specificity control, we tested a target in which seed pairing was supplemented with only seven (instead of 14) nucleotides of complementarity to the miRNA 3′ region. This limited 3′ pairing was designed to represent “3′-supplementary” pairing, a type of pairing that increases affinity to the miRNA but is typically insufficient to direct miRNA degradation^30–34^. As expected, the target with 3′-supplementary pairing was unable to promote ZSWIM8 co-IP (Extended Data Figure 1b,c). Together, these results indicated that preferential ZSWIM8 binding to the AGO2–miRNA–trigger complex underpins much of the selective activity observed in the *in vitro* ubiquitylation assay, and presumably also in cells.

We next examined whether these binding and polyubiquitylation activities were generalizable to other TDMD-competent AGO–miRNA complexes. Indeed, ZSWIM8 displayed binding and polyubiquitylation activity toward another TDMD-competent AGO paralog, AGO1 (ref. 23) (Extended Data Figure 1d,e), and toward another miRNA–trigger pair, miR-27a–HSUR1, which represents a founding example of TDMD, in which a viral noncoding RNA directs the degradation of a host miRNA^7^ (Extended Data Figure 1f–h). Importantly, as in the regulation observed *in vivo*, cognate miRNA–trigger pairing was required in our biochemical reconstitution; miR-27a trigger HSUR1 failed to elicit polyubiquitylation of AGO2–miR-7, and miR-7 trigger CYRANO failed to elicit polyubiquitylation of AGO2–miR-27a (Extended Data Figure 1i). Taken together, these results demonstrate that diverse but specific miRNA–trigger pairs direct the recognition of their common AGO protein partner by an uncharacterized class of CRL.

### Key role of RNA flanking trigger pairing

Previous studies on triggers had largely focused on the miRNA-binding site^6–21,62^. Our *in vitro* binding assay allowed us to assess whether parts of the trigger flanking the miRNA-binding site might also influence ZSWIM8 recognition. Including additional CYRANO sequence flanking the miR-7-binding site, which increased the size of the CYRANO fragment by 85 nt, increased efficiency of AGO2–miR-7– trigger co-IP by 100-fold. For instance, 300 nM ZSWIM8 was required to achieve 15% pulldown in the absence of additional flanking sequence, whereas only 3 nM was required in the presence of additional flanking sequence (Figure 1f). Selectivity remained high when using the longer trigger sequences, with a >100-fold preference observed for the native CYRANO pairing over its seed-only counterpart (Figure 1f, Extended Data Figure 1c). Increased co-IP efficiency was similarly observed for finer-grained extensions of the CYRANO fragment (Extended Data Figure 2a), as well as with AGO2 paralog AGO1 (Extended Data Figure 2b). One explanation for the increased co-IP efficiency would be that trigger RNA flanking regions interact directly with ZSWIM8. Interestingly, the efficiency of co-IP remained high when CYRANO sequences flanking the miR-7 binding site were scrambled (Extended Data Figure 2c,d), which suggested that ZSWIM8 might have some sequence-independent affinity to RNA. Indeed, we detected weak ZSWIM8 binding to RNA alone in filter-binding experiments, with affinity increasing as RNA lengths increased from 28 to 120 nt (Extended Data Figure 2e).

### ZSWIM8 dimer clamps AGO2–miR-7–trigger

To gain structural insights into TDMD substrate recognition, we obtained cryo-EM data for a ZSWIM8–CUL3 complex bound to an AGO2–miR-7–CYRANO complex. The final reconstruction has an overall resolution of 3.1 Å and superior local resolution at key interfaces (Figure 2, Supplementary Table 2, Supplementary Figure 2–4, Supplementary Video 1). Overall, ZSWIM8 forms a dimer, with each protomer projecting an alpha-helical solenoid. Using these solenoids, the two ZSWIM8 protomers form an asymmetric clamp around a single AGO2–miR-7–trigger complex. One protomer, which we call ZSWIM8^NPAZ^, interacts with the N and PAZ domains of AGO2, whereas the other one, which we call ZSWIM8^MID^, interacts with the MID domain of AGO2. As described below, the structure rationalizes the unexpected finding that ZSWIM8 partners with CUL3 to form an E3 ligase, and explains how the trigger RNA orchestrates E3–substrate interactions enabled by its distinct pairing architecture with the miRNA. The signature miRNA–trigger pairing reshapes the conformation of AGO2 and the miRNA–trigger duplex to be compatible with binding ZSWIM8. In doing so, miRNA–trigger pairing renders a pocket within the AGO2 PAZ domain—which is otherwise occupied by the miRNA 3′ end—available to bind ZSWIM8, and guides the flanking regions of the trigger RNA to wrap around the ZSWIM8 clamp and bind positively charged surfaces on ZSWIM8. In this way, disparate parts of the AGO2–miR-7–CYRANO complex present a constellation of ZSWIM8-interacting elements that dictate the selectivity of binding—in a manner that appears to be generalizable across miRNA–trigger pairings.

**Figure 2:**
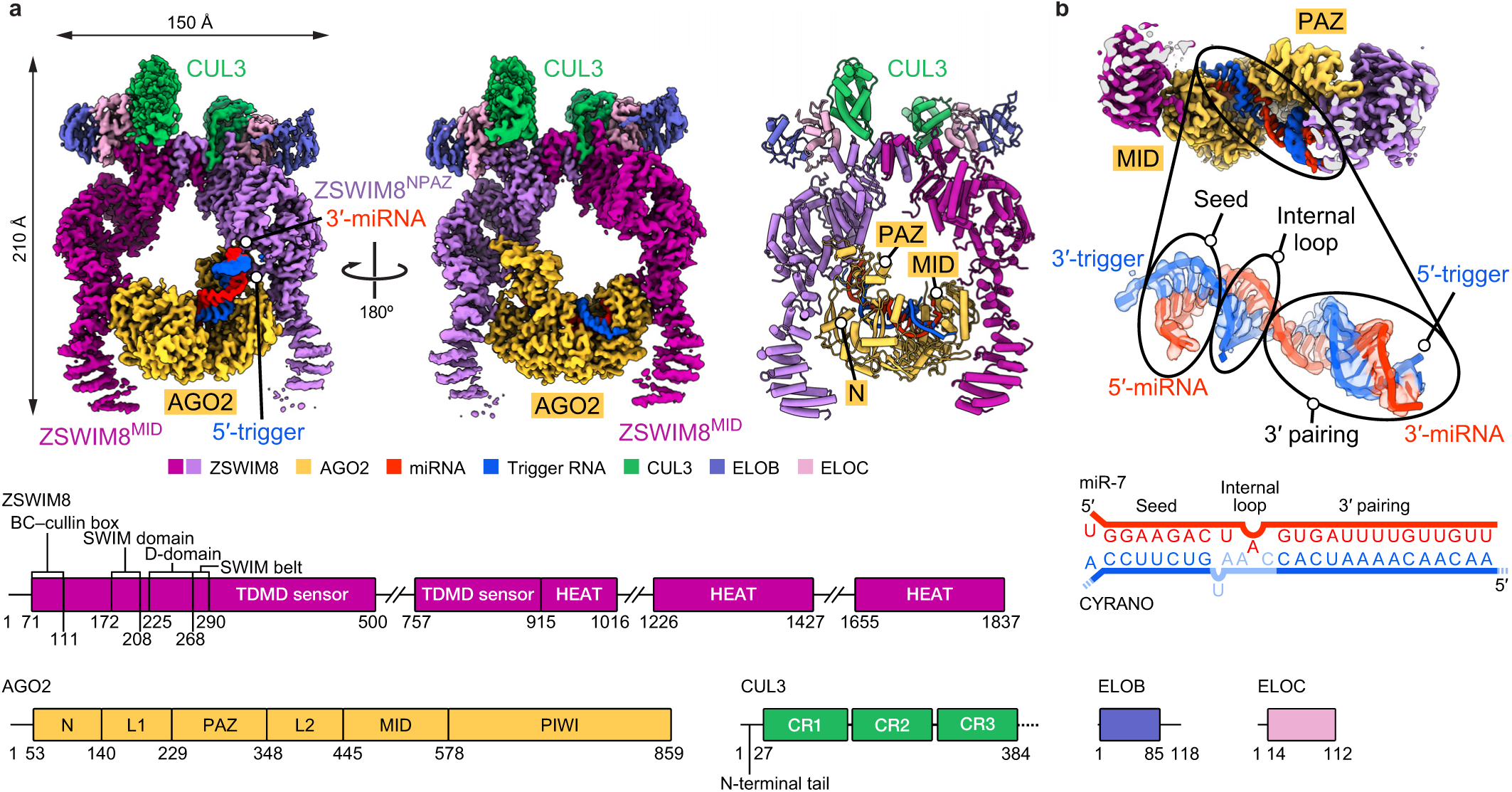
ZSWIM8 dimer clamps around the AGO2–miR-7–trigger complex. **a,** Structure of ZSWIM8–CUL3 bound to AGO2–miR-7–CYRANO. Top left and top middle: Cryo-EM map of the complex. The composite map based on multiple focused-refined maps is shown. Top right: Cartoon representation of the model. Large unstructured regions in ZSWIM8 (500–757, 1016–1226, 1427–1655) are not modeled. For the CYRANO trigger RNA, nucleotides 27–52 could be modeled (Supplementary Table 1). Bottom: Domain architecture of ZSWIM8, AGO2, CUL3, ELOB, and ELOC. Numbers indicate residues at domain boundaries. For CUL3, the N-terminal domains of the truncated protein used in the structural elucidation are shown. **b,** Top: Cut-out cryo-EM map showing AGO2 bound to the miR-7–CYRANO duplex, between ZSWIM8 protomers. Middle: Map and model of the miR-7–CYRANO duplex. Bottom: Schematic representation of the miR-7–CYRANO duplex as observed in the density.

### A distinct class of cullin–RING ligases

ZSWIM8 differs from structurally characterized CRL substrate-binding receptors in having a dimeric E3 superdomain, which coordinates ZSWIM8 dimerization, CUL3 ubiquitin ligase assembly, and substrate binding (Extended Data Figure 3a). The dimeric E3 superdomain is a singular interconnected ZSWIM8 unit comprising four regions. First, a zinc-binding SWIM domain (residues 172– 208) is the central organizer of the dimeric E3 superdomain (Extended Data Figure 3a–c). It is essential for target-directed degradation of miR-7 in cells, as measured using a reporter assay that provides a fluorescent readout of cellular miR-7 activity^22^ upon rescue of *ZSWIM8*-knockout cells with a wild-type or variant *ZSWIM8* transgene of interest (Extended Data Figure 3d–g, Supplementary Figure 5, 6, Supplementary Table 3).

Second, the dimerizing D-domain (residues 225–268) comprises helices from each ZSWIM8 protomer that intertwine in a manner analogous to those of a D-domain observed in β-TRCP^63,64^ (Extended Data Figure 3h–j). In ZSWIM8, the D-domain forms an intermolecular knot, rendering dimerization likely irreversible when the adjacent domains fold and thereby prevent unthreading. To test the role of ZSWIM8 dimerization, we designed a monomeric version of ZSWIM8 (ZSWIM8^mono^) that retained CUL3 recruitment and intrinsic E3 ligase activity as detected by autoubiquitylation (Extended Data Figure 3k,l). ZSWIM8^mono^ was defective at CYRANO-directed polyubiquitylation of AGO2–miR-7 *in vitro* and was correspondingly unable to rescue miR-7 degradation in ZSWIM8-deficient cells (Extended Data Figure 3m,n).

Third, an extended region (residues 269–290) emanates from the D-domain and connects the dimeric E3 superdomain to the downstream substrate-binding domain. We term this element the “SWIM belt” because it wraps around and secures several regions, including the BC-box and SWIM domain of the same protomer, the D-domain of the opposite protomer, and part of the N-terminal tail of CUL3 (Extended Data Figure 3a–c).

Fourth, ZSWIM8 displays a unique BC–cullin-box (residues 71–111)^53^ (Extended Data Figure 4a–c) that collaborates with other regions of the dimeric E3 ligase superdomain to partner with CUL3. Compared to the cullin-binding elements found in BC-family CRL substrate-binding receptors, a ZSWIM-family-specific CUL3-box interacts with CUL3 and presents a steric blockade to other cullins (Extended Figure 4d–f). Moreover, ELOC binds CUL3 cullin repeat 1 in a manner resembling how BTB domains bind CUL3 (ref. 65), which is also reminiscent of how ELOC otherwise binds CUL2 (ref. 66) (Extended Data Figure 4c). Finally, the distinctive N-terminal tail of CUL3 embeds into a unique groove consisting of elements from the BC–cullin-box, D-domain, and SWIM belt^67,68^ (Extended Data Figure 3h, 4g). This N-terminal tail anchors CUL3, enabling its rotation around the D-domain as shown by 3D variability analysis (Supplementary Video 2). Removal or mutation of the CUL3 N-terminal tail reduced the affinity of CUL3 for ZSWIM8 and diminished AGO2 polyubiquitylation *in vitro* (Extended Data Figure 4h,i, Supplementary Figure 7). Reciprocally, mutation of ZSWIM8 residues (E91, E94) that contact the CUL3 N-terminal tail diminished AGO2 polyubiquitylation *in vitro* and abolished TDMD in cells (Extended Data Figure 4i,j). Thus, the unique involvement of CUL3 in ZSWIM8-mediated TDMD is achieved by multivalent interactions with ZSWIM8 that select against non-cognate cullins and select for CUL3. Taken together, these results show how the ZSWIM8 dimeric E3 superdomain coordinates the key functions of ZSWIM8 dimerization and CUL3 binding, which are both essential for the activity of the ZSWIM8 CRL in TDMD. Considering that the key CUL3-binding elements are maintained in other ZSWIM-family BC-box proteins (Extended Data Figure 4d), these results define a distinct class of E3 ligases.

### Trigger reshapes AGO2–miRNA

The substrate-binding regions emanating from the ZSWIM8 dimeric E3 superdomain form a clamp that engulfs the AGO2–miR-7–trigger complex. This clamp makes multivalent interactions with the PAZ, N, and MID domains of AGO2 and with the trigger RNA. A key question was the extent to which the AGO2–miRNA–trigger complex bound by ZSWIM8 resembles the previously determined AGO– miRNA–target structures lacking ZSWIM8. In the absence of ZSWIM8, AGO–miRNA–target complexes populate distinct conformations depending on the miRNA–target pairing^14,69–77^ (Figure 3a, Extended Data Figure 5a). Most prominently, the position of the AGO2 PAZ domain reports on the extent of pairing between the miRNA and the target site, with its position progressively shifting as pairing transitions from 3′-supplementary pairing to TDMD pairing (which involves more extended pairing to the miRNA 3′ region) to full pairing^14,72,76^ (Figure 3a, Extended Data Figure 5a, Supplementary Video 3). Transitioning from 3′-supplementary pairing to TDMD pairing also releases the miRNA 3′ end from the PAZ domain, allowing trigger pairing to propagate towards the miRNA 3′ end along a distinct trajectory out of the AGO2 RNA-binding chamber^14,72^ (Figure 3a, Extended Data Figure 5a,b, Supplementary Video 3).

**Figure 3:**
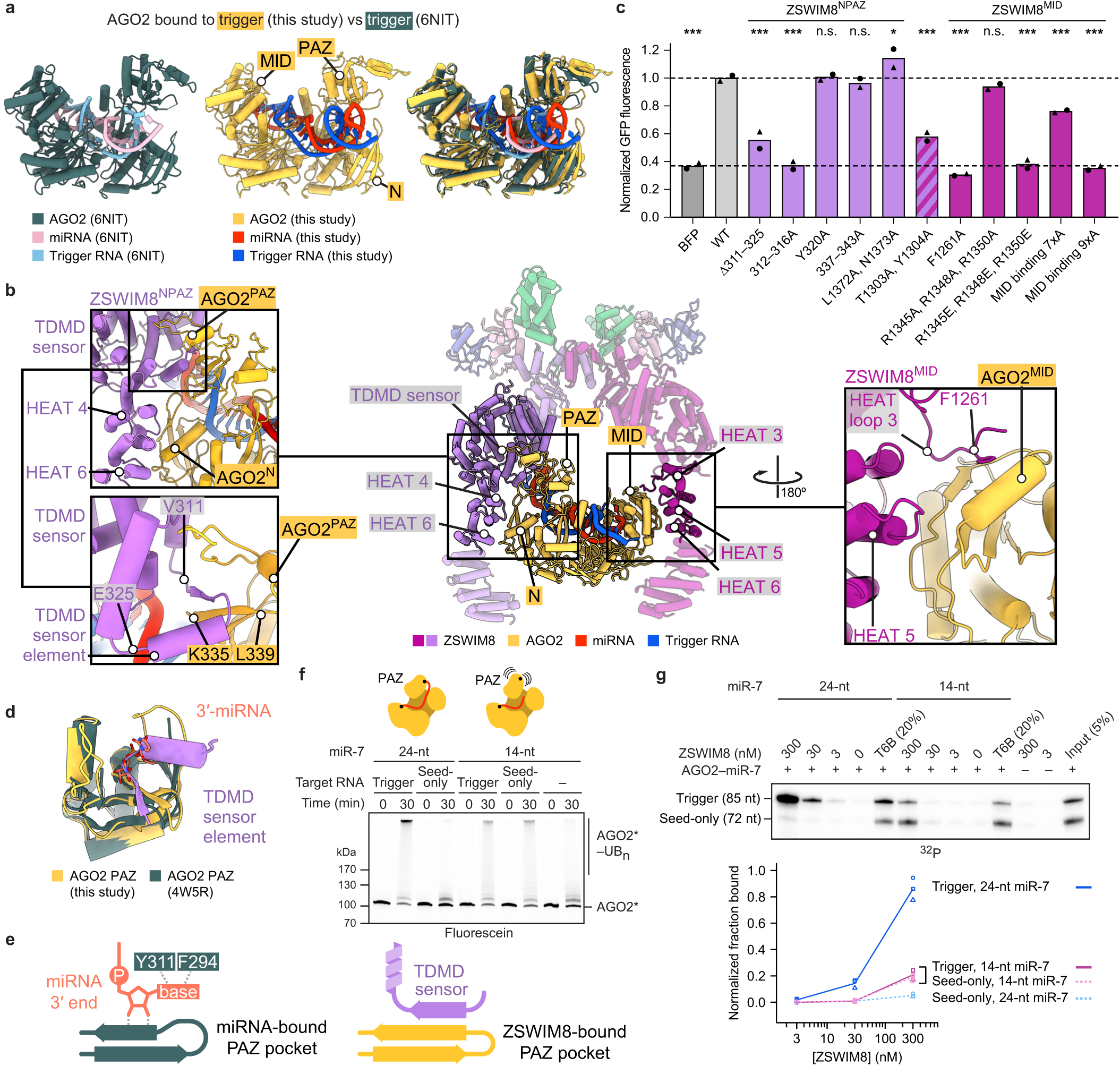
Multivalent interactions between the ZSWIM8 dimeric clamp and AGO2 specify TDMD. **a,** Cartoon representations comparing the structure of trigger-bound AGO2–miRNA in association with ZSWIM8–CUL3 to a previously determined structure of AGO2–miRNA bound to a trigger RNA (but without ZSWIM8–CUL3) (PDB: 6NIT). **b,** Cartoon representation illustrating ZSWIM8–CUL3 bound to AGO2–miR-7–CYRANO, highlighting interactions between ZSWIM8 and AGO2. Middle: Overview of highlighted interactions. Top left: Multivalent interactions between ZSWIM8^NPAZ^ and the AGO2 PAZ and N domains. Bottom left: Close-up of ZSWIM8 TDMD sensor element interacting with the AGO2 PAZ-domain pocket. Starting and ending residues of the TDMD sensor element (V311, E325) are indicated. Right: Multivalent interactions between ZSWIM8^MID^ and the AGO2 MID domain, with the ZSWIM8 F1261 residue indicated. **c,** Importance of ZSWIM8 clamp interactions for CYRANO-directed miR-7 degradation in cells. Plotted are results of the intracellular TDMD assay testing the ability of ZSWIM8 variants to rescue TDMD activity in *ZSWIM8*-knockout cells (Extended Data Figure 3d–g), with a focus on mutations of residues that contact AGO2. Bars for mutant variants are colored by the relevant protomer that uses those residues to make the interaction (purple, ZSWIM8^NPAZ^; magenta, ZSWIM8^MID^; stripes, both). For Δ311–325, residues 311–325 were replaced with (GGGS)_3_; for MID-binding variants, either seven or nine residues were substituted with alanines (MID binding 7xA, MID binding 9xA, respectively) (Supplementary Table 3); for other variants, the indicated residues were substituted with either alanine (A) or glutamate (E). Measurements are plotted as in Extended Data Figure 3f. Significance was measured using an ordinary one-way ANOVA with Dunnett’s multiple comparisons tests to compare the mean of each variant to that of WT (\**P* < 0.05, \*\*\**P* < 0.0001; n.s., not significant). *n* = 2 biological replicates. **d,** Cartoon representation of an isolated PAZ domain illustrating binding of the ZSWIM8 TDMD sensor element to the PAZ-domain pocket, overlaid with binding of a miRNA 3′ terminus to the PAZ-domain pocket, as previously characterized (PDB: 4W5R). **e,** Schematic of mutually exclusive interactions between the PAZ-domain pocket of AGO2 and either the miRNA 3′ end (left) or the ZSWIM8 TDMD sensor element (right). **f,** Effect of miR-7 truncation on AGO2 ubiquitylation. *In vitro* ubiquitylation assays compare polyubiquitylation of AGO2 in complex with either a 24-nt miR-7 or a truncated, 14-nt miR-7, which is too short to occupy the 3′-end binding pocket of the PAZ domain; otherwise, as in Figure 1c. Shown is a representative experiment; *n* = 2 technical replicates. Gels stained with Coomassie blue are shown in Supplementary Figure 1. **g,** Effect of miR-7 truncation on ZSWIM8 binding to AGO2–miR-7–CYRANO. This panel is as in Figure 1f, except the trigger-bound AGO2–miRNA complexes are those of Figure 3f, and measurements corresponding to seed-only-bound complexes are plotted using dashed lines. *n* = 3 technical replicates.

The AGO2–miRNA–trigger conformation bound by ZSWIM8 in our structure resembles the TDMD conformation previously observed for AGO2–miRNA– trigger complexes in the absence of ZSWIM8^14^ (Figure 3a). This resemblance indicates that ZSWIM8 largely recognizes AGO2 in the conformation intrinsically induced by miRNA– trigger pairing. Nonetheless, some differences observed in our E3-bound structure, including a shift of the PAZ domain (Figure 3a), suggest that some features of the AGO conformation are captured upon binding the E3. Additionally, the internal loop of the RNA duplex—which is either partially disordered or fully unpaired in prior structures^14^—is fully resolved in the ZSWIM8 complex and includes an A:U pair and intercalating nucleobases in a configuration that might support the AGO2 conformation recognized by ZSWIM8 (Figure 2b).

Importantly, ZSWIM8 binding appears incompatible with domain conformations observed for AGO2–miRNA complexes not engaged with a TDMD trigger. Superimposing over the MID–PIWI lobe of various AGO2– miRNA complexes (bound to either no target, a seed-only target, a target with 3′-supplementary pairing, or a target with full complementarity to the miRNA) showed that the PAZ and N domains would either clash with or fail to contact ZSWIM8^71,72,76^ (Extended Data Figure 5c). Thus, pairing to the trigger RNA reshapes the AGO2–miRNA complex into a conformation recognized by ZSWIM8.

### Multivalent interactions between ZSWIM8 and AGO2

Following the ZSWIM8 dimeric E3 superdomain is a helix- and loop-rich region (residues 290–915) we term the “TDMD sensor” because it detects most features imparted by the trigger RNA to the AGO2–miR-7 complex. Following this sensor is a stretch of HEAT repeats that also bind the substrate (Figures 2a, 3b, Extended Data Figure 6a,b, Supplementary Video 4). HEAT repeats form curved shapes that undergo accordion-like twisting and turning^78,79^, enabling the two ZSWIM8 protomers to adopt different structures and asymmetrically capture the AGO2 substrate between them (Extended Data Figure 6c, Supplementary Video 5).

In the ZSWIM8^NPAZ^ protomer, the TDMD sensor makes three-way interactions with the miRNA–trigger duplex and the RNA-oriented PAZ and N domains (Figure 3b, Extended Data Figure 6d,e). A loop-helix-turn-helix-loop element (ZSWIM8 residues 309–345) intercalates between a quartet of hairpins in the AGO2 PAZ domain. It contains a “TDMD sensor element” (ZSWIM8 residues 311–325) that is only visible in ZSWIM8^NPAZ^, where it forms a β-sheet with one of the PAZ-domain hairpins (ZSWIM8 residues 311–314, AGO2 residues 335–337) (Supplementary Video 4). Following this β-strand, downstream residues traverse across the interior of the PAZ domain, contact the N domain, then reverse orientation to wrap around the distal edge of the PAZ domain (ZSWIM8 residues 337–343, AGO2 residues 297–303) (Figure 3b). In addition, loops between HEAT repeats 3–4 and 5–6 form a surface that binds the edge of the AGO2 N domain (ZSWIM8 residues 1302–1306 and 1369– 1375, AGO2 residues 116–127 and 81–85, respectively) (Figure 3b, Extended Data Figure 6d).

The opposite protomer, ZSWIM8^MID^, uses the concave interior of HEAT repeats 3–6 to instead recognize the edge of the MID domain (Figure 3b, Extended Data Figure 6d). In addition, an extended ZSWIM8 loop within HEAT repeat 3 anchors the MID domain, primarily through a phenylalanine (F1261) inserted into an exposed hydrophobic cavity of the MID domain. This loop is only observed in the ZSWIM8^MID^ protomer, further illustrating the asymmetric recognition of AGO2 from the two protomers. As the AGO2 MID domain does not undergo substantial structural rearrangement in the transition to the TDMD conformation, we propose that these interactions serve as anchor points to help position the AGO–miRNA– trigger complex in an orientation suitable for the sensor domain of the opposite protomer to selectively recognize trigger-induced conformational changes in the N and PAZ domains, along with the trigger itself.

We then characterized these interactions by introducing mutations at the corresponding interfaces. Due to the unique interactions formed by each ZSWIM8 protomer, mutations were designed with the goal of selectively impacting only one interface at a time. Results from these assays and from assays examining the effects of mutating AGO2, described below (and in Supplementary Table 3), demonstrated the role of the ZSWIM8 dimeric clamp in forming asymmetric, multivalent interactions that recognize TDMD-competent AGO.

#### TDMD sensor interactions

Replacing the TDMD sensor element with a linker (Δ311–325), or mutating residues 312– 316, significantly impaired the ability of ZSWIM8 to rescue miR-7 degradation in ZSWIM8-deficient cells (Figure 3c). This loss of TDMD was attributable to reduced polyubiquitylation, as indicated by biochemical assays (Extended Data Figure 6f).

We also examined the effects of mutating AGO2 PAZ-domain residues recognized by the TDMD sensor element. In a cell-based AGO2 co-IP assay, epitope-tagged AGO2 variants were expressed in wild-type and Δ*Zswim8* mouse embryonic fibroblasts, and the levels of both miR-7 and control miRNAs that co-IPed with each variant were quantified on northern blots (Extended Data Figure 7a). However, results of PAZ mutations were difficult to interpret in this assay, presumably because the mutations also caused defects in PAZ–miRNA association that impaired complex formation or stability in cells, causing reduced co-IP of not only miR-7 but also the control miRNAs—even in the absence of ZSWIM8 (Extended Data Figure 7b,c). Therefore, for such PAZ variants, we turned to our *in vitro* ZSWIM8 co-IP assay, as we were able to generate the recombinant AGO2–miR-7 complexes in cell lysates and purify them with yields sufficient to perform this assay. Replacement of the PAZ loop at AGO2 residues 332–336 resulted in a 12-fold reduction in trigger-dependent ZSWIM8 co-IP (Extended Data Figure 7d,e). Furthermore, replacement of the PAZ-domain loop at AGO2 residues 296– 305, which also contacts the TDMD sensor, also reduced trigger-dependent ZSWIM8 binding, reinforcing the importance of the PAZ domain as one of the binding platforms for ZSWIM8.

#### Anchoring interactions with AGO2 MID

ZSWIM8 residue F1261 docks at the center of the MID-domain interface, and its mutation to alanine reduced AGO2 polyubiquitylation *in vitro* and abrogated TDMD in cells (Figure 3c, Extended Data Figure 6f). Reciprocally, mutation of AGO2 residues (K493, F491) that bind ZSWIM8 F1261 resulted in accumulation of associated miR-7 in cells (Extended Data Figure 7b,c).

#### HEAT repeat interactions

Substitutions at ZSWIM8 HEAT-repeat residues 1303 and 1304 were also detrimental (Figure 3c, Extended Data Figure 6f). However, because these residues appear to contact AGO2 in both protomers, interacting with the N and MID domains in the cases of ZSWIM8^NPAZ^ and ZSWIM8^MID^, respectively, the contributions of these potentially bifunctional residues could not be assigned to a single protomer.

AGO2 residues that interact with ZSWIM8 are conserved across AGO paralogs in humans and across AGO2 homologs in bilaterian species—especially those thought to be TDMD-sensitive (Supplementary Figure 10, 11). Likewise, ZSWIM8 residues that interact with AGO2 are conserved across bilaterian species (Supplementary Figure 12), suggesting a general recognition mode of TDMD substrates by ZSWIM8.

### ZSWIM8 recognizes unoccupied PAZ-domain pocket

The ZSWIM8 TDMD sensor element interacts with the AGO2 PAZ domain near the pocket that usually binds the miRNA 3′ terminus^80^ (Figure 3d,e), suggesting that these are mutually exclusive interactions. Upon extensive pairing between the trigger and the 3′ region of the miRNA, the miRNA 3′ terminus is extracted from the PAZ pocket, which would allow the TDMD sensor element to access its binding site (Figure 3d,e). In this way, the ZSWIM8 TDMD sensor element might specifically recognize not only the position of the PAZ domain but also an unoccupied PAZ pocket as a feature of trigger-bound AGO–miRNA complexes. To test this model, we performed binding assays using AGO2–miR-7 variants designed to display a more constitutively accessible PAZ pocket, as described below.

#### Mutation ofi AGO2 PAZ pocket

With the goal of impairing miRNA 3′-end binding to the PAZ pocket, PAZ residues that interact with the terminal miRNA nucleotide at the nucleobase (F294A, Y311A) were mutated^14,75^ (Figure 3e). These substitutions caused increased binding to the seed-only-bound AGO2–miR-7 (Extended Data Figure 7d,e), consistent with an unoccupied PAZ pocket comprising a feature selectively recognized by ZSWIM8.

#### miRNA truncation

We further tested sufficiency of an unoccupied PAZ pocket for mediating ZSWIM8 binding using another approach. AGO2 was loaded with a shortened miR-7, truncated after nucleotide 14, which was designed to be long enough to achieve seed-based target binding but too short to engage with the PAZ domain^81,82^. This truncation renders the 3′-end binding pocket within PAZ unoccupied, regardless of whether the miRNA is bound to CYRANO or to its seed-only counterpart. Indeed, with truncated miR-7, little difference was observed between the trigger and seed-only versions of CYRANO for both ZSWIM8 binding and AGO2 polyubiquitylation (Figure 3f,g). Nonetheless, trigger-induced binding and polyubiquitylation were both reduced with truncated miR-7 compared to full-length miR-7. Moreover, the residual polyubiquitylation required the presence of a target RNA, even though the truncated guide lacked the nucleotides required to form extensive 3′ pairing. These results suggested that an unoccupied PAZ pocket is important but not wholly sufficient to mediate binding to the ZSWIM8 CRL, and that the purpose of extensive 3′ pairing to the trigger is not simply to vacate the PAZ pocket.

The observation of some binding and polyubiquitylation of AGO2 complexes containing truncated miRNAs suggested that similar complexes containing naturally truncated miRNAs (presumably resulting from extensive 3′-exonucleolytic trimming)^83^ might also be susceptible to ZSWIM8-dependent degradation in cells. To test this, we performed small-RNA sequencing of AGO-associated RNAs in *Zswim8* wild-type and knockout cells to examine whether cells containing ZSWIM8 accumulate fewer miRNAs shorter than 19 nt. A statistically significant, albeit weak, signal for ZSWIM8-dependent reduction of such shortened miRNAs was observed, with a more substantial effect in Drosophila cells than in mammalian cells (Extended Data Figure 8, Supplementary Figure 13). These results suggested that in addition to its role in TDMD, ZSWIM8 might act more broadly to destabilize AGO–miRNA complexes that contain extensively trimmed miRNAs.

### Flanking trigger RNA embraces ZSWIM8

Given the importance of RNA flanking the trigger site in directing AGO2–miR-7 binding to ZSWIM8 (Figure 1f, Extended Data Figure 2), we considered the trajectory of the miRNA–trigger duplex and how it might position the RNA flanking the trigger site. The internal loop of the miRNA– trigger duplex enables a bend between the seed and distal RNA helices^14^. This bend causes the trajectory of the distal RNA helix to differ from that of other AGO–miRNA–target conformations (Figure 4a), facilitating contacts between the end of the distal helix and ZSWIM8^NPAZ^. These interactions were observed at high resolution for a loop within the TDMD sensor domain that we term RNA-binding element 1 (RBE1, residues 395–408). RBE1 wedges between the TDMD sensor element, the miRNA–trigger duplex, and the HEAT repeat subdomain. Together with the AGO2 PAZ domain, RBE1 clasps the final turn of miR-7 as presented in the context of the AGO2-bound miRNA–trigger duplex (Figure 4b). Accordingly, a ZSWIM8 variant harboring charge-reversal substitutions in RBE1 had reduced ability to rescue TDMD in ZSWIM8-deficient cells and reduced polyubiquitylation activity *in vitro* (Figure 4c, Extended Data Figure 9a).

**Figure 4:**
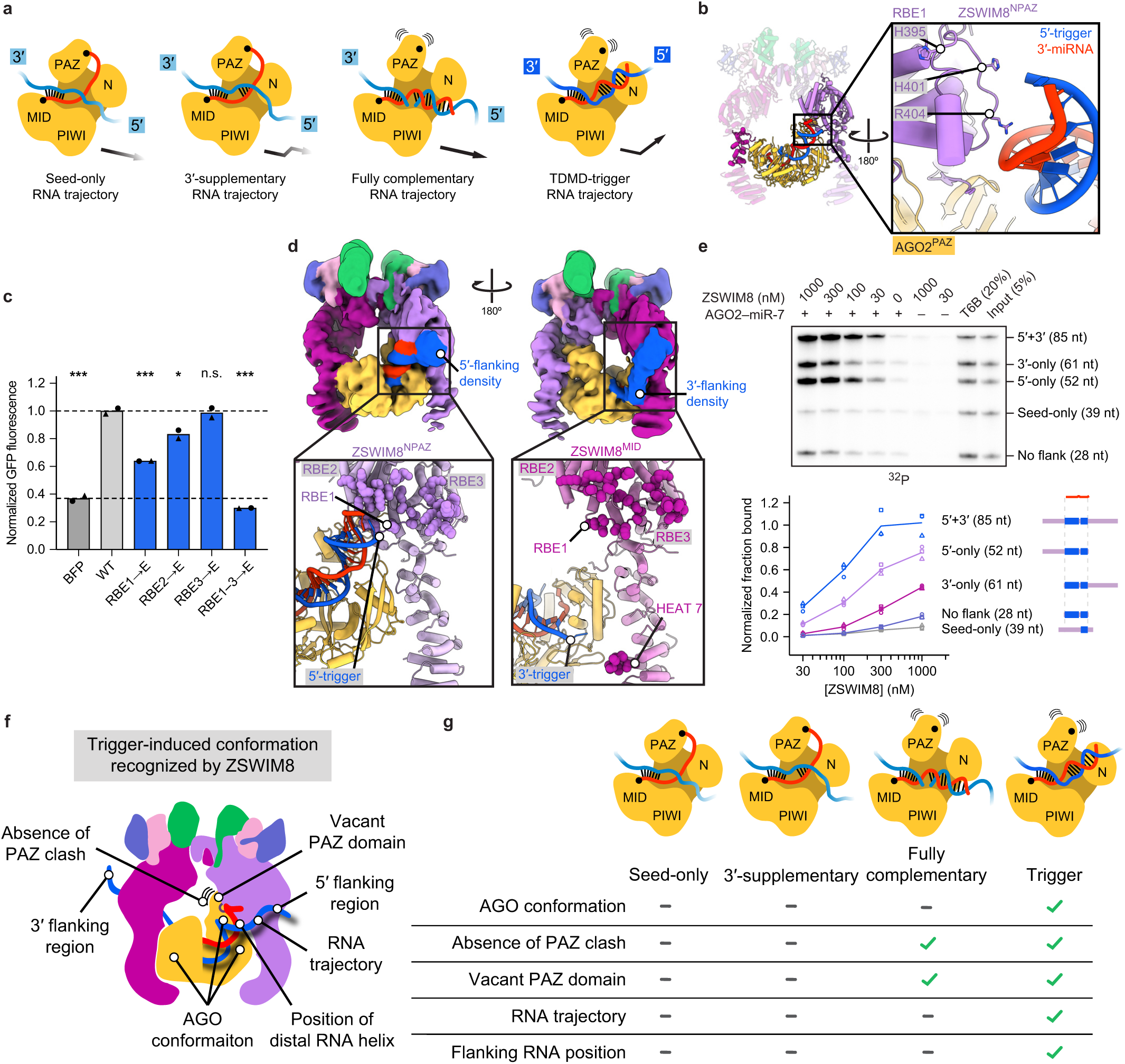
Flanking trigger RNA embraces ZSWIM8. **a,** Comparison of trajectories of target-RNA 5′ regions as they emerge from AGO2–miRNA– target structures with either seed-only pairing, 3′-supplementary pairing, fully complementary pairing, or trigger pairing (based on PDBs: 4W5R, 6N4O, and 9CMP, and this study, respectively). Black arrows represent a simplified RNA trajectory of each target RNA, with opacity roughly indicating the degree of conformational constraint. **b,** Cartoon representation illustrating ZSWIM8^NPAZ^ interactions with the 3′ region of the miRNA and the 5′ region of the trigger RNA. RBE1 and its putative RNA-interacting residues are shown. **c,** Importance of ZSWIM8 RBEs for CYRANO-directed miR-7 degradation in cells. Plotted are results of the intracellular TDMD assay testing the ability of ZSWIM8 variants with mutations in RBEs to rescue ZSWIM8 activity in cells. Otherwise, this panel is as in Figure 3c. Significance was measured using an ordinary one-way ANOVA with Dunnett’s multiple comparisons tests to compare the mean of each variant to that of WT (\**P* < 0.05, \*\*\**P* < 0.0001; n.s., not significant). *n* = 2 biological replicates. **d,** Cryo-EM evidence for RNA in the vicinity of the RBEs of both ZSWIM8^NPAZ^ and ZSWIM8^MID^. Top: Cryo-EM map low-pass filtered to 10 Å. Density corresponding to the modeled complex is colored as in Fig 2. Additional densities assigned as flanking RNA are highlighted in blue. Top-left panel shows additional 5′-trigger density interacting with ZSWIM8^NPAZ^. Bottom-left panel shows a cartoon representation with residues of RBEs 1–3 highlighted as purple spheres. Top-right panel shows additional 3′-trigger density interacting with ZSWIM8^MID^. Bottom-right panel shows a cartoon representation with residues of RBEs 1–3 highlighted as magenta spheres. Colors are as in Figure 2. **e,** Contributions of RNA flanking the 5′ and 3′ ends of the trigger site to ZSWIM8 binding to AGO2–miR-7–CYRANO. Shown are results of the *in vitro* co-IP assay using radiolabeled fragments of the CYRANO trigger RNA with the indicated site-flanking regions and lengths; otherwise, as in Figure 1f. *n* = 3 technical replicates. **f,** Model figure depicting structural changes in the AGO–miRNA complex induced by trigger binding, and the interactions leveraged by ZSWIM8 to detect trigger-bound AGO–miRNA. **g,** Extent to which various AGO–miRNA–target complexes contain the key features for ZSWIM8 recognition.

Although not visible at high resolution, additional density emanating from the 5′ and 3′ ends of the trigger site and embracing both ZSWIM8 protomers was visible at low resolution (Figure 4d, Extended Data Figure 9b). Conformational heterogeneity, as observed in 3D variability analysis of the cryo-EM data, explained why RNA flanking the trigger site was not visible at high resolution (Supplementary Video 6). Nonetheless, the low-resolution maps showed the 5′ flanking region of the trigger RNA surrounding RBE1 as well as two additional positively charged loops we term RBE2 and RBE3 (residues 460–470 and 803–823, respectively) from ZSWIM8^NPAZ^ (Figure 4d, Extended Data Figure 9c). Low-resolution maps also suggested a path for the 3′ flanking region of the trigger RNA; weak density was visible exiting AGO2 adjacent to a C-terminal ZSWIM8^MID^ HEAT repeat and extending to RBEs 1–3 of this protomer (Figure 4d, Supplementary Video 7). Thus, the two trigger regions that flank the miRNA-binding site interact with the two ZSWIM8 protomers to form a cross-brace securing the AGO2–miRNA complex inside the E3 ligase clamp. Note that in contrast to the TDMD sensor, which appears to function only in the context of the ZSWIM8^NPAZ^ protomer, the RBEs appear to function in the context of both the ZSWIM8^NPAZ^ and the ZSWIM8^MID^ protomers, interacting with the 5′ and 3′ flanking regions, respectively. We tested the effects of altering the RBEs and trigger-RNA flanking regions, as described below.

#### RNA binding elements

Although charge-reversal mutations within RBE2 and RBE3 did not individually impact rescue of miR-7 TDMD in ZSWIM8-deficient cells, the combined mutation of all three RBEs exacerbated the defect caused by mutating RBE1, leading to severe loss of TDMD in cells and of polyubiquitylation activity *in vitro* (Figure 4c, Extended Data Figure 9a). Thus, the interactions between the trigger RNA and the ZSWIM8 RBEs revealed in our structure can rationalize the biochemical roles of flanking trigger RNA (Figure 1f, Extended Data Figure 2).

#### Trigger filanking regions

The structure suggested that 25 nt of trigger RNA flanking the miRNA-binding site would be sufficient for both the 5′ and 3′ flanks to access the three RBEs on ZSWIM8^NPAZ^ and ZSWIM8^MID^, respectively. Indeed, lengthening the flanking sequences beyond 25 nt on either side had little effect on the interaction with ZSWIM8 (Extended Data Figure 9d), whereas eliminating the RNA immediately flanking either end of the trigger site reduced ZSWIM8 binding and AGO2 polyubiquitylation (Figure 4e, Extended Data Figure 9e). Deleting the 5′ flank was three-fold more consequential to binding compared to deleting its 3′ counterpart (Figure 4e). This directional hierarchy was maintained with two sets of scrambled flanking sequences, which indicated that the contributions of the 5′ and 3′ flanks did not require sequence-specific contacts (Extended Data Figure 9f,g). The stronger effect of the 5′ flanking sequence depended on pairing to the 3′ region of the miRNA (Extended Data Figure 9h). These results supported the idea that trigger pairing specifies a trajectory of the 5′ flank of the trigger RNA, which favors interaction with the ZSWIM8^NPAZ^ RBEs.

To further examine contributions of the trigger-specified RNA trajectory, we performed binding and ubiquitylation assays with AGO2–miR-7 paired to a fully complementary target. As with TDMD pairing, fully complementary pairing extracts the miRNA 3′ end from its pocket in the PAZ domain^75–77^. However, in contrast to TDMD pairing, which induces a 40° bend in the duplex, the fully paired duplex adopts a relatively straight conformation and thus exits AGO2 along a different trajectory (Figure 4a). Assays performed using a fully complementary target RNA and an AGO2 active-site variant (D669A, to prevent slicing of fully complementary target RNA)^84^ showed weak binding to ZSWIM8 and polyubiquitylation activity intermediate between that of the seed-only and trigger RNAs (Extended Data Figure 10). Therefore, despite presenting an unoccupied PAZ pocket, AGO–miRNA bound to a fully complementary target is a suboptimal partner for ZSWIM8, which we attribute to its misaligned RNA. Together, these results reinforce the concept that miRNA 3′-end release and trigger-specific RNA trajectory are emergent features of miRNA–trigger pairing, which drive AGO recognition by ZSWIM8.

## Discussion

Our results revealed how the exquisite selectivity of TDMD is achieved through numerous features orchestrated by the trigger RNA. In particular, pairing of the trigger RNA to an AGO2–miRNA complex induces 1) a structural remodeling of the AGO2 protein that arranges the MID, N, and PAZ domains for simultaneous engagement by two interlocked ZSWIM8 protomers; 2) displacement of the miRNA 3′ end from its binding pocket in the AGO2 PAZ domain, which exposes the binding pocket for recognition by the ZSWIM8 TDMD sensor domain; and 3) a unique trajectory of the miRNA–trigger duplex, which favors direct contacts to the ZSWIM8 sensor domain and guides flanking regions of the trigger to engage with positively charged ZSWIM8 elements (Figure 4f,g). This multi-factorial selectivity ensures that miRNA degradation is tightly regulated such that AGO– miRNA complexes paired to non-trigger transcripts remain active as needed for biological regulation. We suspect that the reduced recognition of fully complementary targets by ZSWIM8 also helps explain the high efficacy and long duration of siRNA therapies^85^.

The myriad trigger-RNA-induced interactions specifying substrate recognition by the ZSWIM8 E3 ligase do not correspond to a traditional degron motif^26–28^. Rather than interacting with either a linear peptide or a single extended surface, ZSWIM8 uses its dimeric architecture to interact with two extended surfaces of the substrate, and with the RNA. As such, the structure of a ZSWIM8–CUL3-bound AGO2–miR-7–CYRANO complex illuminated a two-RNA-factor authentication mechanism determining E3 ligase substrate specificity. The effects of matching two RNA factors—the miRNA and the trigger RNA—propagate allosterically across both RNAs and the AGO2 protein to drive binding to ZSWIM8. These conformational changes are likely conserved across many miRNA–trigger pairs, converging on a general trigger-bound AGO–miRNA conformation distinct from any other AGO–miRNA conformation. Although the possibility that some triggers might use sequence-specific factors or interactions to compensate for suboptimal miRNA–trigger pairing cannot be excluded^19,62^, this generalized binding mode is supported by several observations. These include the charge-driven, sequence-independent interaction of ZSWIM8 with the trigger RNA within the complex that we examined, and the high sequence conservation specifically among TDMD-sensitive AGO homologs (Supplementary Figure 10, 11). Thus, overall, our work provides a structural and mechanistic framework for how RNA–RNA base pairing can induce generalizable ubiquitin-dependent protein degradation to ultimately induce degradation of a particular RNA.

## Acknowledgements

We thank the Max-Planck Institute of Biochemistry Protein Production Facility (RRID:SCR_025741) for protein expression, the Biochemistry Core Facility (RRID:SCR_025743) for peptide synthesis and access to BLI instruments, and the Cryo-EM facility (RRID:SCR_025744) with Daniel Bollschweiler and Tillman Schäfer for their help with cryo-EM analysis. We thank the Whitehead Institute Flow Cytometry Core for assistance with flow cytometry, the Whitehead Institute Genome Technology Core for high-throughput sequencing, and the Whitehead Bioinformatics and Research Computing Core for assistance with statistical analyses. We thank Joshua Mendell for sharing cell lines. We thank Jonathan Weissman, Kacper Rogala, and Anders Hansen for sharing plasmids. This study was supported by grants from the European Union (ERC, UPSmeetMet, 101098161), Deutsche Forschungsgemeinschaft (DFG, German Research Foundation), and the NIH (GM118135). J.F. was supported by a Postdoctoral Fellowship of the Peter & Traudl Engelhorn Stiftung. E.S. was supported by the National Science Foundation Graduate Research Fellowship Program. P.Y.W. was supported by the MIT Office of Graduate Education Fellowship. D.H.L. was an HHMI fellow of the Damon Runyon Cancer Research Foundation (DRG-2345-18). D.P.B. is an investigator of the Howard Hughes Medical Institute. We thank Josef Kellerman and Filiz Çivril for their support. We thank Leo Kiss, J. Rajan Prabu, Samuel Maiwald, Joanna Liwocha, Lukas Henneberg, Charlie Shi, Bradley Wierbowski, Lara Elcavage, Michelle Frank, Arash Latifkar, and other members of the Schulman and Bartel labs for fruitful discussions. This preprint was typeset according to @chrelli (https://github.com/chrelli/bioRxiv-word-template).

## Author contributions

J.F. performed *in vitro* ubiquitylation assays. S.V.G. performed insect cell expression. E.S. and P.Y.W. performed *in vitro* co-IP assays. E.S. and D.H.L. performed cellular assays testing ZSWIM8 and AGO2. J.F. prepared, collected, processed, and performed in-depth analyses of the cryo-EM structure. E.S., P.Y.W., D.H.L., B.A.S., and D.P.B. also analyzed the cryo-EM structure. L.W.B. performed sRNA sequencing and associated analyses. J.F., E.S., B.A.S., and D.P.B. prepared the manuscript with input from all other authors. The study was supervised by B.A.S. and D.P.B.

---

**Extended Data Figure 1.**
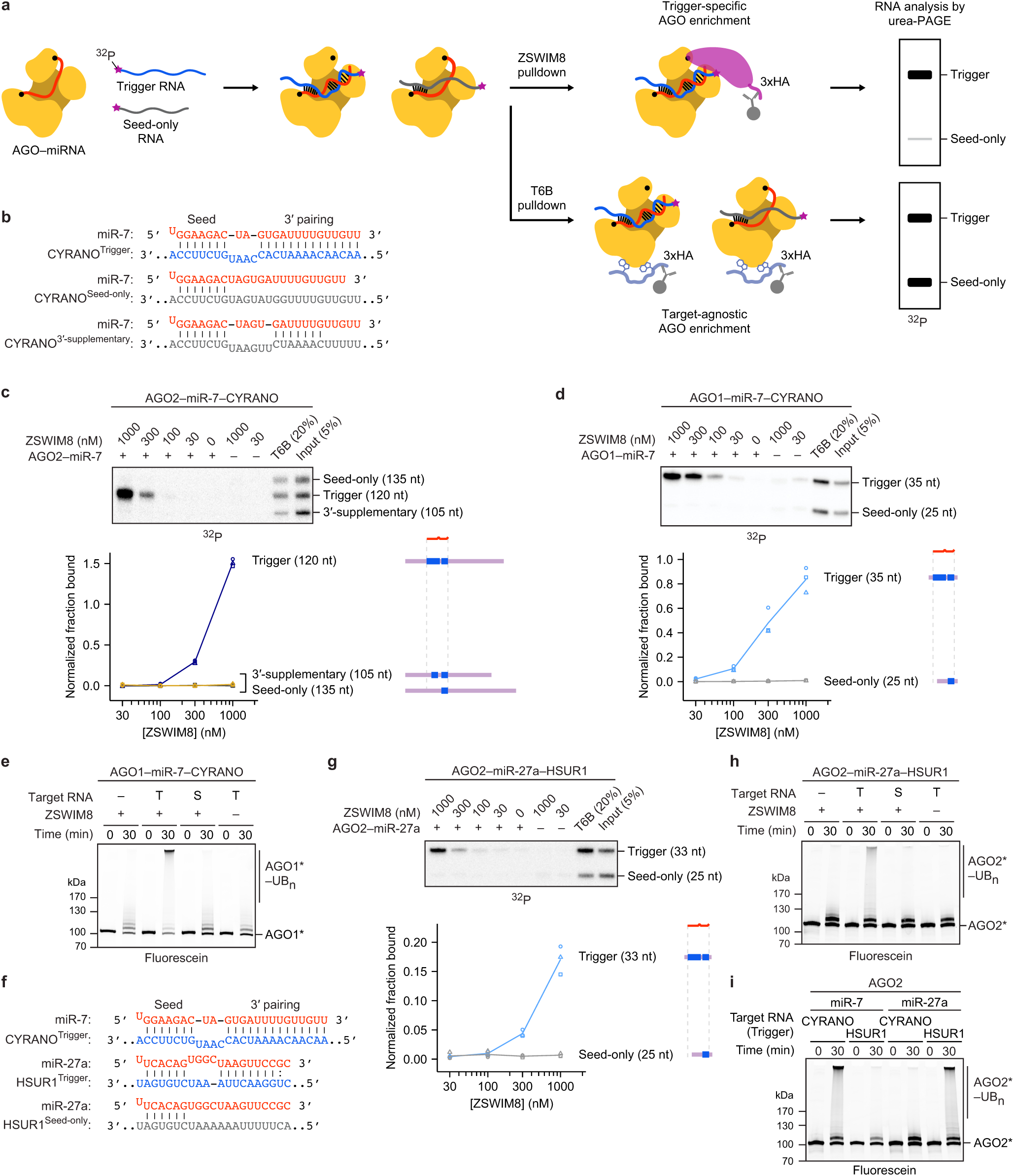
ZSWIM8 binds to AGO–miRNA–trigger complexes to direct the polyubiquitylation of AGO. **a,** Schematic of the *in vitro* co-IP assay for detecting binding of ZSWIM8 to AGO–miRNA–target complexes. AGO–miRNA complexes are incubated with a mixture of 5′-radiolabeled target RNAs of different lengths. These RNAs are in large excess over the *K*_D_ values observed between AGO–miRNA and target RNA, and the AGO–miRNA complex is in slight excess over the total concentration of the target RNAs. The target RNAs are designed to differ with respect to their pairing to the miRNA, such as seed-only or trigger pairing (shown), or with respect to the amount of RNA flanking the site (not shown). The AGO–miRNA–target ternary complexes are then incubated with an excess of either 3xHA-tagged ZSWIM8 or a 3xHA-tagged T6B peptide (derived from TNRC6B). Complexes associating with ZSWIM8 or T6B are enriched by 3xHA IP. Following washes, IPs are eluted and the co-IPed target RNAs are resolved on a denaturing gel. Radiolabeled species are visualized by phosphor imaging. The T6B IP is designed to detect any differences in AGO–miRNA binding to different target RNAs, because T6B is expected to bind AGO–miRNA complexes irrespective of their bound target RNA. Thus, for quantification, signal from each band is background-subtracted using the 0 nM ZSWIM8 sample and then normalized to that of the T6B sample. **b,** Diagrams showing pairing between miR-7 and the miRNA-binding regions of CYRANO^Trigger^, CYRANO^Seed-only^, and CYRANO^3′-supplementary^ target RNAs with their cognate miRNAs used for *in vitro* co-IP assays. Vertical lines represent W–C–F base pairing. **c,** The relative effects of trigger pairing, 3′-supplementary pairing, and seed-only pairing on ZSWIM8 binding to AGO2–miR-7 complexes. Top: Shown are results from the *in vitro* co-IP assay with ZSWIM8 and AGO2–miR-7 mixed with radiolabeled target RNAs derived from the CYRANO trigger containing either trigger, seed-only, or 3′-supplementary pairing to miR-7 (Supplementary Table 1). This panel is as in Figure 1f, except 10 nM of AGO2–miR-7 and ∼1 nM of target RNAs were used (instead of 0.75 nM of AGO2–miR-7 and ∼0.25 nM of target RNAs), and 10 µg/mL of heparin was included (instead of 1 µg/mL of heparin). Bottom: Quantification of radiolabeled co-IPed RNA in the assay shown above. The lane-profile tool was used to better quantify the weak signal representing binding to seed-only and 3′-supplementary targets; otherwise, as in Figure 1f. *n* = 3 technical replicates. This quantification revealed a 100-fold preference for co-IP of target RNAs with trigger pairing over 3′-supplementary pairing, and a larger preference for trigger pairing over seed-only pairing, although this larger preference was difficult to quantify due to the very low signal for co-IP of the seed-only target. **d,** Biochemical reconstitution of selective ZSWIM8 binding to AGO1–miR-7–CYRANO^Trigger^ complexes. This panel is as in Figure 1e, except human AGO2 was replaced by human AGO1. In addition, 6 nM of AGO1–miR-7 and ∼1 nM of target RNAs were used instead of 0.75 nM of AGO2–miR-7 and ∼0.25 nM of target RNAs. *n* = 3 technical replicates. **e,** Biochemical reconstitution of trigger- and ZSWIM8-dependent ubiquitylation of AGO1 using the miR-7–CYRANO miRNA–trigger pair. This panel is as in Figure 1c, except fluorescent AGO2 was replaced by fluorescent AGO1. Trigger and seed-only variants of CYRANO are indicated by T and S, respectively. Shown is a representative experiment; *n* = 2 technical replicates. Gels stained with Coomassie blue are shown in Supplementary Figure 1. **f,** Diagrams showing pairing of miRNA-binding regions of CYRANO^Trigger^, HSUR1^Trigger^, and HSUR1^Seed-only^ target RNAs with their cognate miRNAs used for *in vitro* ubiquitylation and co-IP assays. Dots represent G:U wobble; otherwise as in **b**. **g,** Biochemical reconstitution of selective ZSWIM8 binding to AGO2–miR-27a–HSUR1^Trigger^ complexes. This panel is as in Figure 1e, except miR-7 was replaced by miR-27a, and CYRANO was replaced by HSUR1 (panel **f**). In addition, 10 nM of AGO2–miR-7 and ∼1 nM of target RNAs were used instead of 0.75 nM of AGO2–miR-7 and ∼0.25 nM of target RNAs. *n* = 3 technical replicates. **h,** Biochemical reconstitution of trigger- and ZSWIM8-dependent ubiquitylation of AGO2 using the miR-27a–HSUR1 miRNA–trigger pair. This panel is as in Figure 1c, except miR-7 was replaced by miR-27a, and CYRANO was replaced by HSUR1 (panel **f**). Trigger and seed-only variants of HSUR1 are indicated by T and S, respectively. Shown is a representative experiment; *n* = 2 technical replicates. Gels stained with Coomassie blue are shown in Supplementary Figure 1. **i,** Requirement for cognate miRNA–trigger pairing for AGO2 ubiquitylation. Shown is *in vitro* ubiquitylation of AGO2 comparing activity with different combinations of miRNA–trigger pairs using miRNAs miR-7 and miR-27a, and trigger RNAs CYRANO and HSUR1. Otherwise, this panel is as in Figure 1c. CYRANO trigger (120 nt) and HSUR1 trigger (144 nt) were used as target RNAs (Supplementary Table 1). Shown is a representative experiment; *n* = 2 technical replicates. Gels stained with Coomassie blue are shown in Supplementary Figure 1.

**Extended Data Figure 2.**
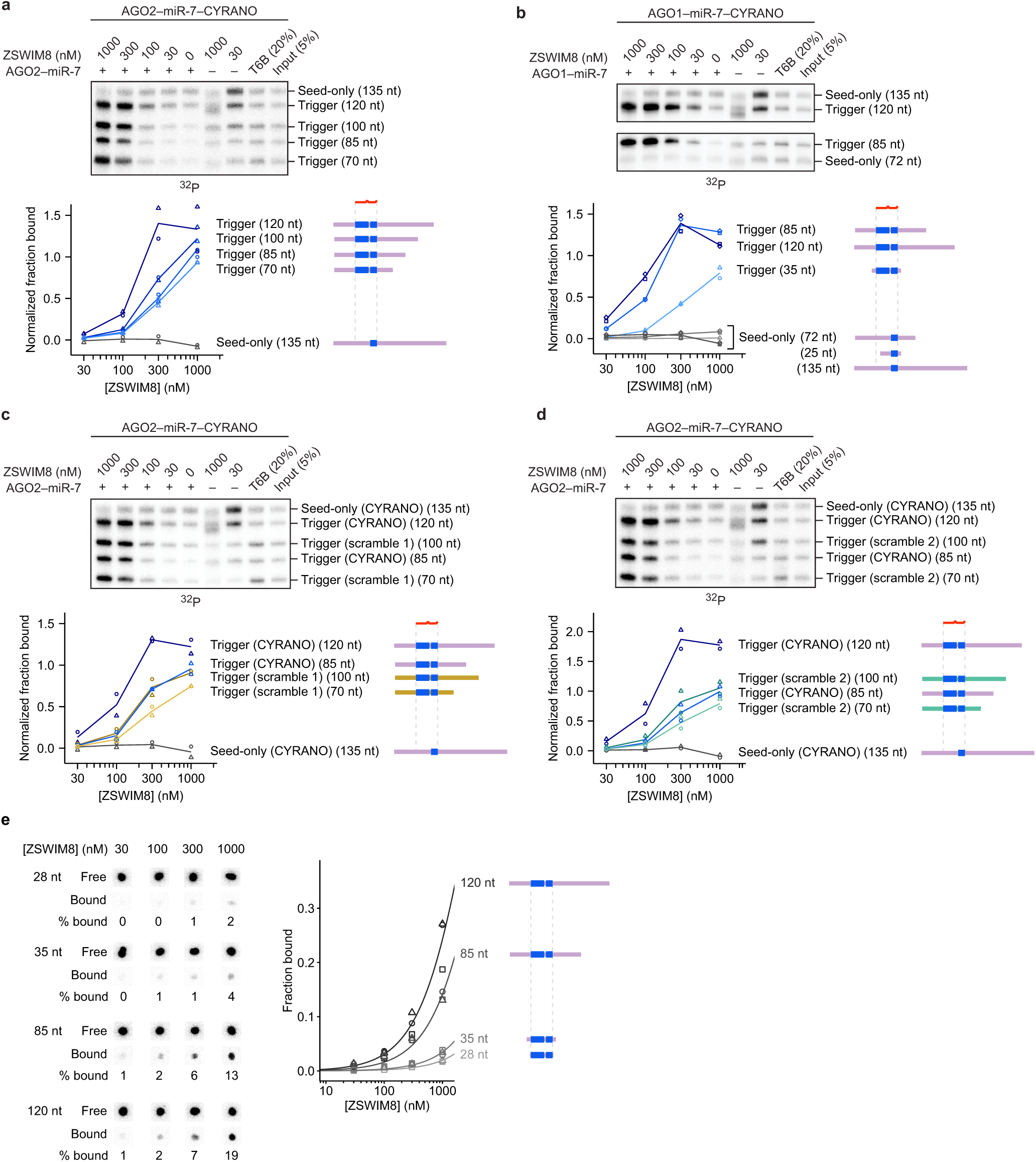
RNA flanking the trigger site contributes to ZSWIM8 affinity. **a,** Effect of finer increments of trigger RNA flanking the 3′ end of the miRNA-binding site on ZSWIM8 binding to AGO2–miR-7–CYRANO. Shown is an *in vitro* co-IP assay like that in Figure 1e, except target RNAs containing trigger pairing had variable amounts CYRANO sequence flanking the 3′ end of the miRNA-binding site. In addition, 10 nM of AGO2– miR-7 and ∼1 nM of target RNAs were used instead of 0.75 nM of AGO2–miR-7 and ∼0.25 nM of target RNAs. *n* = 2 technical replicates. **b,** Effect of trigger RNA flanking the miRNA-binding site on ZSWIM8 binding to AGO1–miR-7–CYRANO. Shown is an *in vitro* co-IP assay like that in **a**, except that 6 nM of AGO1–miR-7 was used instead of 10 nM of AGO2–miR-7. *n* = 2 technical replicates. Results from Extended Data Figure 1d are replotted for comparison. **c,** Sequence specificity of the contribution of trigger flanking sequences to ZSWIM8 binding to AGO2–miR-7–CYRANO. Shown is an *in vitro* co-IP assay like that in **a**, except the flanking sequences were either the native CYRANO sequence, or a scrambled sequence maintaining the nucleotide composition of the original CYRANO sequence (scramble 1). *n* = 2 technical replicates. **d,** Sequence specificity of the contribution of trigger flanking sequences to ZSWIM8 binding to AGO2–miR-7–CYRANO. This panel is as in **c**, except a different scramble combination was used (scramble 2). *n* = 2 technical replicates. **e,** Intrinsic ZSWIM8 affinity for RNA alone. Left: Measurements of RNA of different lengths binding to ZSWIM8 in the absence of AGO–miRNA, as detected by filter binding. Filters were stacked such that the binding reaction passed first through the nitrocellulose filter (which binds proteins and protein-associated RNA) and then through the nylon filter (which binds free RNA). Shown are phosphorimager scans of the filters used to determine the fraction of RNA bound for each concentration of ZSWIM8. For each RNA, the top row shows the amount of radiolabeled RNA designated as free RNA because it passed through the nitrocellulose filter and bound to the nylon filter. The bottom row shows the amount of radiolabeled RNA designated as bound to ZSWIM8 because it was retained with ZSWIM8 on the nitrocellulose filter. Right: Fraction of RNA bound as a function of ZSWIM8 concentration, as detected by filter binding on the left. Spot intensities were background-subtracted, and the bound RNA signal was normalized to the sum of the free and bound RNA signal. The symbols show data from independent measurements. *n* = 3 technical replicates. Curves were fit to a standard fraction-bound curve for visualization, with the plateau constrained to 1.

**Extended Data Figure 3.**
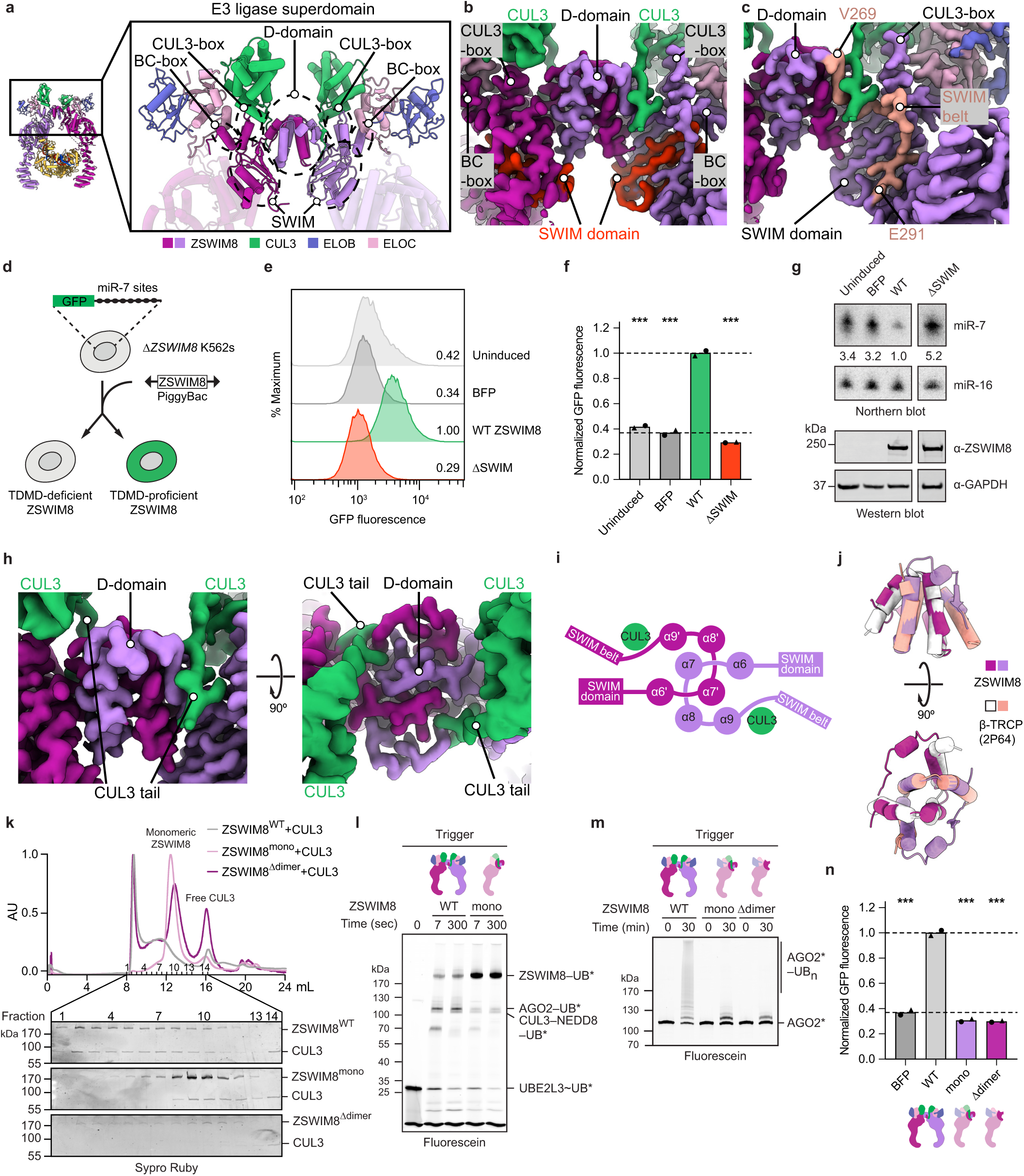
ZSWIM8 contains a dimeric E3 superdomain. **a,** Cartoon representation showing the ZSWIM8 dimeric E3 superdomain and the individual domains associated with it. The E3 superdomain consists of the D-domain, CUL3-box, BC-box, and SWIM domain. **b,** Density showing the SWIM domain and its organization of the E3 superdomain. The SWIM domain is highlighted in red. Density was derived from the composite map. **c,** Density showing the SWIM belt, which connects the D-domain and SWIM domain to the TDMD sensor domain. Density showing the SWIM belt is highlighted in salmon. Density was derived from the composite map. **d,** Schematic of the intracellular TDMD reporter assay. The assay is performed using a clonal *ZSWIM8*-knockout K562 cell line that expresses a GFP reporter containing multiple copies of a miR-7 slicing site^22^. Lack of TDMD in these cells results in elevated levels of endogenous miR-7, which leads to increased repression of *GFP* reporter mRNA, causing reduced GFP fluorescence. To assay the activity of a ZSWIM8 variant, it is introduced using the PiggyBac system and expressed under a doxycycline-inducible promoter, and then its effect on GFP levels is measured by flow cytometry. ZSWIM8 variants proficient at TDMD rescue miR-7 degradation, resulting in increased GFP levels, whereas ZSWIM8 variants deficient in TDMD fail to decrease miR-7 levels, resulting in a lack of increase in GFP levels. **e,** Validation of the intracellular TDMD reporter assay. GFP levels in cells containing wild-type (WT) *ZSWIM8* without doxycycline induction (uninduced, light gray) were determined, as were those in cells expressing BFP (dark gray), WT ZSWIM8 (green), and a variant of ZSWIM8 with alanine substitutions in the SWIM domain (red). To the right of each histogram are median GFP values, each normalized to that of WT ZSWIM8. ZSWIM8 expression levels were detected by western blotting (Supplementary Figure 6). **f,** Results of two biological replicates of the intracellular TDMD reporter assay in **e**. Each point represents the median GFP fluorescence, and each bar height represents the mean of these measurements from two independently derived PiggyBac lines. The top dashed line is drawn at the normalized mean GFP level (1.00) in cells expressing WT ZSWIM8, and the bottom dashed line is drawn at the normalized mean GFP level (0.37) in cells expressing BFP. Significance was measured using an ordinary one-way ANOVA with Dunnett’s multiple comparisons tests to compare the mean of each variant to that of WT (\*\*\**P* < 0.0001). **g,** Levels of miR-7 and ZSWIM8 in reporter cells used to generate the measurements in **e**. For each cell population, northern blots measured the levels of miR-7 and miR-16 (a control miRNA), and western blots detected expression of ZSWIM8 protein. Numbers show the ratio of miR-7:miR-16, normalized to that of cells expressing WT ZSWIM8. **h,** Density showing the ZSWIM8 D-domain and the associated CUL3 N-terminal tail. Density was derived from the composite map. **i,** Cartoon representation illustrating the knot-like assembly of the ZSWIM8 D-domain, along with the associated CUL3 N-terminal tail. Numbers indicate the order of alpha helices as counted from the ZSWIM8 N terminus. **j,** Superimposition of the ZSWIM8 D-domain with the D-domain of β-TrCP (PDB: 2P64). ZSWIM8 residues 220–270 were used for alignment. The D-domains align with a root-mean-square deviation of 1.1 Å. Remaining ZSWIM8 residues are not shown. **k,** Size-exclusion chromatograms of ZSWIM8–CUL3, ZSWIM8^mono^–CUL3, and ZSWIM8^Δdimer^– CUL3 complexes (top) and corresponding SDS-PAGE analysis (bottom). ZSWIM8^mono^ contained a second copy of the D-domain transplanted into its D-domain, creating a constitutively monomeric ZSWIM8 with a folded D-domain. ZSWIM8^Δdimer^ contained a deletion of its D-domain, which was replaced by a G/S-rich linker. Both ZSWIM8^mono^ and ZSWIM8^Δdimer^ eluted at later fractions, indicating lower molecular weight. ZSWIM8^WT^ and ZSWIM8^mono^ associated with CUL3, whereas ZSWIM8^Δdimer^ did not. Note that fraction 14 was not contiguous with the other fractions. **l,** Requirement of ZSWIM8 dimerization for AGO2 ubiquitylation. Shown is a stepwise ubiquitin transfer assay using fluorescently labeled ubiquitin testing the activity of ZSWIM8^mono^. AGO2–miR-7 complexes were bound to a target RNA derived from CYRANO, which harbored trigger pairing to miR-7. CYRANO trigger (120 nt) was used as the target RNA (Supplementary Table 1). UBE2L3 was charged with ubiquitin, quenched by addition of apyrase, and mixed with remaining components. Ubiquitylation of ZSWIM8^mono^ indicates that ZSWIM8^mono^ is still capable of interacting with CUL3. Ubiquitylation was detected by SDS-PAGE, followed by in-gel fluorescence to visualize fluorescently labeled ubiquitin. Shown is a representative experiment; *n* = 2 technical replicates. Gels stained with Coomassie blue are shown in Supplementary Figure 1. **m,** Requirement of ZSWIM8 dimerization for AGO2 ubiquitylation. Shown is an *in vitro* ubiquitylation assay like that in Figure 1c, but testing the activity of monomeric ZSWIM8 mutants described in **k**. Shown is a representative experiment; *n* = 2 technical replicates. Gels stained with Coomassie blue are shown in Supplementary Figure 1. **n,** Requirement of ZSWIM8 dimerization for TDMD in cells. Shown are results of the intracellular TDMD assay diagrammed and piloted in **d**–**g**, testing mutations affecting ZSWIM8 dimerization, described in **k**. The results are plotted as in **f**. Significance was measured using an ordinary one-way ANOVA with Dunnett’s multiple comparisons tests to compare that of each variant to the mean of WT (\*\*\**P* < 0.0001). *n* = 2 biological replicates.

**Extended Data Figure 4.**
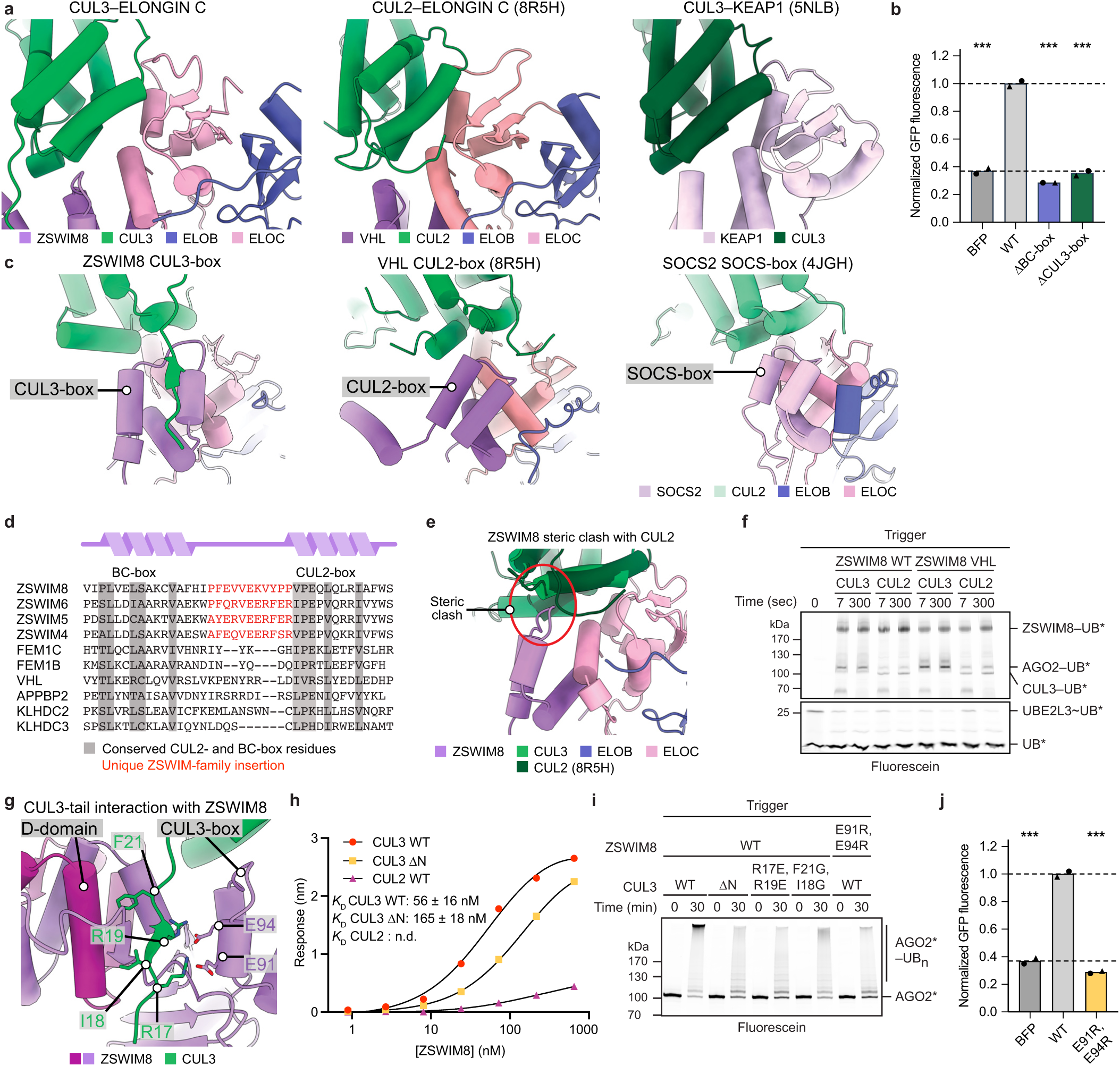
ZSWIM8–CUL3 defines a distinct class of cullin–RING ligases. **a,** Cartoon representations comparing CUL3 binding to ELOC in a ZSWIM8–ELOB/C complex (our structure) (left) with CUL2 binding to ELOC in a VHL–ELOB/C complex (PDB: 8R5H) (middle) and CUL3 binding to KEAP1 (PDB: 5NLB) (right). Complexes were aligned to the first 100 residues of CUL3 in the CUL3–ZSWIM8–AGO–trigger complex. **b,** Importance of ZSWIM8 interactions with ELOB/C and CUL3 for TDMD in cells. Shown is quantification of the intracellular TDMD reporter assay testing mutations in the ZSWIM8 BC-box and CUL3-box, plotted as in Extended Data Figure 3f. Significance was measured using an ordinary one-way ANOVA with Dunnett’s multiple comparisons tests to compare the mean of each variant to that of WT (\*\*\**P* < 0.0001). *n* = 2 biological replicates. **c,** Cartoon representations comparing ZSWIM8 CUL3-box binding to CUL3 (our structure) (left) with VHL CUL2-box binding to CUL2 (PDB: 8R5H) (middle) and SOCS2 SOCS-box binding to CUL5 (PDB: 4JGH) (right). Complexes were aligned to the first 100 residues of CUL3 in the CUL3– ZSWIM8–AGO–trigger complex. **d,** Sequence alignment of ZSWIM8 with other members of the ZSWIM family and with CUL2 substrate receptors. BC- and CUL2-boxes are shown with conserved regions highlighted. A ZSWIM-family-specific insertion between the BC-box and CUL2-box is shown in red. Hence, we annotate ZSWIM family members as containing a unique CUL3-box. **e,** Overlay of the ZSWIM8–CUL3 interaction with CUL2 (PDB: 8R5H), showing a steric clash between the ZSWIM8 CUL3-box and CUL2. The red circle highlights a clash of ZSWIM8 residues and CUL2 N-terminal residues. **f,** Effect of ZSWIM8 cullin-box–cullin pairing on AGO2 ubiquitylation. Shown are results of a stepwise ubiquitin transfer assay using fluorescently labeled ubiquitin testing the activity of a ZSWIM8 variant in which the CUL3-box residues (96–102) were replaced with the VHL CUL2-box residues (177–183) (ZSWIM8 VHL). Otherwise, this panel is as in Extended Data Figure 3l. This substitution improved the ability of CUL2 to function with ZSWIM8. Shown is a representative experiment; *n* = 2 technical replicates. Gels stained with Coomassie blue are shown in Supplementary Figure 1. **g,** Cartoon representation illustrating the interaction of the CUL3 N-terminal tail with ZSWIM8. The CUL3 N terminus forms an intermolecular amphipathic beta sheet with part of the ZSWIM8 D-domain. In addition, the ZSWIM8 D-domain binds to CUL3 via hydrophobic interactions, and ZSWIM8 residues in the CUL3-box form a negatively charged surface. **h,** ZSWIM8 binding to CUL3 and other cullin variants. Shown is biolayer interferometry data comparing binding of ZSWIM8 to CUL3, CUL3 with a truncation of residues 1–24 (CUL3 ΔN), and CUL2. CUL2 and CUL3 N-terminal domains were biotinylated and immobilized; ZSWIM8 dilutions were used as the analyte. The maximal response is plotted against the concentration of ZSWIM8–ELOB/C complex. Binding affinity was determined by fitting to a single-site binding model. Concentrations of ZSWIM8 were too low to estimate a binding affinity for CUL2. *n* = 2 technical replicates. Sensorgrams are shown in Supplementary Figure 7. **i,** Importance of ZSWIM8 interactions with the CUL3 N-terminal tail for AGO2 ubiquitylation. Shown is an *in vitro* ubiquitylation assay like that in Figure 1c but testing the activity of CUL3 variants with mutations in the N-terminal tail (ΔN, truncation of residues 1–24) and a ZSWIM8 variant with mutations in two CUL3-interacting residues. Shown is a representative experiment; *n* = 2 technical replicates. Gels stained with Coomassie blue are shown in Supplementary Figure 1. **j,** Importance of ZSWIM8 interactions with the CUL3 N-terminal tail for TDMD in cells. Shown is quantification of the intracellular TDMD reporter assay testing a ZSWIM8 variant that impairs CUL3 recruitment, plotted as in Extended Data Figure 3f. Significance was measured using an ordinary one-way ANOVA with Dunnett’s multiple comparisons tests to compare the mean of each variant to that of WT (\*\*\**P* < 0.0001). *n* = 2 biological replicates.

**Extended Data Figure 5.**
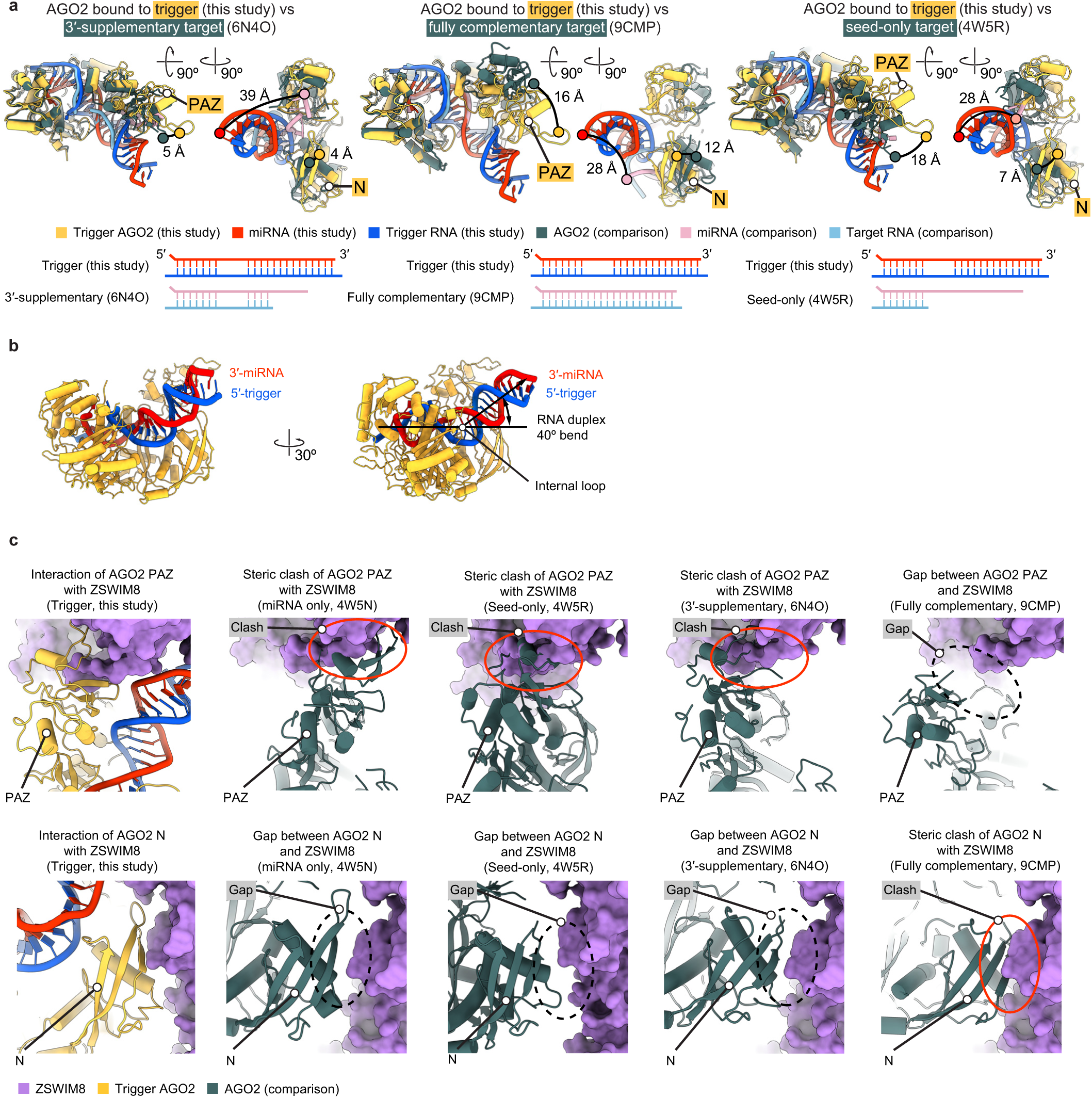
Trigger RNA reshapes the AGO2–miRNA complex into a conformation recognized by ZSWIM8. **a,** Cartoon representations comparing the structure of trigger-bound AGO2–miRNA in association with ZSWIM8–CUL3 to previously determined structures of AGO2– miRNA bound to a target RNA with either 3′-supplementary pairing (PDB: 6N4O) (left); fully complementary pairing (PDB: 9CMP) (middle); or pairing to only the seed (PDB: 4W5R) (right). For each comparison, the cartoon representation on the left highlights the movement of the PAZ domain, the cartoon representation on the right highlights the movements of the PAZ and N domains and the miRNA 3′ terminus from a rotated view, and the lower panel diagrams the W–C–F base pairing between the miRNA and the target RNA. **b,** Cartoon representation illustrating the characteristic internal loop and upward bend of the miRNA–trigger duplex within AGO2. The internal loop enables a 40° deviation from a perfectly straight duplex. The angle was measured between the miRNA 3′ terminus and the extension of the seed helix into a perfectly straight duplex, with the center of rotation placed at the internal loop. **c,** Cartoon and surface representations illustrating steric clashes or gaps formed with the AGO2 PAZ (top) or N (bottom) domains and the ZSWIM8 TDMD sensor when modelling possible interactions with other AGO2–miRNA–target RNA complexes. AGO2 complexes were aligned with residues 349–859 of the L2, MID, and PIWI domains. ZSWIM8 is shown in surface representation. Steric clashes are indicated by red ovals; gaps are indicated by black dashed ovals.

**Extended Data Figure 6.**
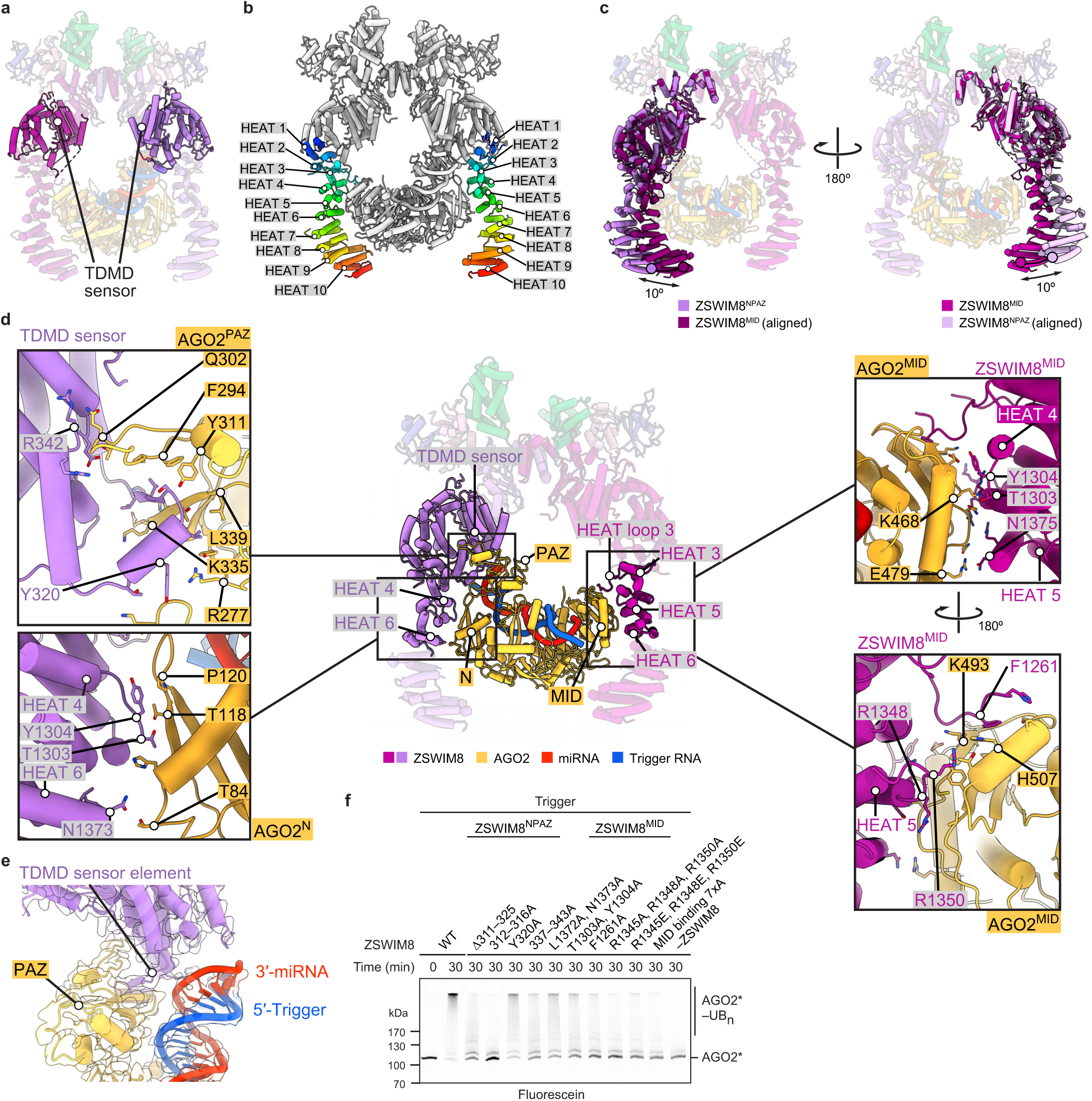
Interactions between the ZSWIM8 dimeric clamp and AGO2 specify TDMD. **a,** Cartoon representation highlighting the position of the TDMD sensor within the ZSWIM8 dimer. The TDMD sensor spans residues 290–915 but is missing density for a large, apparently unstructured region encompassing residues 500–758. **b,** Cartoon representation highlighting the positions of the HEAT repeats within the ZSWIM8 dimer. HEAT repeats are continuously colored from N-terminal repeats to C-terminal repeats. **c,** Cartoon representation illustrating structural rearrangements of the two ZSWIM8 protomers. Each ZSWIM8 protomer was aligned with the other protomer using the E3 ligase superdomain and TDMD sensor (residues 1–915). The maximal relative rotation of each protomer was measured at the most C-terminal alpha helix. **d,** Cartoon representation showing interactions of ZSWIM8^NPAZ^ and ZSWIM8^MID^ with the AGO2 PAZ and MID domains, respectively. Middle: Cartoon representation showing an overview of highlighted interactions. Top left: Cartoon representation of ZSWIM8^NPAZ^ interacting with the PAZ domain. The TDMD sensor element forms an intermolecular beta sheet with a portion of the AGO2 PAZ domain (AGO2 residues 335–337). Bottom left: Cartoon representation showing interaction of ZSWIM8^NPAZ^ with the AGO2 N domain. Multiple residues from HEAT repeats 4 and 6 interact with the N domain. Top right: Cartoon representation showing interaction of ZSWIM8^MID^ with the AGO2 MID domain. Residues in HEAT repeats 3, 4, and 5 interact with the MID domain. Bottom right: Same interaction as in top right, rotated by 180°. **e,** Density showing three-way interactions formed by ZSWIM8^NPAZ^, AGO2 PAZ domain, and the miRNA–trigger duplex. Density is derived from the composite map. The underlying cartoon representation is shown. The TDMD sensor element inserts between the PAZ domain and the RNA duplex. **f,** Importance of ZSWIM8 clamp interactions with AGO2. Shown is an *in vitro* ubiquitylation assay like that in Figure 1c, except it tested the consequences of mutating ZSWIM8^NPAZ^ and ZSWIM8^MID^ residues that interact with AGO2. Shown is a representative experiment; *n* = 2 technical replicates. Gels stained with Coomassie blue are shown in Supplementary Figure 8.

**Extended Data Figure 7.**
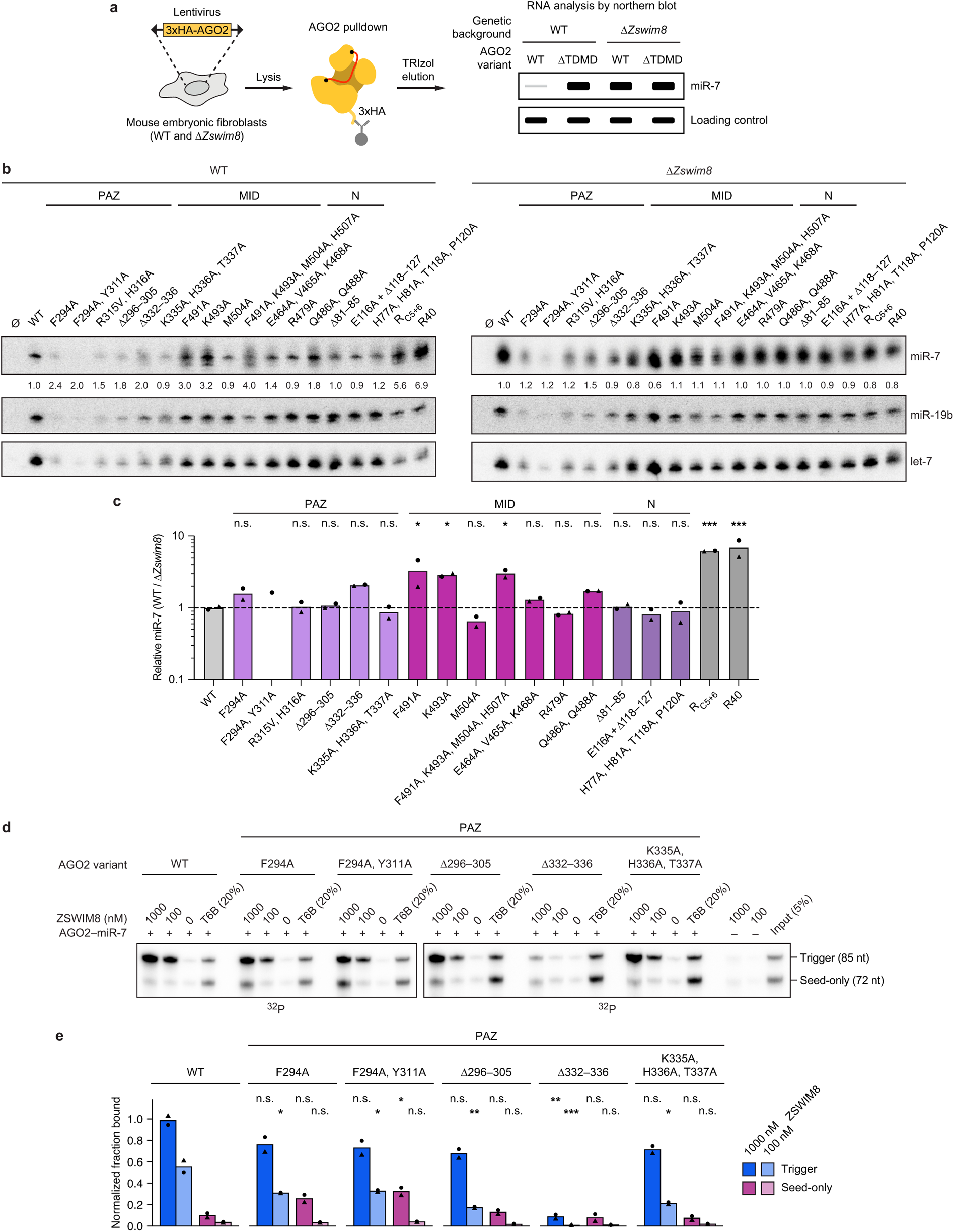
AGO2 residues make important interactions with ZSWIM8. **a,** Schematic of the intracellular AGO2 co-IP assay for measuring the levels of miR-7 associated with an AGO2 variant^23^. Wild-type (WT) or *Zswim8*-knockout (Δ*Zswim8*) mouse embryonic fibroblasts (MEFs) are transduced with a lentiviral construct expressing an epitope-tagged AGO2 variant of interest (3xHA-AGO2). After selection, cells are harvested and an IP for 3xHA is performed to enrich for RNAs associated with the ectopically expressed AGO2 variant. These co-precipitating RNAs are then analyzed by northern blot. The relative amounts of miR-7 associated with each AGO2 variant in WT versus Δ*Zswim8* MEFs reports on the ZSWIM8 sensitivity of the respective AGO2 variant. For example, an AGO2 variant that cannot undergo TDMD (ΔTDMD) is expected to be associated with elevated miR-7 levels in WT cells, and these levels are not expected to be further elevated in Δ*Zswim8* cells, whereas an AGO2 variant that can fully undergo TDMD is expected to resemble WT AGO2, in being associated with a low level of miR-7 in WT cells and an increased level of miR-7 in Δ*Zswim8* cells. **b,** Effect of mutations in AGO2 ZSWIM8-interacting residues on miR-7 levels in cells, assessed as diagrammed in **a**. Shown are representative northern blots measuring levels of miR-7 and control miRNAs (miR-19b, let-7) in IPs from WT and Δ*Zswim8* MEFs expressing the indicated 3xHA-AGO2 variant (Ø, untransduced; R_C5+6_ and R40, previously characterized AGO2 variants with lysine-to-arginine substitutions) (Supplementary Table 3). For each AGO2 variant, the ratio of miR-7:miR-19b and the ratio of miR-7:let-7 was calculated. Shown is the geometric mean of these two ratios normalized to that of WT AGO2. AGO2 levels were detected by western blotting (Supplementary Figure 9). **c,** Quantification of AGO2 co-IP assays. For each AGO2 variant, the geometric mean of the miR-7:miR-19b and miR-7:let-7 ratios in WT MEFs was normalized to that in Δ*Zswim8* MEFs, and this ratio was normalized to the geometric mean of normalized ratios across two WT AGO2 replicates. The bar height represents the geometric mean of two independent measurements observed in two pairs of independently derived WT and Δ*Zswim8* clonal cell lines. Significance was measured using an ordinary one-way ANOVA with Dunnett’s multiple comparisons tests to compare the mean of each variant to the mean of WT (*P < 0.05, ***P < 0.0001; n.s., not significant). *n* = 2 biological replicates. With the F294A, Y311A variant, one of the replicates did not have detectable signal above background, and thus the relative miR-7 levels of this replicate could not be calculated. The quantification for the other replicate is indicated by a black circle. **d,** Importance of AGO2 PAZ-domain interactions for ZSWIM8 binding. Shown are *in vitro* co-IP assays as in Figure 1f, except wild-type AGO2–miR-7 was replaced with complexes formed using the indicated variants in the AGO2 PAZ domain. *n* = 2 technical replicates. **e,** Quantification of co-IP assays of panel **d**. Plotted are measurements for radiolabeled target RNA bands associated with each AGO2–miR-7 variant. Band intensities were first background-subtracted using the 0 nM ZSWIM8 sample and normalized to those in the T6B sample. The symbols show data from independent measurements. The bar height represents the mean of these measurements. Significance was measured using an ordinary one-way ANOVA with Dunnett’s multiple comparisons tests to compare for each RNA (trigger or seed-only) the mean of each fraction that co-IPed with a given AGO2 variant to that which co-IPed with WT AGO2 at the corresponding ZSWIM8 concentration (\**P* < 0.05, \*\**P* < 0.001, \*\*\**P* < 0.0001; n.s., not significant). *n* = 2 technical replicates.

**Extended Data Figure 8.**
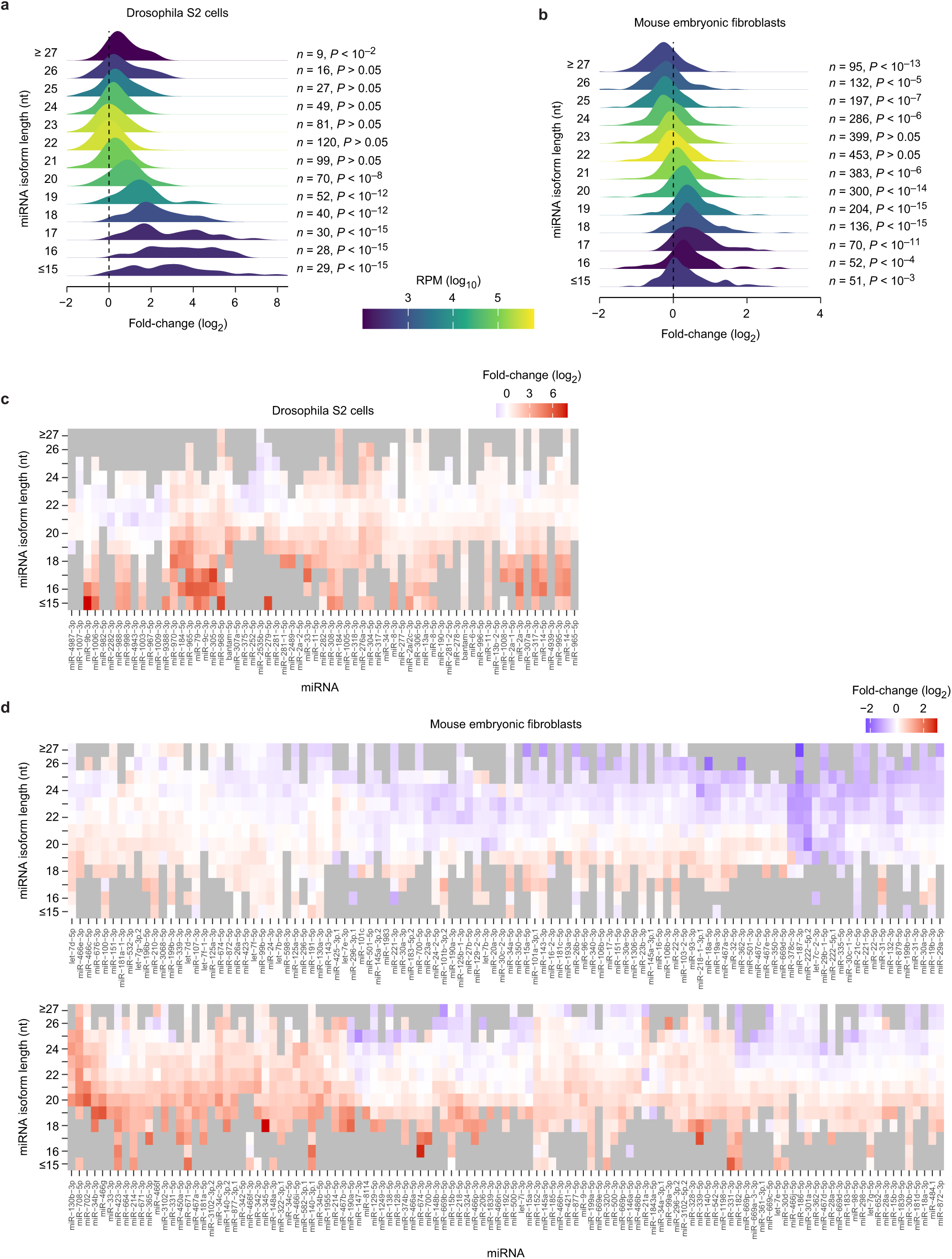
ZSWIM8 recognizes an unoccupied AGO2 PAZ-domain pocket. **a,** Relationship between miRNA isoform length and ZSWIM8 sensitivity in Drosophila cells. The distributions show fold-changes (log2) in levels of the indicated miRNA length isoforms in *Zswim8*-knockout Drosophila S2 cells, relative to control cells (*n* = 3 biological replicates). This analysis excludes miRNAs with evidence of ZSWIM8 sensitivity of full-length isoforms, for reasons described in the following paragraph. Fold-changes were adjusted to center the median of the 22-nt distribution at zero, and distributions were colored by the summed expression in control cells of isoforms composing each bin (key). Isoforms longer than 26 nt or shorter than 16 nt were aggregated into the two terminal bins. Each distribution of log2 fold-changes was compared with the distribution of log2 fold-changes generated by summing the counts for all lengths of a given miRNA using a two-sided Kolmogorov-Smirnov test. The resulting Benjamini-Hochberg-adjusted *P* values and the numbers of different miRNAs contributing to each distribution are listed. The greater fold-changes observed for shorter isoforms support the idea that shorter isoforms are broadly susceptible to ZSWIM8-mediated degradation. One consideration potentially confounding this interpretation is that some TDMD sites also promote the removal or the untemplated addition of nucleotides to the 3′ end of miRNAs, which is presumably due to liberation of the miRNA 3′ end from AGO upon extensive 3′ pairing to a trigger RNA. In the absence of ZSWIM8 or its orthologs, these trimmed or tailed isoforms can accumulate. To maintain focus on a potential effect of miRNA length on ZSWIM8 sensitivity —rather than changes in miRNA length downstream of trigger binding —we sought to exclude from the analysis miRNAs with evidence for classical ZSWIM8 sensitivity. To this end, we summed the reads associated with all lengths of a given miRNA, calculated fold-changes in ZSWIM8-knockout versus control cells for each miRNA, and excluded all putative ZSWIM8-sensitive miRNAs (those with log_2_ fold-change > 0 and *P*_adj_ < 0.05, those having undergone an increase significantly larger than that of their passenger strand, and those previously annotated as ZSWIM8-sensitive). ZSWIM8-dependent fold-changes for all isoforms of the remaining, insensitive miRNAs were calculated. Analogous plots for miRNAs whose levels are significantly or potentially affected by ZSWIM8 are shown in Supplementary Figure 13. **b,** Relationship between miRNA isoform length and ZSWIM8 sensitivity in murine cells. The distributions show fold-changes (log_2_) in miRNA levels in *Zswim8*-knockout mouse embryonic fibroblasts (MEFs), relative to control cells (*n* = 3 biological replicates). This panel is as in **a**, except analysis is of data from MEFs. **c,** Heatmap of fold-changes (log_2_) of miRNA molecules of the indicated lengths in *Zswim8*-knockout versus control S2 cells for all miRNAs shown in **a** that have at least four isoforms of different lengths, ordered by similarity based on Euclidean distance. **d,** Heatmap of fold-changes (log_2_) of miRNA molecules of the indicated lengths in *Zswim8*-knockout versus control MEFs for all miRNAs shown in **b** that have at least six isoforms of different lengths, plotted as described in **c**. In both cell types, shorter isoforms (< 22 nt) accumulated upon loss of ZSWIM8 to a greater extent than the dominant miRNA isoforms, which were typically 22 nt. In Drosophila S2 cells and MEFs, < 19-nt guides and 17–19-nt guides, respectively, underwent the highest mean ZSWIM8-associated degradation. Of note, *in vitro* measurements of miRNA 3′ end accessibility to nucleases indicate that miRNAs typically require 18–19 nt to be stably bound by the PAZ domain of human AGO2 and AGO3, depending on the primary sequence of the 3′-terminal nucleotides^81^. Thus, our data are consistent with the idea that short (< 20 nt) guides whose 3′-terminal nucleotides are unable or less likely to occupy the PAZ binding pocket in AGO are recognized by ZSWIM8 through an interaction between ZSWIM8 and PAZ, resulting in miRNA degradation. However, despite stringent removal of putative ZSWIM8 substrates from the analyses, we cannot fully rule out the possibility that the preferential accumulation of short guides in ZSWIM8 knockouts results from trimming of weakly ZSWIM8-sensitive miRNAs that have extensive 3′ pairing to unknown trigger RNAs. Similarly, if the extent of trimming or tailing increases over the lifetime of a miRNA, miRNAs even mildly destabilized by ZSWIM8 could accumulate more trimmed and tailed isoforms in the absence of ZSWIM8.

**Extended Data Figure 9.**
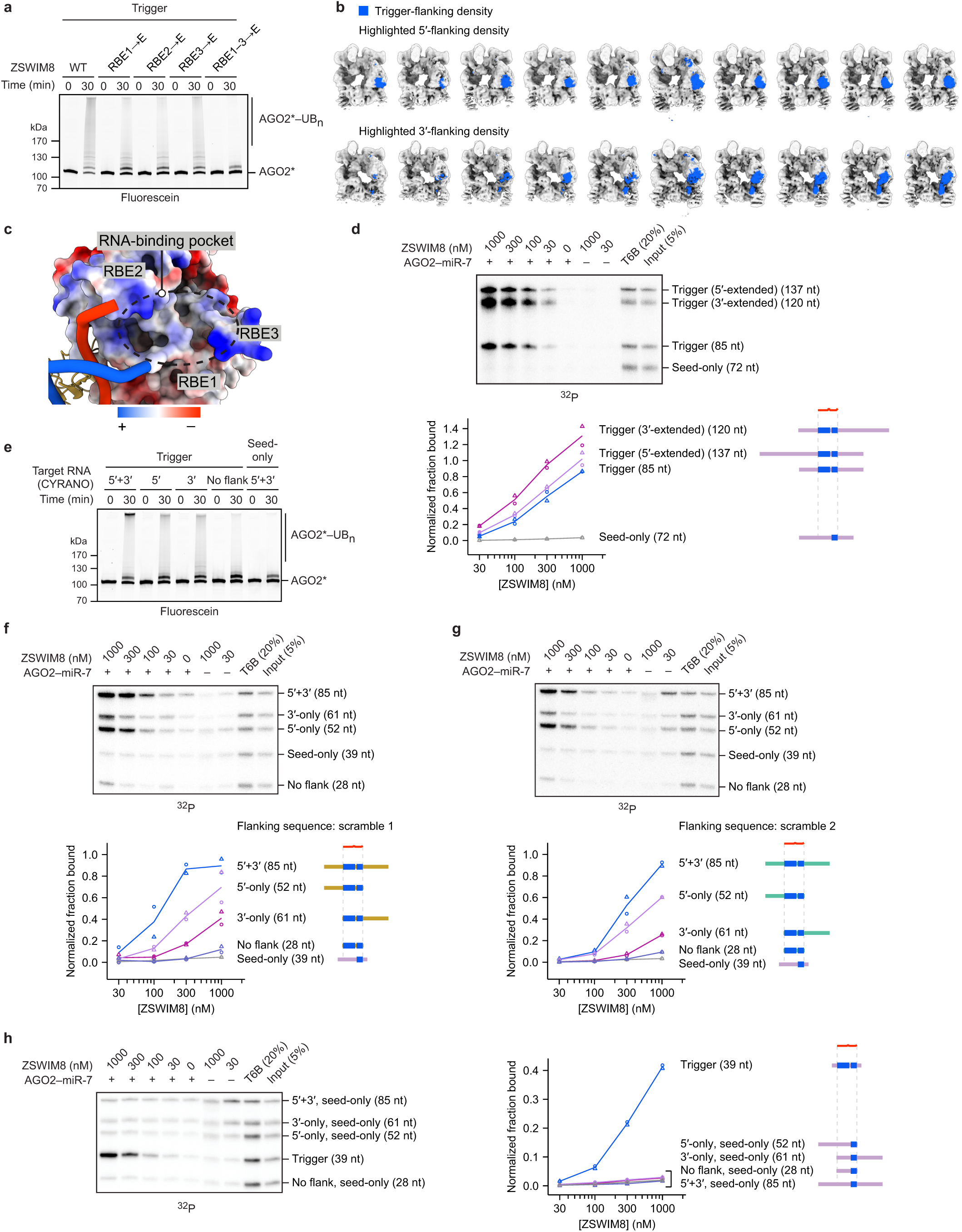
Flanking trigger RNA interacts with ZSWIM8. **a,** The importance of RBEs for AGO2 ubiquitylation. Shown are *in vitro* ubiquitylation assays like that in Figure 1c, except ZSWIM8 was mutated at the indicated RBE or at all three RBEs. Residues in RBEs were mutated to glutamates (E). Shown is a representative experiment; *n* = 2 technical replicates. Gels stained with Coomassie blue are shown in Supplementary Figure 8. **b,** Cryo-EM densities obtained by 3D classification showing different extents of additional density assigned as either 5′- or 3′-flanking trigger RNA (top and bottom, respectively). Densities assigned as RNA are shown in blue. The density assigned as 3′-flanking RNA showed considerable variability in its volume and position, indicating structural heterogeneity. **c,** Charge-driven interactions of ZSWIM8 RBEs with RNA. Shown is a surface rendering of ZSWIM8^NPAZ^ showing the calculated Coulombic potential of RBEs 1–3 positioned to interact with the miRNA–trigger duplex and flanking trigger RNA. Coulombic potential is shown in the range of –10–10 kcal/(mol·*e*). **d,** Testing the effect of further extending the trigger flanking regions on ZSWIM8 binding to AGO2–miR-7–CYRANO. This panel is as in Figure 1f, except the 85-nt trigger RNA was extended in either direction by additional CYRANO sequence. In addition, a higher concentration of heparin (10 µg/mL) was included. *n* = 2 technical replicates. **e,** Contributions of 5′ and 3′ trigger flanking sequences to AGO2 ubiquitylation. Shown are *in vitro* ubiquitylation assays like that in Figure 1c, except the target RNA had CYRANO sequence flanking the trigger site at either both ends, one end, or neither end. CYRANO trigger 5′+3′ (85 nt), 5′-only (52 nt), 3′-only (61 nt), no flank (28 nt), and seed-only (39 nt) target RNAs were used (Supplementary Table 1). Shown is a representative experiment; *n* = 2 technical replicates. Gels stained with Coomassie blue are shown in Supplementary Figure 8. **f,** Testing sequence specificity of the contributions of trigger flanking sequences to ZSWIM8 binding to AGO2–miR-7–CYRANO. This panel is as in Figure 4e, except the flanking regions derived from CYRANO were replaced with scrambled sequences that maintained the nucleotide composition of the original CYRANO sequence. In addition, heparin was not included in the reaction. *n* = 2 technical replicates. **g,** Testing sequence specificity of the contributions of trigger flanking sequences to ZSWIM8 binding to AGO2–miR-7–CYRANO. This panel is as in **f** but with different scrambled sequences (scramble 2). *n* = 2 technical replicates. **h,** Testing the contributions of trigger flanking sequences to ZSWIM8 binding to AGO2–miR-7–CYRANO^Seed-only^. This panel is as in Figure 4e, except most target RNAs contained seed-only pairing instead of trigger pairing, and heparin was not included in the reaction. *n* = 2 technical replicates.

**Extended Data Figure 10.**
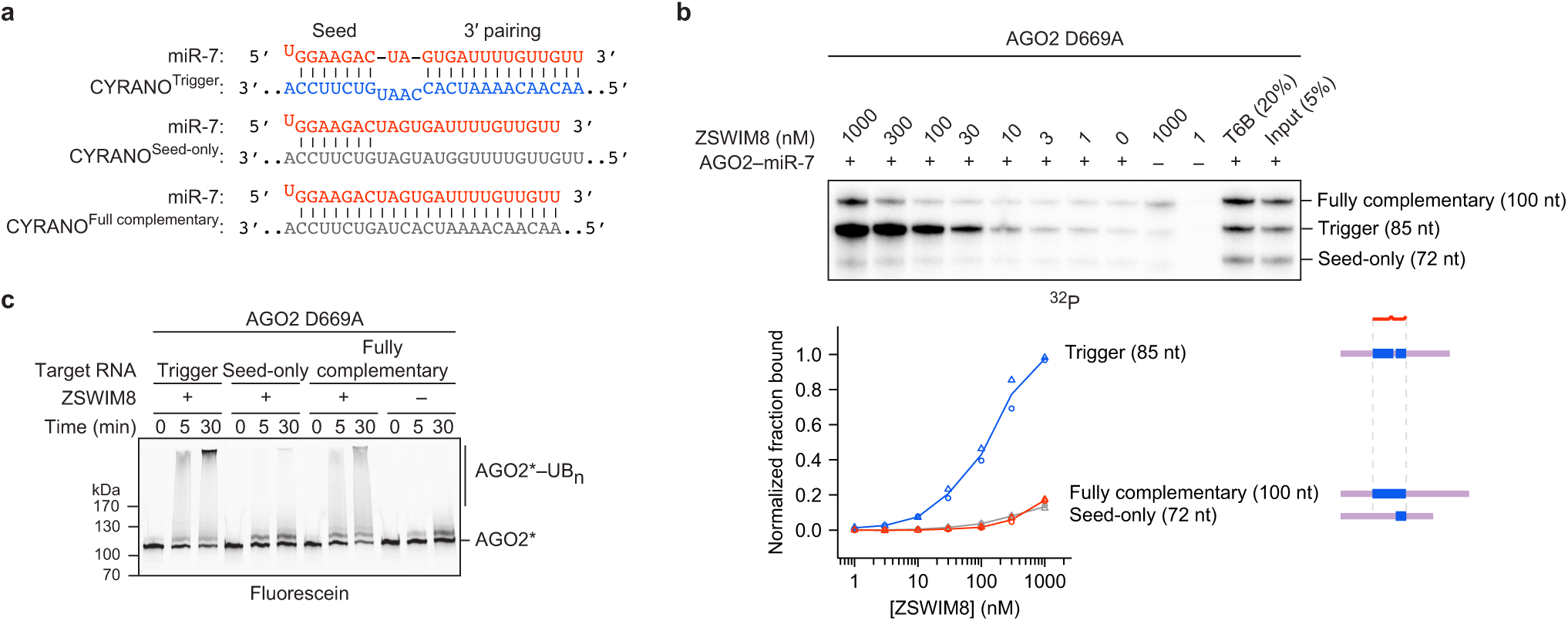
ZSWIM8 recognizes trigger-specific RNA trajectory. **a,** Diagrams showing pairing between miR-7 and the miRNA-binding regions of CYRANO^Trigger^, CYRANO^Seed-only^, and CYRANO^Fully complementary^ target RNAs used for *in vitro* ubiquitylation and co-IP assays. Vertical lines represent W–C–F base pairing. **b,** Effect of fully complementary target pairing on ZSWIM8 binding to AGO2–miR-7 complexes. Shown is an *in vitro* co-IP assay like that in Figure 1f, except it used an active-site AGO2 variant (D669A) and included a 100-nt radiolabeled target RNA with full complementarity to miR-7 (panel **a**) (Supplementary Table 1). The symbols show data from independent measurements. *n* = 2 technical replicates. **c,** Effect of fully complementary target pairing on AGO2 ubiquitylation. Shown is an *in vitro* ubiquitylation assay like that in Figure 1c, but also testing the effects of fully complementary pairing to miR-7 (panel **a**) using an active-site AGO2 variant (D669A). CYRANO trigger (120 nt), seed-only (135 nt), and fully complementary (100 nt) target RNAs were used (Supplementary Table 1). Shown is a representative experiment; *n* = 2 technical replicates. Gels stained with Coomassie blue are shown in Supplementary Figure 8.

**Supplementary Figure 1.**
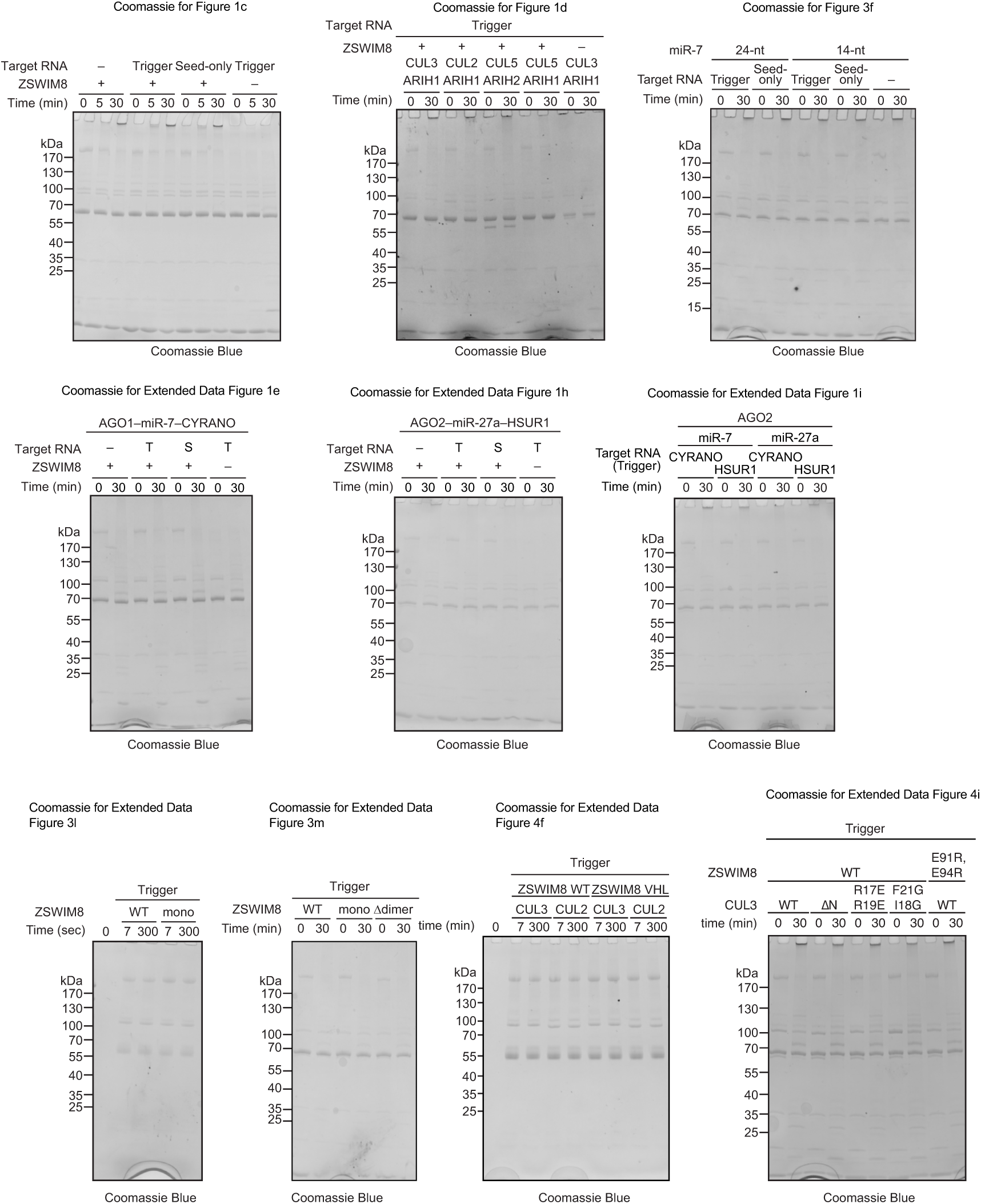
Gels stained with Coomassie Blue for ubiquitylation assays shown in Figure 1c, 1d, and 3f, and Extended Data Figures 1e, 1h, 1i, 3l, 3m, 4f, and 4i.

**Supplementary Figure 2.**
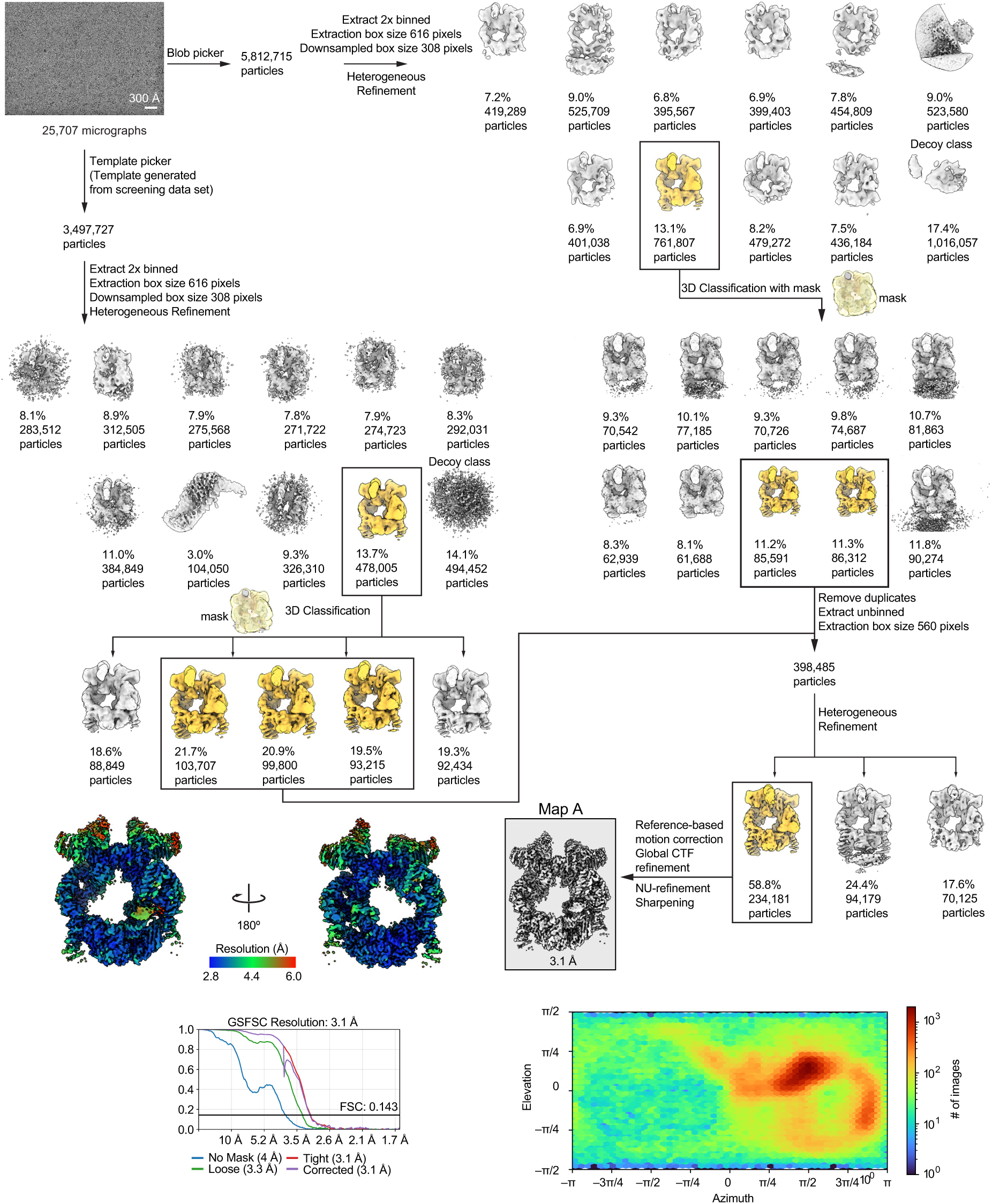
Processing scheme for AGO2–miR-7–CYRANO and ZSWIM8–CUL3 complex. Representative micrograph is shown. Classes selected from classification step are shown in yellow. Masks for masked classification are shown in transparent yellow. Heterogeneous refinements contained one decoy class. Local resolution of Map A is shown. Bottom left, Gold-standard Fourier shell correlation (GSFSC) is shown at a cut-off of 0.143 providing a resolution of 3.1 Å. Bottom right, orientation distribution plot of particles used to generate Map A.

**Supplementary Figure 3.**
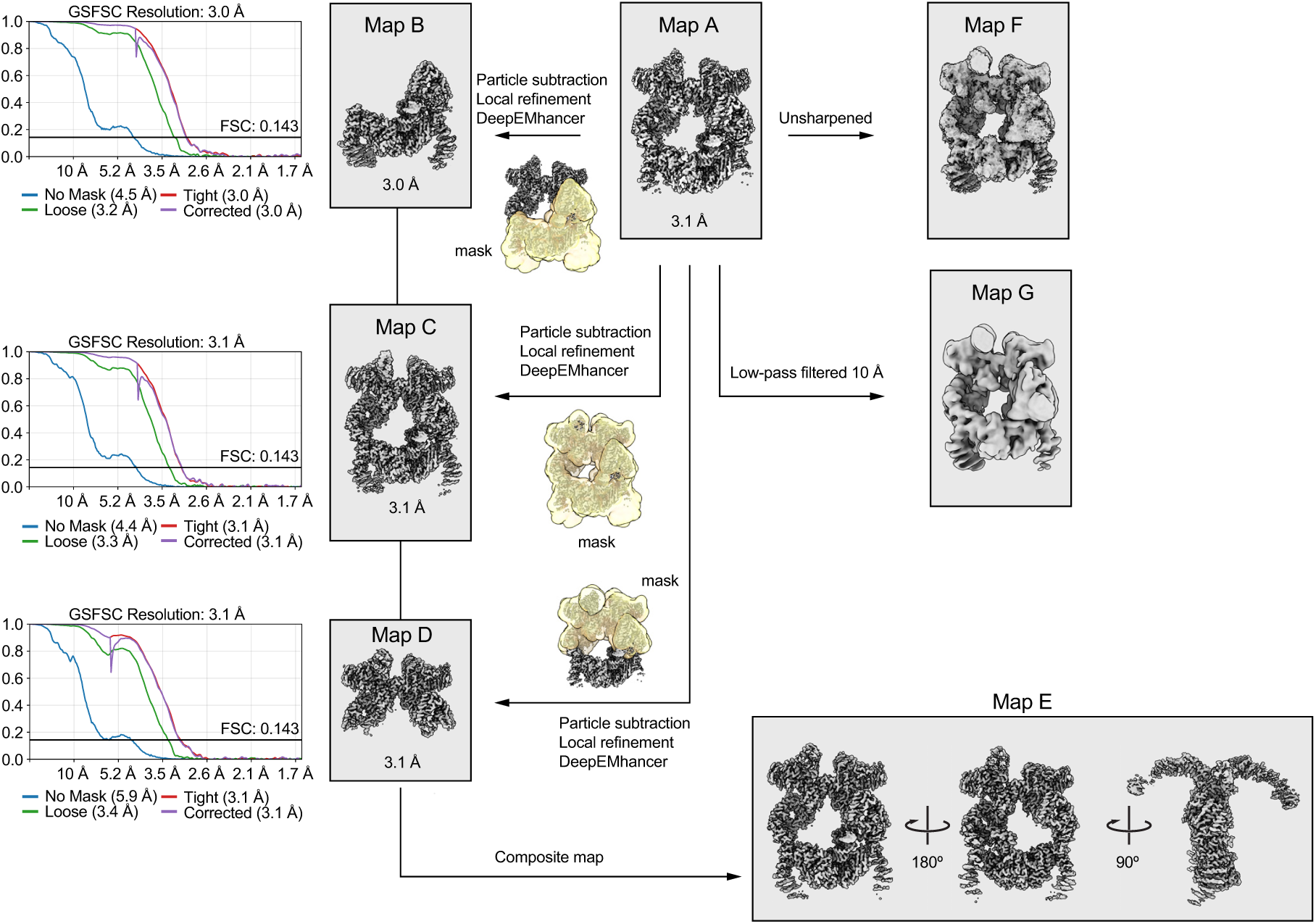
Processing scheme to generate maps derived from consensus map A. Focused refinements to generate maps B, C, and D are shown. Masks used for focused refinement are shown in yellow. Gold standard Fourier shell correlation (GSFSC) is shown at a cut-off of 0.143. Composite map E was generated from maps B, C, and D. Unsharpened map F and low-pass filtered map G were derived from consensus map A.

**Supplementary Figure 4.**
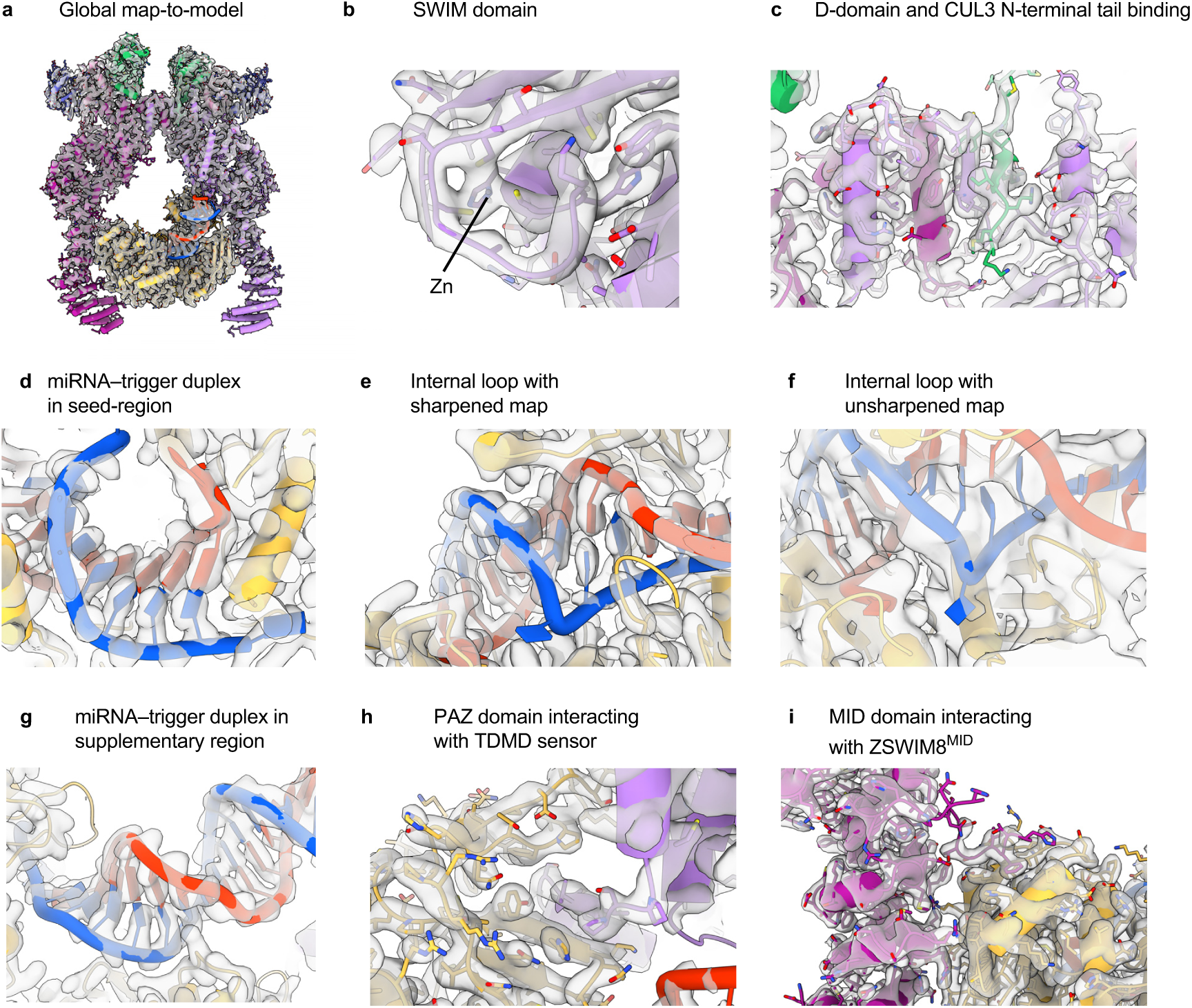
Map-to-model fit of composite Map E to the atomic model. For each panel, the model is colored as in Figure 2. **a**, Global map-to-model fit. **b**, Fit of the SWIM domain. The zinc ion is highlighted for clarity. **c**, Fit of the dimerization domain (D-domain) and the associated CUL3 N-terminal tail. **d–g**, Fit of the RNA model to the density in several regions: **d**, seed region; **e**, internal loop with sharpened Map E; **f**, internal loop with unsharpened Map F; **g**, distal region. **h**, Fit of the TDMD sensor of the ZSWIM8^NPAZ^ domain binding to the AGO2 PAZ domain. **i**, Fit of the ZSWIM8^MID^ domain interaction with the AGO2 MID domain.

**Supplementary Figure 5.**
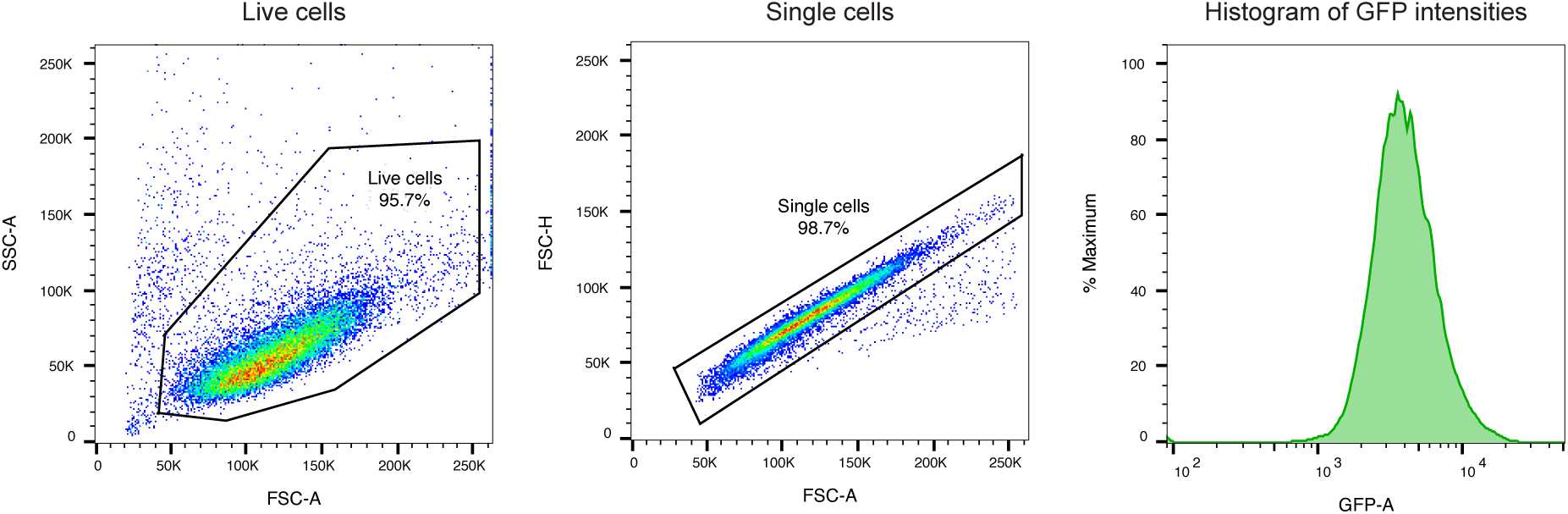
Gating strategy for the ZSWIM8 intracellular rescue assay. Shown are representative plots from cells expressing wild-type ZSWIM8. Cells were gated to obtain live, single cells (left and middle, respectively). The GFP fluorescence was recorded for this subpopulation of cells (right). Approximately 20,000 live, single cells were analyzed for each sample.

**Supplementary Figure 6.**
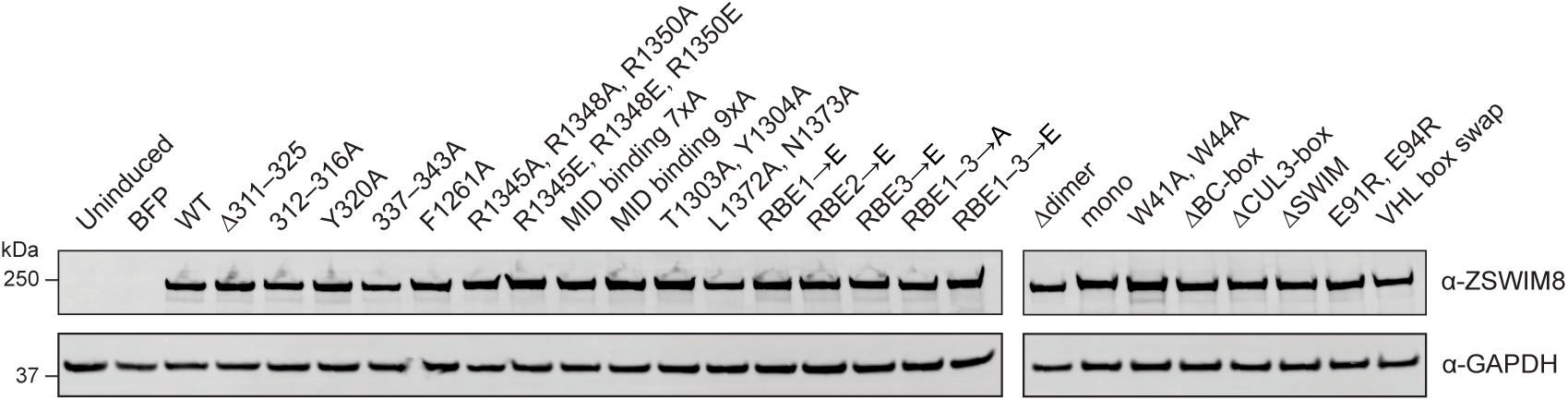
Levels of ZSWIM8 protein variants used in the intracellular TDMD reporter assays shown in Figures 3c and 4c, and Extended Data Figures 3e–g, 3n, 4b, and 4j. Shown are representative western blots detecting expression of each ZSWIM8 variant as well as of GAPDH, which served as a loading control. All ZSWIM8 variants were expressed at levels comparable to that of WT ZSWIM8.

**Supplementary Figure 7.**
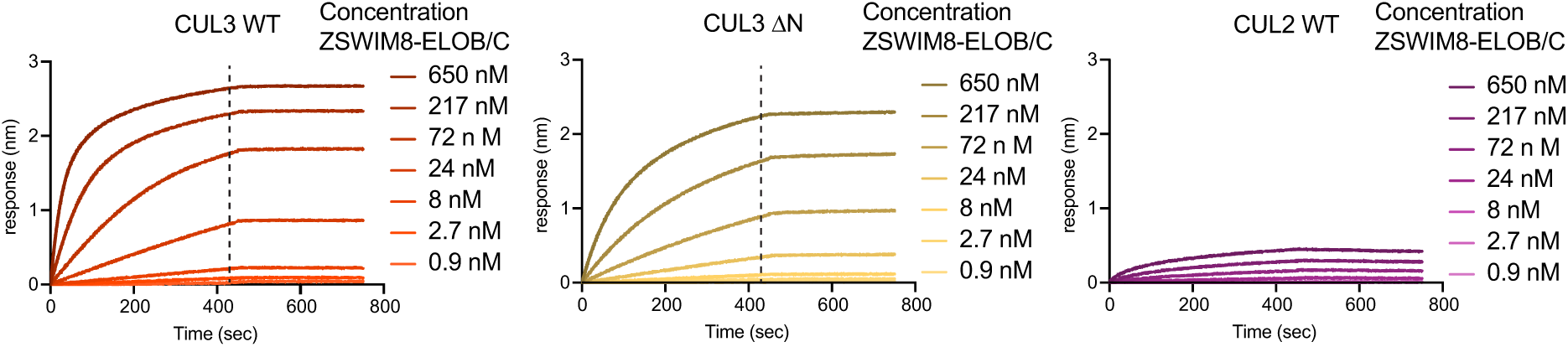
Biolayer interferometry measurements to determine binding affinity of ZSWIM8 for CUL3 WT, CUL3 ΔN (lacking residues 1–24), and CUL2 WT N-terminal domains (Extended Data Figure 4h). Sensorgrams were normalized to the start of the association step. The dashed line indicates the time point at which the maximum response was measured and used for the determination of binding affinity. The dissociation step starting at 430 seconds did not induce any dissociation of ZSWIM8.

**Supplementary Figure 8.**
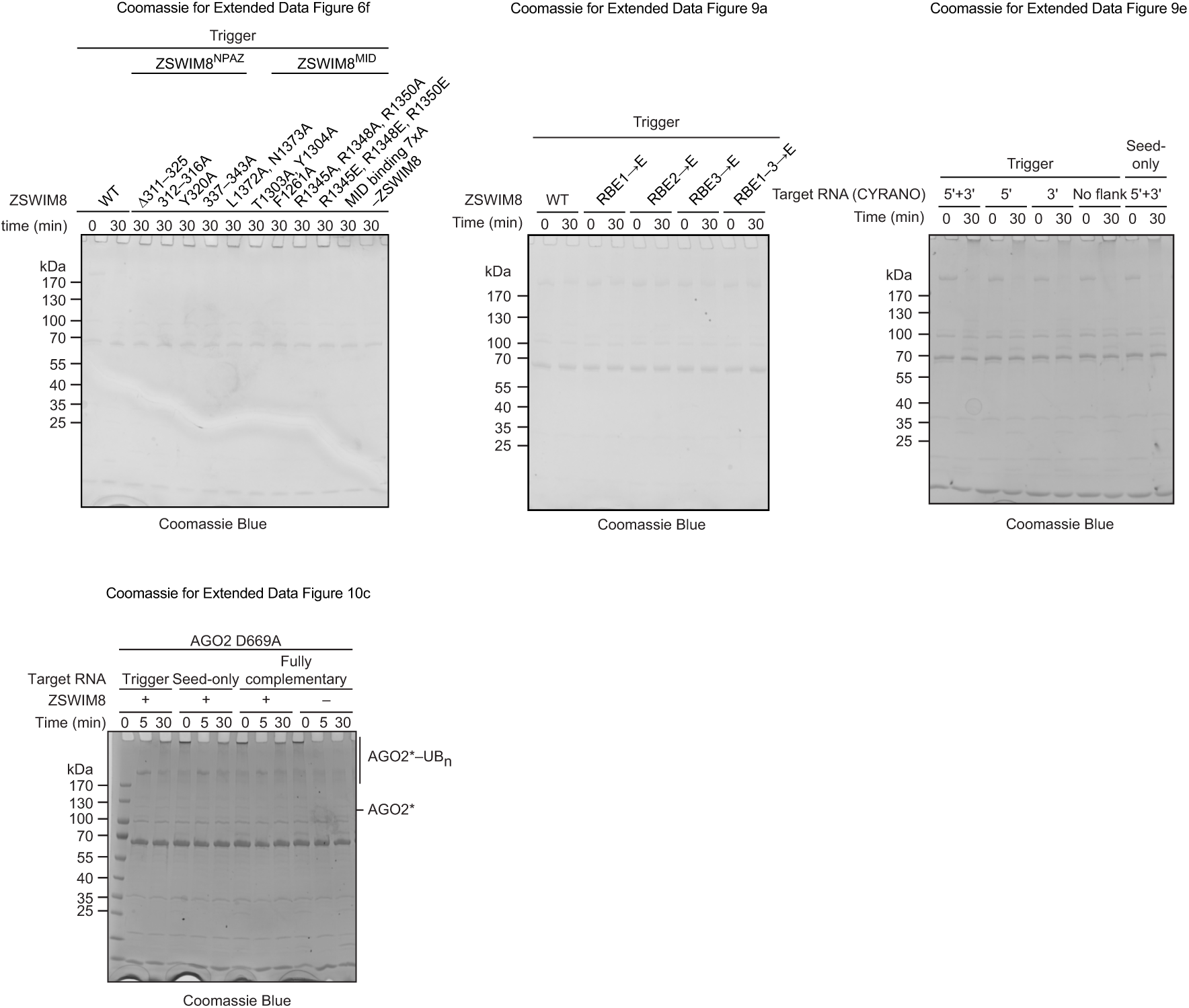
Gels stained with Coomassie Blue for ubiquitylation assays shown in Extended Data Figures 6f, 9a, 9e, and 10c.

**Supplementary Figure 9.**
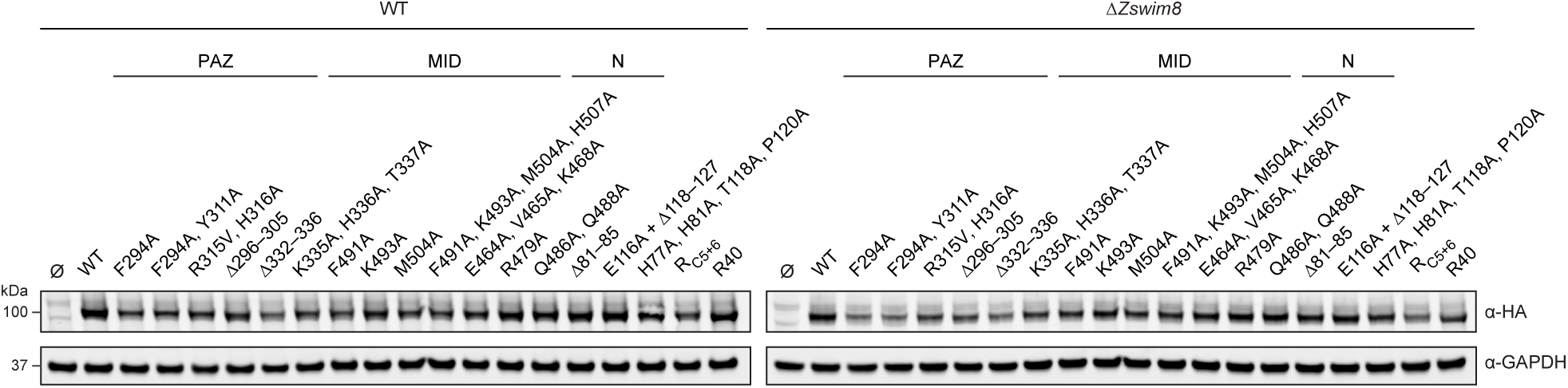
Levels of AGO2 protein variants used in the intracellular AGO2 co-IP assay shown in Extended Data Figure 7b and c. Shown are representative western blots detecting expression of each HA-tagged AGO2 variant as well as of GAPDH, which served as a loading control. All AGO2 variants were expressed at levels comparable to that of WT AGO2.

**Supplementary Figure 10.**
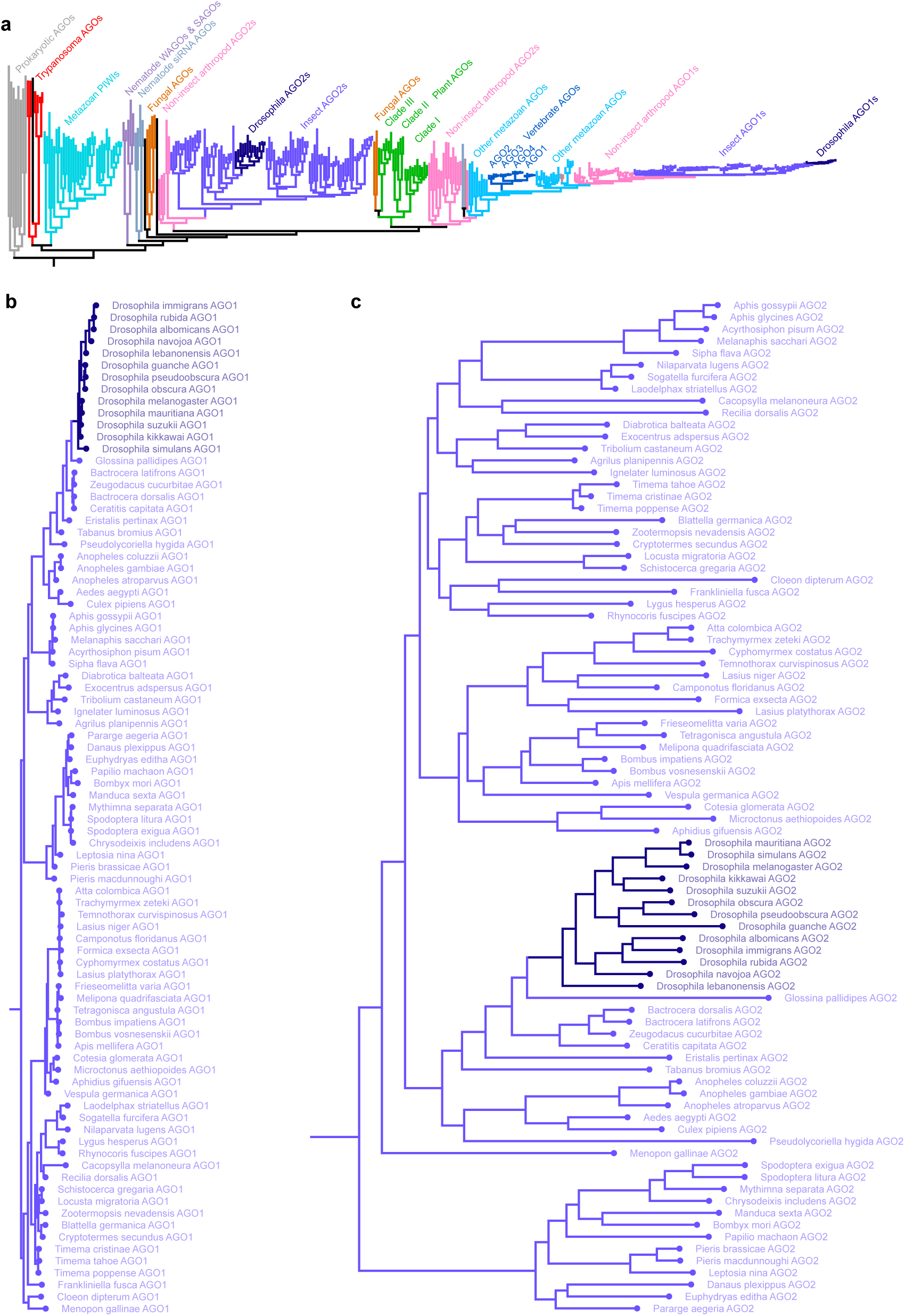
Phylogenetic comparison of insect AGO homologs with different TDMD competencies. **a**, Phylogenetic tree of 347 homologs of AGO-family proteins, including some prokaryotic AGOs, Trypanosoma AGOs, nematode-specific AGOs, metazoan PIWIs, and AGOs from plants, fungi, metazoans, and other eukaryotes. Among these homologs are 85 matched pairs of AGO1 and AGO2 homologs in insects, including those from 13 Drosophila species. Phylogeny was calculated using FastTree 2.2 (ref. 86) based on a multiple-sequence alignment of protein sequences obtained from UniProt^87^, aligned using the MUSCLE algorithm^88^ with the SnapGene software. Homolog groups of interest are labelled and highlighted in distinct colors. Branch lengths are scaled to the rate of amino-acid changes between homologs. **b**, Phylogenetic tree of 85 insect AGO1 proteins, subsetted and magnified from **a**. Colors are as in **a**. Drosophila homologs are highlighted in dark purple. **c**, Phylogenetic tree of 85 insect AGO2 proteins, otherwise as in **b**, from the same 85 insect species and at the same branch-length scale. Branch lengths for insect AGO2 homologs, which are not thought to be subject to TDMD (based on studies of the fly protein)^89^, are longer than those for insect AGO1 homologs, which are subject to TDMD^17^. The shorter branch lengths of AGO1 indicates that its sequence is much more evolutionarily constrained.

**Supplementary Figure 11.**
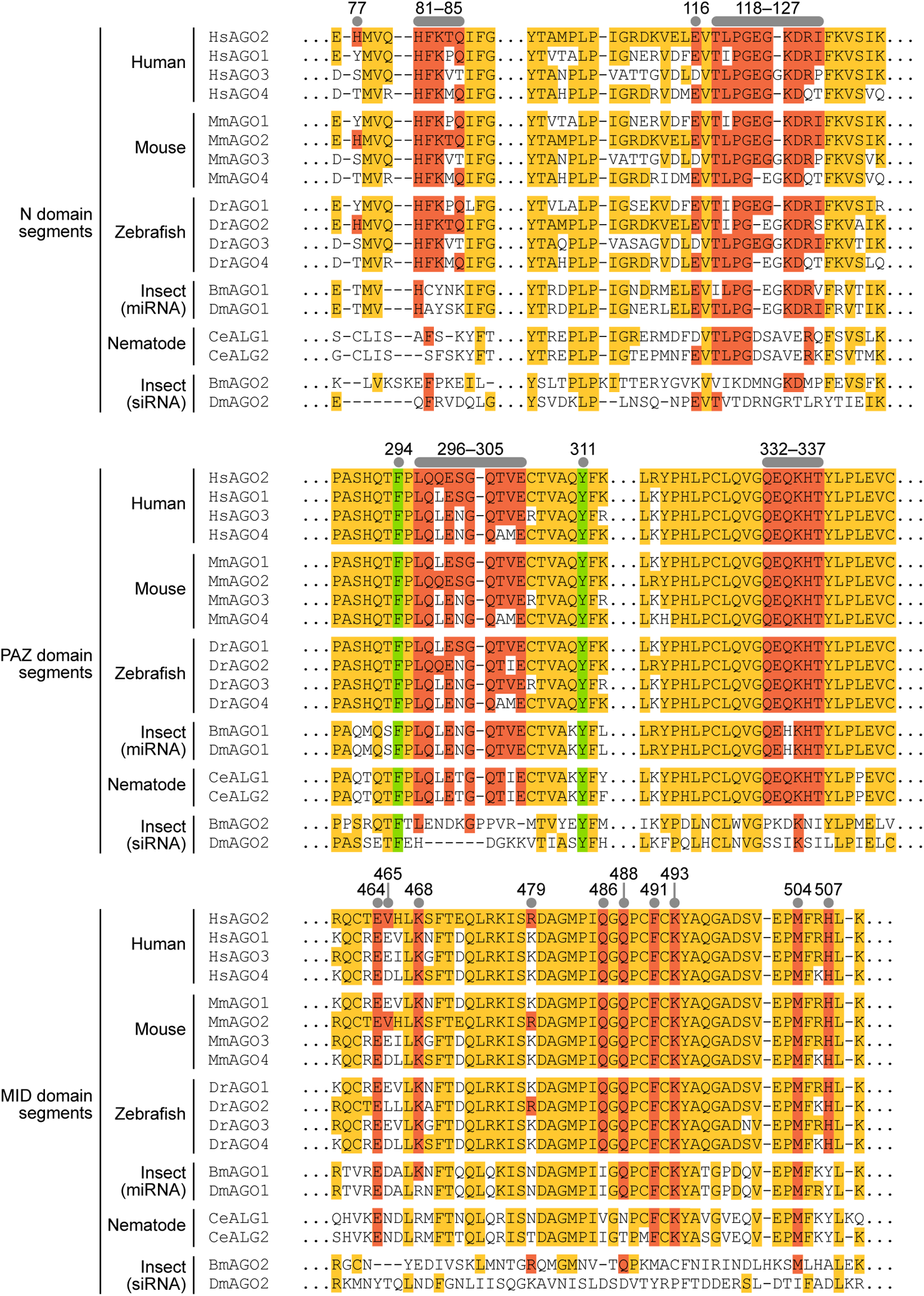
Conservation of AGO protein sequence among homologs in diverse bilaterian species. Shown is a multiple-sequence alignment of regions that bind the ZSWIM8 protein, extracted from a sequence alignment of 111 AGO and PIWI homologs among metazoans, plants, fungi, and prokaryotes^76^. Residues identical to human AGO2 (HsAGO2) are highlighted in orange or red. Human AGO2 residues identified as ZSWIM8 contacts and assayed by substitutions are highlighted in red, as are homologous residues with conserved identity in other proteins. AGO residues identified as ZSWIM8 contacts are also marked with circles and with numbers indicating the position of the corresponding human AGO2 residue; F294 and Y311 were assayed to investigate the effect of a vacated PAZ pocket but not as ZSWIM8 contacts, and so are highlighted in green instead. For comparison, results for insect AGO2 are also included. Insect AGO2 arose from an evolutionary lineage distinct from that of insect AGO1 and other metazoan AGOs^90^ (Supplementary Figure 10). It preferentially loads siRNAs instead of miRNAs and, based on studies of the fly protein^89^, is not thought to be subject to TDMD.

**Supplementary Figure 12.**
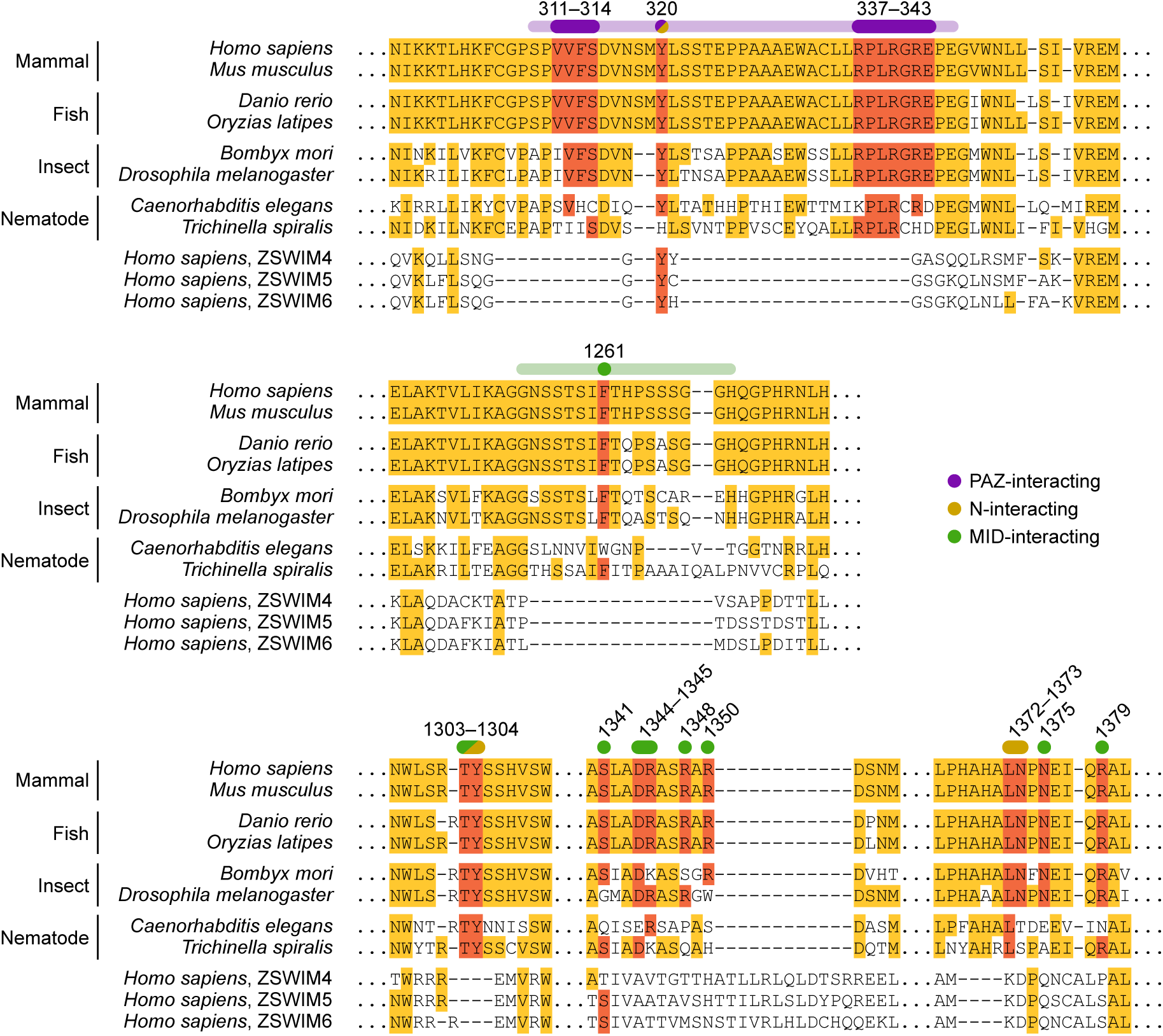
Conservation of ZSWIM8 protein sequence among orthologs in diverse bilaterian species, Three human paralogs (ZSWIM4, ZSWIM5, and ZSWIM6) are also included for comparison. Shown is a multiple-sequence alignment of regions that bind the AGO protein, extracted from a sequence alignment of 1532 full-length ZSWIM4/5/6/8 metazoan homologs. Residues identical to human ZSWIM8 are highlighted in orange or red. Human ZSWIM8 residues that contact AGO are highlighted in red, as are homologous residues with conserved identity in other proteins. ZSWIM8 residues identified as AGO contacts are also marked with numbers indicating the position of the corresponding human ZSWIM8 residue, and with circles colored based on the domain of AGO with which they interact (key). ZSWIM4, 5, and 6, although related to ZSWIM8, are not thought to mediate TDMD. Indeed, ZSWIM8 residues that interact with AGO2 are mostly found in insertions unique to ZSWIM8 and not present in other members of the ZSWIM family.

**Supplementary Figure 13.**
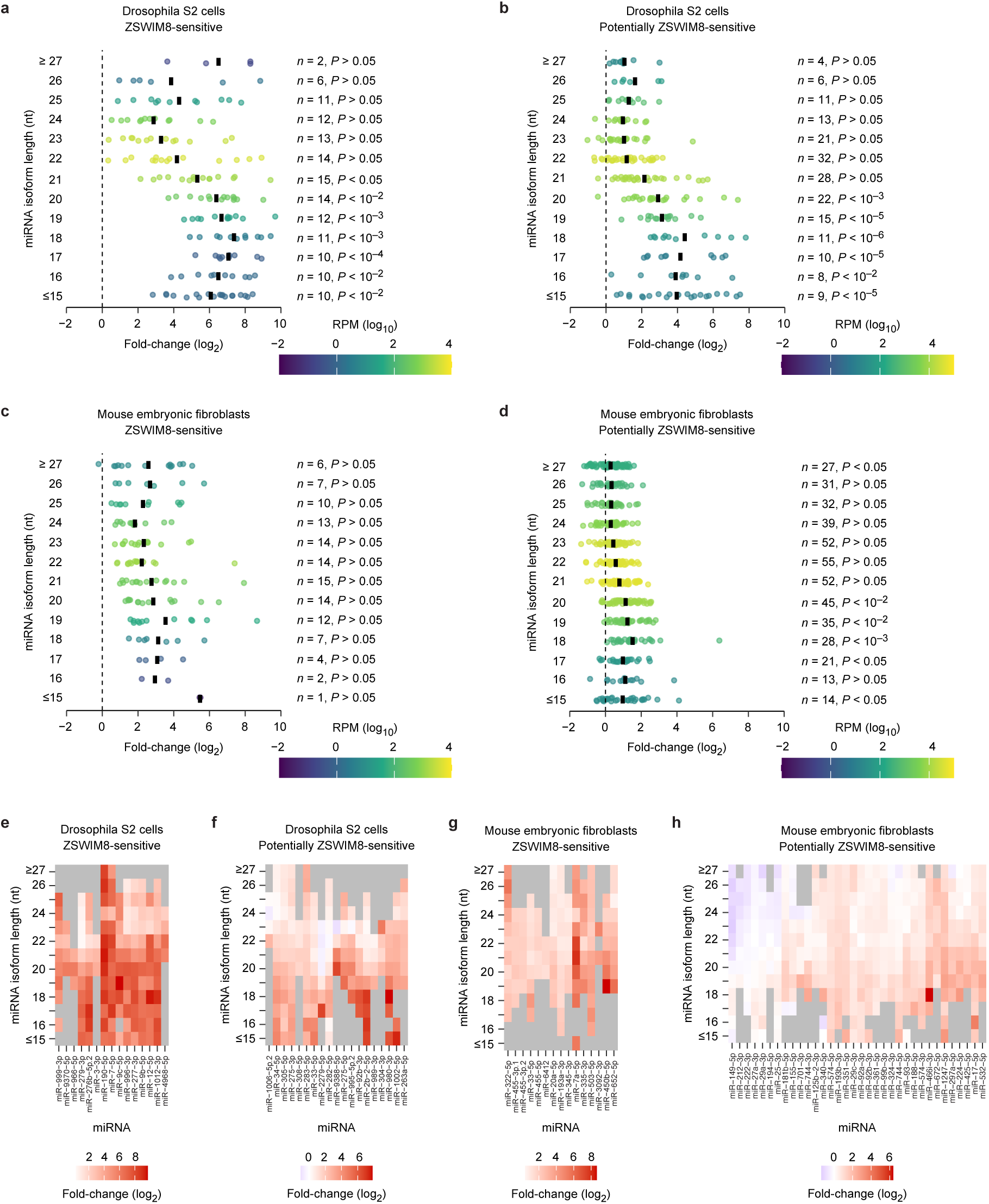
Relationship between miRNA isoform length and ZSWIM8 sensitivity for ZSWIM8-sensitive miRNAs. **a**, Distribution of fold-changes (log_2_) of miRNA molecules of the indicated lengths in *Zswim8*-knockout versus control Drosophila S2 cells for ZSWIM8-sensitive miRNAs. This panel is as in Extended Data Figure 8a, except it considers miRNAs classified as accumulating significantly upon loss of ZSWIM8 by BBUM analysis (FDR-adjusted *P*-value < 0.05)^91^. Each point represents a unique miRNA isoform of a given length, and each vertical black line represents the mean of the distribution of miRNA isoforms of the indicated length. **b**, Distribution of fold-changes (log_2_) of miRNA molecules of the indicated lengths in *Zswim8*-knockout versus control Drosophila S2 cells for potentially ZSWIM8-sensitive miRNAs. This panel is as in **a**, except it considers miRNAs that fail to meet the more stringent cutoff for ZSWIM8 sensitivity in **a** and are instead classified as potentially ZSWIM8-sensitive based on meeting one of three criteria: 1) a log_2_ fold-change > 0 and *P*_adj_ < 0.05, 2) a log_2_ fold-change significantly larger than that of their passenger strands, or 3) previous annotation as ZSWIM8-sensitive in S2 cells or embryos^17,23^. **c**, Distribution of fold-changes (log_2_) of miRNA molecules of the indicated lengths in *Zswim8*-knockout versus control MEFs for ZSWIM8-sensitive miRNAs. This panel is as in **a**, except the analysis is of data from MEFs. **d**, Distribution of fold-changes (log_2_) of miRNA molecules of the indicated lengths in *Zswim8*-knockout versus control MEFs for potentially ZSWIM8-sensitive miRNAs. This panel is as in **b**, except the analysis is of data from MEFs, and miRNAs were classified according to previous annotations of ZSWIM8 sensitivity in mouse tissues^24^. **e**, Relationship between miRNA isoform length and ZSWIM8 sensitivity in Drosophila cells for ZSWIM8-sensitive miRNAs. This panel is as in Extended Data Figure 8c, except it considers miRNAs classified as ZSWIM8-sensitive, as in panel **a**. All ZSWIM8-sensitive miRNAs were included in the analysis, regardless of the number of isoforms passing the expression cutoff. **f**, Relationship between miRNA isoform length and ZSWIM8 sensitivity in Drosophila cells for potentially ZSWIM8-sensitive miRNAs. This panel is as in **e**, except it considers miRNAs classified as potentially ZSWIM8-sensitive, as in panel **b.** Results for all potentially ZSWIM8-sensitive miRNAs with at least four isoforms of different lengths are shown. **g**, Relationship between miRNA isoform length and ZSWIM8 sensitivity in MEFs for ZSWIM8-sensitive miRNAs. This panel is as in **e**, except the analysis is of data from MEFs. All ZSWIM8-sensitive miRNAs were included in the analysis, regardless of the number of isoforms passing the expression cutoff. **h**, Relationship between miRNA isoform length and ZSWIM8 sensitivity in MEFs for potentially ZSWIM8-sensitive miRNAs. This panel is as in **f**, except the analysis is of data from MEFs. Results for all potentially ZSWIM8-sensitive miRNAs with at least six isoforms of different lengths are shown.

