## Supplementary material for "The E3 ubiquitin ligase mechanism specifying target-directed microRNA degradation": Methods

### Plasmids

Cloning was performed either by Gibson assembly with the NEBuilder HiFi DNA Assembly Master Mix (New England Biolabs (NEB), E2621) or by site-directed mutagenesis by direct PCR with the KAPA HiFi HotStart Ready Mix (Roche, 07958935001) or Q5 Master Mix (NEB, M0492S). Plasmids were purified using the Plasmid Plus Midi Kit (QIAGEN, 12945) and verified by Sanger or nanopore sequencing.

### Cell culture

All mammalian cells were cultured at 37 °C with 5% $CO_2$. Expi293F cells (Gibco, A14527) were cultured in Expi293 expression media (Gibco, A1435102) and passaged 1:10 every 3 days to maintain a maximal cell density of 3 million cells/mL. K562 cells harboring a miR-7-sensitive GFP reporter were a gift from Joshua Mendell[1]. K562 cells were cultured in RPMI 1640 (Gibco, 11875119) supplemented with 10% (v/v) fetal bovine serum (FBS) (Takara Bio, 631367) and passaged 1:4 every 3 days to maintain a maximal cell density of 800,000 cells/mL. Mouse embryonic fibroblast cells (MEFs) and HEK293FT cells were cultured in DMEM (Corning, 17-207-CV) with 10% FBS. Except in the case of contact inhibition for AGO2 intracellular co-IP and sRNA-seq experiments, described below, MEFs were passaged 1:4 or 1:8 as needed to maintain a maximal confluency of approximately 70%. HEK293FT cells were passaged 1:4 or 1:8 as needed to maintain a maximal confluency of approximately 70%. For passaging, adherent cells were dissociated using 0.25% trypsin–EDTA (Gibco, 25200114). For Expi293F cells, transfections were performed as described in the section "AGO–miRNA complexes for *in vitro* co-IP assays." For K562 and HEK293FT cells, transfections were performed with Lipofectamine 2000 (Invitrogen, 11668019) and Opti-MEM (Gibco, 31985062) according to the manufacturer's instructions. Drosophila S2 cells were cultured at 26 °C in Schneider's Drosophila Medium (Gibco, 21720001) supplemented with 10% heat-inactivated FBS (Gibco, 16140071). S2 cells were passaged every 3–6 days to maintain an approximate concentration between 1–10 million cells/mL.

### Proteins and RNAs

#### Proteins for *in vitro* ubiquitylation assays and for cryo-EM

Unless otherwise stated, all proteins are of human origin. Sequences of proteins and associated tags are listed in Supplementary Table 4. Genes were synthesized by TwistBiosciences or obtained as stated. Ubiquitin (UB) was expressed untagged and purified as described[2]. Cys-UB, where UB carries an N-terminal cysteine for protein labelling, and NEDD8, UBE2L3, UBE2R2, UBE2M, ARIH1, and ARIH2 were expressed as GST-TEV or GST-3C fusions in BL21(DE3) RIL bacterial cells, as described[3–6]. Following glutathione affinity pull-down (Cytiva, 17075605) and TEV protease cleavage, the protein was further purified by ion-exchange and size-exclusion chromatography. SrtA-5M, CUL3(NTD), CUL3(NTD, 1–384, I342R, L346D)-Avi, and CUL3(ΔN, NTD, 25–384, I342R, L346D)-Avi were expressed in BL21(DE3) RIL cells and purified by immobilized metal-affinity chromatography (Ni-INDIGO, Cube Biotech, 75105), followed by ion-exchange chromatography and size-exclusion chromatography, as described[7,8]. CUL2, CUL3, CUL5, GST-TEV-RBX1(5–108), GST-TEV-RBX2(5–113), and CUL2(NTD, 1–380, L344K, V3776D)-Avi were expressed in *Trichoplusia ni* High-Five insect cells using two baculoviruses for assembly of CUL–RBX complexes and purified following reported procedures[3,4]. CUL2–RBX1, CUL3–RBX1, and CUL5–RBX2 were neddylated and purified as described[3,4].

ZSWIM8-Twin-Strep was cloned into pFLN. ELOB and ELOC were cloned into pFLN and assembled into pBig1a using biGBac Assembly[9]. ZSWIM8–ELOB–ELOC complex (hereafter referred to as ZSWIM8–ELOB/C) was expressed in *Trichoplusia ni* High-Five insect cells using two baculoviruses for three days at 27 ºC. Cells were resuspended in lysis buffer (50 mM HEPES, 300 mM NaCl, and 10% (v/v) glycerol, pH 7.5) supplemented with protease inhibitors and 1 mM TCEP. Cells were disrupted by sonication and cell debris was removed by centrifugation. ZSWIM8–ELOB/C was enriched using Twin-Strep affinity purification (Strep-Tactin, IBA Lifesciences, 2-1201-025) and eluted with lysis buffer supplemented with 3 mM desthiobiotin, followed by size-exclusion chromatography (HiLoad Superdex 200, Cytiva).

ZSWIM8$^{mono}$ was designed based on our cryo-EM structure. We hypothesized that CUL3 binding to ZSWIM8 requires the presence of a folded D-domain. To functionally separate the requirement of ZSWIM8 dimerization for AGO binding and ubiquitylation from its impact on CUL3 binding, we designed a ZSWIM8 variant that retained a folded D-domain and retained binding to CUL3. We expected that transplanting a second copy of the D-domain into the native sequence would allow the D-domain to fold in *cis* and resemble the D-domain that is natively folded in *trans*. We inserted D-domain residues E222–P273 with two flanking GGSGGS linkers in between D-domain residues Q228 and R229 (Supplementary Table 3). AlphaFold3 structure predictions supported the expected assembly of the D-domain. This ZSWIM8 variant enabled us to functionally separate the involvement of ZSWIM8 dimerization in AGO binding and ubiquitylation from its role in CUL3 binding.

For structural studies and stepwise ubiquitin transfer assays, $His_6$-TEV-AGO2 was cloned into pFLN and expressed in *Trichoplusia ni* High-Five insect cells. AGO was purified following an adapted purification protocol[10]. In brief, insect cells were resuspended in 50 mM Tris, 100 mM KCl, and 5 mM DTT, pH 8.0. Following sonication, PEI was added to a concentration of 0.2% (v/v), mixed by inversion, and immediately centrifuged to remove cell debris. AGO was enriched by immobilized metal-affinity chromatography (Ni-INDIGO, Cube Biotech, 75105). Following affinity chromatography, a nucleic-acid-free AGO fraction (as judged by 260/280 ratio) was isolated by cation-exchange chromatography (Source 15S or MonoS, Cytiva) with a gradient of 0–500 mM KCl in 20 mM Tris, pH 8.0. Nucleic-acid-free AGO was incubated with synthetic single-stranded miRNA (Metabion; IDT; Supplementary Table 5) at room temperature for 30 min. Excess miRNA was removed by size-exclusion chromatography in 25 mM HEPES, 150 mM NaCl, and 1 mM TCEP, pH 7.8 (Superdex 200, Cytiva).

To label AGO, we used a split-intein-mediated labelling approach because of its high efficiency and site selectivity. We used the split intein gp41-1, which consists of an N-terminal part, gp41-1-N, and a C-terminal part, gp41-1-C[11,12]. These two parts excise themselves upon trans-splicing, leaving a minimal ligation scar on the protein consisting of two short extein motifs. gp41-1-N was cloned into pGEX downstream of a TEV cleavage site. Following the TEV cleavage site, an N-terminal GGG motif was incorporated for sortase labelling, along with the native extein sequence SGY. GST-gp41-1-N was expressed in Rosetta *E. coli* by induction with IPTG. Following glutathione-affinity purification and TEV protease cleavage, the protein was further purified by size-exclusion chromatography. gp41-1-C was cloned downstream of a Twin-Strep site into pFLN to create Twin-Strep-gp41-1-C-AGO2 or Twin-Strep-gp41-1-C-AGO1. These constructs contained the native extein linker, SSS, inserted between the split intein and the AGO sequence. AGO was purified following the procedure described above. These AGO variants were used for all *in vitro* polyubiquitylation assays and for *in vitro* co-IP assays using AGO2 loaded with a 14-nt miR-7.

**Labelling and biotinylation of proteins**

Peptides for sortase labelling were obtained from the Peptide Synthesis Facility (Max-Planck Institute of Biochemistry Core Facility). The sortase peptide had the following sequence: GSGGLPETGG, with an N-terminally linked carboxyfluorescein. Peptides were >90% pure as judged by HPLC and mass spectrometry.

gp41-1-N was fluorescently labelled at its N-terminal GGG using sortylation with a fluorescein-labelled acceptor peptide (fluorescein-GSGGLPETGG) (Supplementary Table 4). gp41-1-N (100 µM) was mixed with peptide (350 µM) in 25 mM HEPES, 150 mM NaCl, and 10 mM $CaCl_2$, pH 7.5. The reaction was initiated by addition of SrtA-5M (10 µM)[13] and allowed to proceed for 1 h at room temperature. Ni-INDIGO beads (20 µL) (Cube Biotech, 75105) were added and excess peptide was removed using a PD-MidiTRAP G-25 (Cytiva) followed by size-exclusion chromatography (Superdex 75) in 25 mM HEPES, 150 mM NaCl, and 1 mM TCEP, pH 7.5.

For labeling AGO proteins with fluorescein-gp41-1-N, the nucleic-acid-free fraction containing Twin-Strep-gp41-1-C-AGO2 or Twin-Strep-gp41-1-C-AGO1, isolated through ion-exchange chromatography, was incubated with miRNA and fluorescein-gp41-1-N at 1.2X molar excess over AGO. The reaction was allowed to proceed for 1 h at room temperature. Excess miRNA and unreacted fluorescein-gp41-1-N were removed by size-exclusion chromatography in 25 mM HEPES, 150 mM NaCl, and 1 mM TCEP, pH 7.5. Following labelling, AGO is modified at its N terminus with fluorescein-GGGSGYSSS (Supplementary Table 4). Fluorescently labelled AGO was used in all *in vitro* polyubiquitylation assays, as described below.

For biotinylation of CUL-NTDs, Avi-tagged CUL-NTDs were incubated with 50 mM ATP, 25 mM $MgCl_2$, 5 mM biotin, and 0.5 mg/mL BirA (produced in-house (22227598)), at 4 ºC for 18 h following ion-exchange chromatography. Biotinylated NTDs were purified by size-exclusion chromatography.

For fluorescent labelling of ubiquitin, Cys-UB (1 mM) was buffer-exchanged using a PD MidiTrap G-25 column (Cytiva, 28918008) into 25 mM HEPES, 150 mM NaCl, and 0.5 mM TCEP and incubated with 5-(Iodoacetamido)fluorescein (5 mM, Sigma-Aldrich, I9271) for 18 h at 4 ºC. Excess 5-

(Iodoacetamido)fluorescein was removed using a PD MidiTrap G-25 column followed by size-exclusion chromatography (Superdex 75) in 25 mM HEPES and 150 mM NaCl to yield fluorescently labelled UB (UB*). UB* was used in stepwise ubiquitin transfer assays, as described below.

**AGO–miRNA complexes for *in vitro* co-IP assays**

Plasmids expressing AGO2 variants were generated from the pcDNA3.3−3xFLAG−SUMO$^{Eu1}$−HsAGO2 plasmid (Addgene, 231372). The plasmid expressing AGO1 was subcloned into the same backbone from an existing AGO1-expressing plasmid. Sequences of proteins and associated tags are listed in Supplementary Table 4. AGO–miRNA complexes were prepared largely as described[14,15]. Guide and passenger RNAs were each chemically synthesized (IDT) and gel-purified on a urea-polyacrylamide gel, then annealed into duplexes at 5 µM final concentration in annealing buffer (30 mM Tris-HCl pH 7.5, 100 mM NaCl, and 1 mM EDTA). Annealing reactions were heated to 90 °C and slowly cooled to 30 °C over 1.5 h, before chilling on ice and storage at –80 °C. To generate cell lysate overexpressing the appropriate AGO variant, we transfected 200 mL of Expi293F cells at ~2 million/mL cell density, with 190 µg of the AGO expression plasmid, 10 µg of pMaxGFP (Lonza), and 600 µg of polyethylenimine (Polysciences, 23966) incubated in 10 mL of Opti-MEM at room temperature for 20 min. After culture for 40 h, cells were collected and lysed in 8 mL of hypotonic buffer (10 mM HEPES pH 8.0, 10 mM KOAc, 1.5 mM $Mg(OAc)_2$, 2% (v/v) glycerol, 5 mM NDSB-256, and 0.5 mM TCEP, with 1 tablet of cOmplete Mini EDTA-free Protease Inhibitor Cocktail (Roche, 11873580001) per 10 mL buffer) by a dounce homogenizer. Cell lysate was clarified by centrifugation at 3,000*g* for 15 min at 4 °C, then again for 30 min. The supernatant was re-equilibrated by adding 25% volume of re-equilibration buffer (150 mM HEPES pH 8.0, 700 mM KOAc, 15 mM $Mg(OAc)_2$, 42% (v/v) glycerol, 5 mM NDSB-256, and 0.5 mM TCEP, with 1 tablet of cOmplete Mini EDTA-free Protease Inhibitor Cocktail per 10 mL buffer), then further clarified by centrifugation at 60,000*g* for 20 min at 10 °C. Lysate was either used immediately or flash-frozen as aliquots in liquid nitrogen for storage at −150 °C.

To assemble each AGO–miRNA complex, 5–10 mL of cell lysate was first incubated with 600 µL of 5 µM annealed guide−passenger duplex for 1–2 h at 25 °C, to allow the in-lysate assembly of complexes. Meanwhile, 1 mL slurry of Dynabeads MyOne Streptavidin C1 (Invitrogen, 65002) was washed according to manufacturer's instructions, then pre-bound to 1 nmol of gel-purified biotinylated, 2′*O*-methylated capture oligonucleotide containing an 8mer seed-match site with complementarity to the loaded miRNA (Supplementary Table 5), at 25 °C for 30 min with end-over-end rotation, followed by four washes with ice-cold equilibration buffer (18 mM HEPES pH 7.4, 100 mM NaCl, 1 mM $MgCl_2$, 10% (v/v) glycerol, 0.003% (v/v) IGEPAL CA-630, 0.033 mg/mL recombinant albumin (NEB, B9200), and 0.005 mg/mL yeast tRNA (Life Technologies, AM7119)), then set aside at 4 °C until use. The incubated lysate was centrifuged at 6,000*g* for 10 min at 4 °C, filtered through a 5-µm PVDF filter (Millipore, SLSV025LS), and then added to the prepared beads and incubated at 25 °C for 1 h with end-over-end rotation. Beads were then washed twice with 5 mL equilibration buffer and twice with 5 mL capture-wash buffer (18 mM HEPES pH

7.4, 2 M NaCl, 1 mM $MgCl_2$, 10% (v/v) glycerol, 0.003% (v/v) IGEPAL CA-630, 0.033 mg/mL recombinant albumin, and 0.005 mg/mL yeast tRNA), then incubated with 1 nmol of 3′-biotinylated DNA competitor oligonucleotide (IDT) complementary to the capture oligonucleotide (Supplementary Table 5) in 18 mM HEPES pH 7.4, 1 M NaCl, 1 mM $MgCl_2$, 10% (v/v) glycerol, 0.003% (v/v) IGEPAL CA-630, 0.033 mg/mL recombinant albumin, and 0.005 mg/mL yeast tRNA, at 25 °C for 2 h with end-over-end rotation. The eluate was then incubated with 50 µL of anti-FLAG M2 magnetic bead slurry (Millipore, M8823) pre-equilibrated in equilibration buffer according to manufacturer's instructions, at 25 °C for 2 h with end-over-end rotation. The beads were washed twice with 500 µL of equilibration buffer and twice with 500 µL of storage buffer (18 mM HEPES pH 7.4, 100 mM NaCl, 1 mM $MgCl_2$, 5% (v/v) glycerol, and 0.003% (v/v) IGEPAL CA-630), before incubation in 100 µL of $SENP^{EuB}$ elution solution (500 nM of in-house purified $SENP^{EuB}$ protease[15] in storage buffer) at 4 °C with gentle rotation for 1 h. The eluate was combined with an additional 100-µL wash with $SENP^{EuB}$ elution solution, then supplemented with β-mercaptoethanol (BME) (Sigma-Aldrich, M6250) to a final concentration of 0.5 mM. Aliquots were flash-frozen in liquid nitrogen for storage at −80 °C. AGO2 loaded with a 14-nt miR-7 was purified using the same method as for purifying AGO2–miR-7 complexes for *in vitro* ubiquitylation assays, described above.

Purified AGO–miRNA complexes for use in *in vitro* co-IP assays were quantified by filter-binding titration. A serial dilution of limiting concentrations of complexes were incubated with 2 nM of a radiolabeled target RNA, in 18 mM HEPES pH 7.4, 100 mM NaCl, 1 mM $MgCl_2$, 2.5% (v/v) glycerol, 0.003% (v/v) IGEPAL CA-630, 0.5 mM BME, 0.017 mg/mL recombinant albumin, and 0.003 mg/mL yeast tRNA, at 37 °C for 1 h. Filter binding was conducted as described above, except membranes were pre-equilibrated in 18 mM HEPES pH 7.4, 100 mM NaCl, and 1 mM $MgCl_2$ and washed with 18 mM HEPES pH 7.4, 100 mM NaCl, 1 mM $MgCl_2$, and 5 mM DTT. Quantified fractions of target RNA bound across AGO–miRNA dilutions were fit to a quadratic equation by nonlinear least-squares regression in R using the Levenberg-Marquardt algorithm (nlsLM from the R package minpack.lm):

$$F_{bound} = \frac{\left([\text{stock}] \cdot DF + [\text{target}_T] + K_D\right) - \sqrt{\left([\text{stock}] \cdot DF + [\text{target}_T] + K_D\right)^2 - 4 \cdot [\text{stock}] \cdot DF \cdot [\text{target}_T]}}{2 \cdot [\text{target}_T]} \cdot F_{\max}$$

where $F_{\text{bound}}$ represents fraction of target bound, [$\text{target}_T$] represents the concentration of total target oligonucleotide, [stock] represents stock concentration of RISC, $DF$ represents the dilution factor, $K_D$ represents the dissociation constant for the affinity between RISC and the target, and $F_{\max}$ represents the maximal fraction of target bound at the plateau. A range of bounds and initiation values were tested for [stock], $K_D$, and $F_{\max}$ to ensure robust estimation of [stock]. $F_{\max}$ was always limited to the range (0, 1). $K_D$ was fit in log-transformed space.

**ZSWIM8 for *in vitro* co-IP assays**

3xFLAG-$SUMO^{Eu1}$-ZSWIM8-3xHA was cloned into pDARMO_CMVT (a gift from Kacper Rogala)[16]. ELOB and ELOC were cloned into pRK5. Sequences of proteins and associated tags are listed in

Supplementary Table 4. For 200 mL of culture, Expi293F cells at ~3 million/mL cell density were transfected with 95 µg of the ZSWIM8 expression plasmid, 47.5 µg of the ELOB expression plasmid, 47.5 µg of the ELOC expression plasmid, 10 µg of pMaxGFP, and 600 µg of polyethylenimine incubated in 24 mL of Opti-MEM at room temperature for 20 min. After culture for 48 h, cells were pelleted, washed with PBS, and resuspended in double the cell pellet volume of lysis buffer (18 mM HEPES pH 7.4, 150 mM KOAc, 10 µM $Zn(OAc)_2$, 5% (v/v) glycerol, 0.5% (v/v) IGEPAL CA-630, and 0.5 mM TCEP, with 1 tablet of cOmplete Mini EDTA-free Protease Inhibitor Cocktail per 10 mL buffer). The resuspension was passed through a 23G 1M 1.5′ needle and incubated at 4 °C for 15 min with end-over-end rotation. Lysate was clarified by centrifugation at 1,000*g* for 2 min at 4 °C, followed by centrifugation at 40,000*g* for 30 min at 4 °C. Meanwhile, 1.2 mL of anti-FLAG magnetic beads (Genscript, L00835) were washed twice with wash buffer (18 mM HEPES pH 7.4, 150 mM KOAc, 5% (v/v) glycerol, 0.01% (v/v) IGEPAL CA-630, and 0.5 mM TCEP). Clarified lysate was filtered through a 5-µm PVDF filter, then incubated with the prepared beads at 4 °C for 1.5 h with end-over-end rotation. After incubation with lysate, beads were washed four times with 3 mL wash buffer and eluted with 1 mL of elution buffer (5 µM of in-house purified $SENP^{EuB}$ protease[15] in wash buffer) at 4 °C for 1.5 h with end-over-end rotation. The eluate was centrifuged at 17,000*g* for 5 min at 4 °C, and the supernatant was collected and quantified by nanodrop. Aliquots were flash-frozen in liquid nitrogen for storage at −80 °C.

**T6B peptide**

3xHA-tagged T6B peptide (6xHis-SUMO-TNRC6B(599–683)-3xHA[17], otherwise referred to as T6B-3xHA peptide) was expressed and purified as described previously[18], with some modifications. 3xFLAG-tagged T6B peptide (6xHis-SUMO-TNRC6B(599–683)-3xFLAG, otherwise referred to as T6B-3xFLAG peptide) used in sRNA-seq experiments, described below, was purified by an analogous method. Sequences of proteins and associated tags are listed in Supplementary Table 4.

6xHis-SUMO-TNRC6B(599–683)-3xHA was cloned into pET28a. The expression construct was transformed into BL21 (DE3) competent *E. coli* (NEB, C2527H). A single transformant was used to seed an overnight culture in LB supplemented with 50 µg/mL kanamycin (GoldBio, K-120-5) and 35 µg/mL chloramphenicol (GoldBio, C-105-25) and grown at 30 °C overnight with shaking. Each 1.5-L batch of LB was supplemented with 50 µg/mL kanamycin and 35 µg/mL chloramphenicol and seeded with 15 mL of overnight culture. This culture was grown at 37 °C with shaking. Once the culture reached an $OD_{600}$ of ~0.2, it was transferred to 18 °C with shaking, and once it reached an $OD_{600}$ of ~0.6, 0.5 mM of IPTG (GoldBio, I2481C50) was added. Induced cells were grown further overnight at 18 °C with shaking.

Cells were collected by centrifuging at 8,980*g* for 6 min at 4 °C. Pellets were resuspended in lysis buffer (50 mM Tris pH 8.0, 500 mM NaCl, 5% (v/v) glycerol, 15 mM imidazole, 4 mM BME, and 1 tablet of cOmplete EDTA-free Protease Inhibitor Cocktail (Roche, 11873580001)), with 1 mL lysis buffer per 240 mL of overnight culture. Cells were lysed by sonication, and lysate was clarified by centrifuging at 41,400*g* for 30 min at 4 °C. Clarified lysate was added to Ni-NTA agarose beads (QIAGEN, 30210) that had been washed twice with lysis buffer (280 µL of slurry per 1 L of culture), and the slurry was incubated at 4 °C for 2 h with end-over-end rotation. The beads were collected by centrifugation at 100*g* for 2–5 min at 4 °C and washed three times with 40 bead volumes of lysis buffer. Following the last wash, beads were resuspended in 5 mL of lysis buffer and transferred to a gravity-flow column. Captured protein was eluted with elution buffer (50 mM Tris pH 8.0, 500 mM NaCl, 5% (v/v) glycerol, 250 mM imidazole, and 4 mM BME).

The eluate was dialyzed at 4 °C overnight in dialysis buffer (50 mM Tris pH 8.0, 100 mM NaCl, 5% (v/v) glycerol, 15 mM imidazole, and 4 mM BME) using a 10-kDa MW cutoff dialysis cassette (Thermo Scientific, 69570). On the following day, the dialyzed eluate was purified by anion-exchange chromatography on either a Resource Q column (Cytiva, 17-1179-01) or a HiTrap Q XL column (Cytiva, 17515901) pre-equilibrated in buffer A (50 mM Tris pH 8.0, 100 mM NaCl, 5% (v/v) glycerol, and 4 mM BME). T6B-3xHA peptide was eluted off the column with a linear gradient of buffer B (50 mM Tris pH 8.0, 2 M NaCl, 5% (v/v) glycerol, and 4 mM BME). Fractions containing T6B-3xHA peptide were pooled and concentrated by centrifugal filtration using a concentrator pre-equilibrated with buffer A (Amicon Ultra 10 kDa cutoff, Millipore, UFC8010). Concentrated T6B-3xHA peptide was further purified using size-exclusion chromatography with a Superdex 200 Increase column (Cytiva, 28990944) in buffer A. Fractions containing T6B-3xHA peptide were pooled, then centrifuged at 17,000*g* for 5 min at 4 °C, and the supernatant was collected and quantified by nanodrop. Aliquots were flash-frozen in liquid nitrogen for storage at −80 °C.

**Target RNAs**

Shorter target RNAs were chemically synthesized (IDT; Supplementary Tables 1,5) with 5′ hydroxyl chemistry, and gel-purified on a urea-polyacrylamide gel. Longer target RNAs were *in vitro* transcribed using chemically synthesized and gel-purified single-stranded DNA templates annealed to an oligonucleotide containing the T7 promoter sequence (IDT; Supplementary Tables 1,5). Transcription reactions using in-house purified T7 RNA polymerase were conducted in 5 mM ATP, 2 mM UTP, 5 mM CTP, 8 mM GTP, 5 mM DTT, 40 mM Tris pH 7.9, 2.5 mM spermidine, 26 mM $MgCl_2$, 0.01% (v/v) Triton X-100, 5 mM DTT, SUPERase•In (1 U/µL; Invitrogen, AM2694), and Thermostable Inorganic Pyrophosphatase (0.0083 U/µL; NEB, M0296). After incubation at 37 °C for 3−4 h, DNA templates were digested with RQ1 DNase (Promega, M6101) at 37 °C for 30 min. RNA products were then gel-purified on a urea-polyacrylamide gel.

Gel-purified chemically synthesized RNAs were radiolabeled directly using T4 Polynucleotide Kinase (PNK) (NEB, M0201) and [γ-$^{32}$P] ATP (Revvity, BLU035C005MC) in T4 PNK reaction buffer (NEB, B0201S) at 37 °C for 1–2 h, then desalted using Micro Bio-Spin P-6 columns (Bio-Rad, 7326221) and gel-purified on a urea-polyacrylamide gel. *In vitro* transcription products were first dephosphorylated using Quick CIP (NEB, M0525) at 37 °C for 15 min, followed by heat-inactivation at 80 °C for 3 min. Dephosphorylated RNAs were then immediately radiolabeled using T4 PNK and [γ-$^{32}$P] ATP in rCutSmart buffer (NEB, B6004S) supplemented with 5 mM DTT (Invitrogen, 15508013) at 37 °C for 1–2 h, then desalted using Micro Bio-Spin P-30 columns (Bio-Rad, 7326250), Micro Bio-Spin P-6 columns, or Amersham MicroSpin G-25 columns (Cytiva, 27532501) and gel-purified on a urea-polyacrylamide gel.

#### Biochemical assays

##### *In vitro* polyubiquitylation assays

The identification of cullin–RING ligase subunits and the ARIH1 RBR-type E3 as TDMD effectors[1] suggested that ZSWIM8 performs ubiquitylation by the previously defined E3–E3 mechanism, which is a variation on the canonical E1–E2–E3 pathway[3,19]. In the E3–E3 mechanism, ubiquitin is transferred from the E2 UBE2L3 to the active site of ARIH1 only upon ARIH1 activation through binding a neddylated CRL. ARIH1 then ubiquitylates the substrate recruited to the CRL substrate receptor, and generates short ubiquitin chains. Optimal polyubiquitylation is achieved in collaboration with a UBE2R-family E2, also activated by the neddylated CRL. The design of our *in vitro* polyubiquitylation assays was informed by this mechanism.

All concentrations stated refer to the final concentrations used. Fluorescently labelled AGO2 (0.2 µM) loaded with the indicated miRNA was incubated with a target RNA (0.22 µM) for 15 min at room temperature. In the meantime, CUL-NEDD8–RBX1 (0.5 µM), ZSWIM8–ELOB/C (0.5 µM), ARIH1 (0.4 µM), UBE2L3 (1.5 µM), UBE2R2 (1.5 µM), UB (50 µM), BSA (0.5 mg/mL) were mixed in 50 mM HEPES, 50 mM NaCl, and 1 mM TCEP, pH 7.5. AGO–miRNA–target RNA complex was added to the remaining components and incubated for 1 h at room temperature. ATP (5 mM) and $MgCl_2$ (7.5 mM) were added, and the reaction was initiated by addition of UBA1 (0.1 µM). At indicated time points, samples were removed and quenched with an equal volume of 2X SDS-PAGE sample buffer (100 mM Tris-HCl, 20% (v/v) glycerol, 30 mM EDTA, 4% (v/v) SDS, and 45 BME) and incubated at 95 ºC for 5 min. Samples were resolved on 4–22% SDS-PAGE gels and visualized on a Typhoon 9410 Imager by in-gel fluorescence, followed by gel staining with Coomassie blue. Assays were performed with $n \geq 2$, and representative gels are shown. For reactions containing CUL5 and ARIH2, CUL-NEDD8–RBX1 was substituted for CUL5-NEDD8–RBX2, and ARIH1 was substituted with ARIH2. The assay setup as described above was used to generate the data shown in Figure 1c, d; 3f; Extended Data Figure 1e, h, i; 3m; 4i; 6f; 9a, e; 10c. Gels stained with Coomassie blue are provided in the Supplementary Figures.

##### Stepwise ubiquitin transfer assays

Stepwise transfer assays were performed in a two-step process consisting of a loading reaction and a discharge reaction. In the loading reaction, the UB-carrying enzyme (UBE2L3) is charged by UBA1, and quenched by apyrase, which prevents further UB activation once UBE2L3 is discharged. The discharge reaction is initiated by addition of the remaining reaction components (CRL, substrate receptor, substrate, and additional E3s). Stepwise ubiquitin transfer assays follow the transfer of ubiquitin not just to the substrate but also to and from other proteins such as E2 and E3 components. Substrate receptors that interact with CRLs are commonly autoubiquitylated and thus autoubiquitylation provides a functional readout of the ability of a substrate receptor to interact with a catalytically competent CRL.

All concentrations stated refer to the final concentrations used. AGO2 (0.2 µM) loaded with the indicated miRNA was incubated with a target RNA (0.22 µM) at room temperature for 15 min. In the meantime, CUL-NEDD8–RBX1 (0.5 µM), ZSWIM8–ELOB/C (0.5 µM), ARIH1 (0.4 µM), and BSA (0.5 mg/mL) were mixed in 50 mM HEPES, 50 mM NaCl, and 1 mM TCEP, pH 7.5, and incubated at room temperature for 15 min. In the meantime, UBE2L3 (10 µM) was mixed with fluorescein-UB (15 µM) in 25 mM HEPES, 100 mM NaCl, 2.5 mM $MgCl_2$, and 1 mM ATP, at room temperature. E2 charging was initiated by addition of UBA1 (0.4 µM) and allowed to proceed for 15 min. The reaction was quenched by addition of apyrase (NEB, 0.25 units) and incubated at 4 ºC for 5 min. Assuming complete charging of UBE2L3, UBE2L3~UB* was further diluted to 2 µM in 25 mM HEPES and 150 mM NaCl, pH 7.5. Ubiquitylation was initiated by addition of UBE2L3~UB to the other components at a final concentration of 0.4 µM. For the sample at $t_0$, an aliquot of UBE2L3~UB was diluted to 0.4 µM in 25 mM HEPES and 150 mM NaCl, pH 7.5, and a sample was immediately removed and quenched with 2X SDS-PAGE sample buffer (100 mM Tris-HCl, 20% glycerol, 30 mM EDTA, and 4% SDS). Samples were removed and quenched at indicated time points. Samples were resolved on 4–22% SDS-PAGE gels and visualized on a Typhoon 9410 Imager by in-gel fluorescence, followed by gel staining with Coomassie blue. Assays were performed with $n \geq 2$, and representative gels are shown. The assay setup as described above was used to generate the data shown in Extended Data Figure 3l; 4f. Gels stained with Coomassie blue and are provided in the Supplementary Figures.

***In vitro* co-IP assays**

All concentrations stated refer to the final concentrations used. Pierce anti-HA magnetic bead slurry (Thermo Scientific, 88837, 15 µL per reaction) was washed twice with blocking buffer (18 mM HEPES pH 7.4, 150 mM KOAc, 1 mM $Mg(OAc)_2$, 5% (v/v) glycerol, 0.01% (v/v) IGEPAL CA-630, 0.1 mg/mL recombinant albumin, and 0.01 mg/mL yeast tRNA) and incubated in blocking buffer at 4 °C for 2 h with end-over-end rotation. Blocking buffer was then removed immediately before use. Radiolabeled target RNAs were prepared as described above and pre-mixed. Pre-mixed RNA concentrations were calculated based on a conservative estimate of 0–50% loss during radiolabeling. AGO–miRNA complex was pre-diluted in storage buffer (18 mM HEPES pH 7.4, 100 mM KOAc, 1 mM $Mg(OAc)_2$, 20% (v/v) glycerol, 0.003% (v/v) IGEPAL CA-630, 0.5 mM BME, 0.033 mg/mL recombinant albumin, and 0.005 mg/mL yeast tRNA). Unless other concentrations are specified, 0.75 nM of pre-diluted AGO–miRNA complex was pre-incubated with ~0.25 nM target RNAs in co-IP buffer (18 mM HEPES pH 7.4, 150 mM KOAc, 2 mM $Mg(OAc)_2$, 5% (v/v) glycerol, 0.01% (v/v) IGEPAL CA-630, and 0.5 mM BME) supplemented with blocking reagents (0.1 mg/mL recombinant albumin and 0.01 mg/mL yeast tRNA) at 37 °C for 30–60 min. Where specified, heparin (Sigma-Aldrich, H3393), which reduces nonspecific binding of RNA (especially longer

RNA) to the beads, was also included in the blocking buffer and co-IP buffer. 1 μg/mL heparin was included in assays shown in Figure 1f; 3g; 4e; Extended Data Figure 7d; 10b. 10 μg/mL heparin was included in assays shown in Extended Data Figure 1c; 9d. Heparin was not included in assays shown in Figure 1e; Extended Data Figure 1d, g; 2a–d; 9f–h. In the meantime, bait proteins ZSWIM8-3xHA and T6B-3xHA were pre-diluted in ZSWIM8 buffer (18 mM HEPES pH 7.4, 150 mM KOAc, 5% (v/v) glycerol, 0.01% (v/v) IGEPAL CA-630, and 0.5 mM TCEP).

Pre-incubated AGO–miRNA–target complex and pre-diluted bait proteins (or ZSWIM8 buffer) were mixed at 1:1 ratio and incubated at 25 °C for 1 h, at a final volume of 30 μL per reaction. To reduce non-specific binding during the subsequent immunoprecipitation, any aggregation-prone species were removed by centrifugation at 17,000–21,000*g* for 5 min at 4 °C. 5% of the supernatant was taken from the no-bait control to serve as input. Supernatants were otherwise resuspended with pre-blocked beads and incubated at 4 °C for 1.5 h, with intermittent shaking at 1,200 RPM for 15 s every 2 min. Following incubation, reactions were transferred to fresh tubes, and beads were washed four times with 125 μL of 18 mM HEPES pH 7.4, 150 mM KOAc, 1 mM $Mg(OAc)_2$, 5% (v/v) glycerol, 0.01% (v/v) IGEPAL CA-630, and 0.5 mM BME, with a tube change during the fourth wash. Immunoprecipitated species were eluted by resuspension of beads with gel loading buffer (8 M urea, 25 mM EDTA, 0.025% (w./v.) xylene cyanol, 0.025% (w./v.) bromophenol blue) and shaking at 1,000 RPM at 65 °C for 10 min. Eluates were run on a urea-polyacrylamide gel. The gel was frozen at –20 °C while exposing a phosphorimager plate, and radioactivity was subsequently imaged on a phosporimager. Band intensities were quantified using ImageQuant TL (v10.2); image background was estimated either by subtracting the mean signal of equal-sized rectangles drawn at regions with no bands, or, where indicated, by additionally examining the lane profile tool drawn across each lane. In the assays shown in Extended Data Figure 7d and e, statistical analyses were performed in GraphPad Prism (v10.6.1), using an ordinary one-way ANOVA with Dunnett's multiple comparisons tests to compare each AGO2 variant to WT AGO2 (*$P < 0.05$, **$P < 0.001$, ***$P < 0.0001$; n.s., not significant). The assay setup as described above was used to generate the data shown in Figure 1e, f; 3g, 4e; Extended Data Figure 1c, d, g; 2a–d; 7d, e; 9d, f–h; 10b.

**Filter binding assays**

To measure intrinsic affinity of ZSWIM8 for RNAs, a dilution series of purified ZSWIM8-3xHA was incubated in excess with radiolabeled RNAs in 18 mM HEPES pH 7.4, 150 mM KOAc, 1 mM $Mg(OAc)_2$, 5% (v/v) glycerol, 0.01% (v/v) IGEPAL CA-630, 0.25 mM BME, 0.25 mM TCEP (Supelco, 646547), 0.1 mg/mL recombinant albumin, and 0.01 mg/mL yeast tRNA, at 25 °C for 2 h.

Nitrocellulose (Amersham Protran, 0.45 μm pores; Cytiva, 10600062) and nylon (Amersham Hybond-XL; Fisher Scientific, 45001147) membrane filters were cut with a circle punch into discs of 0.5-inch diameter each and pre-equilibrated at room temperature for at least 20 min in 18 mM HEPES pH 7.4, 100 mM KOAc, and 1 mM $Mg(OAc)_2$. For each sample, a nitrocellulose disc was stacked on top of a nylon disc, atop a circular pedestal mounted on a Visiprep SPE Vacuum Manifold (Supelco, 57250-U), set at approximately −20 kPa. 10 µL of reaction was promptly applied to the stacked discs, followed by 100 µL of ice-cold wash buffer (18 mM HEPES pH 7.4, 100 mM KOAc, 1 mM $Mg(OAc)_2$, and 5 mM DTT). Filter membrane discs were separated with tweezers, air-dried, then exposed on a phosphorimager plate. Radioactivity was imaged on a Typhoon phosporimager (Cytiva), and spot intensities were quantified using ImageQuant TL (v10.2). The assay setup as described above was used to generate the data shown in Extended Data Figure 2e.

**Analytical size-exclusion chromatography**

ZSWIM8–ELOB/C, $ZSWIM8^{mono}$–ELOB/C, and $ZSWIM8^{\Delta dimer}$–ELOB/C (10 µM) were each incubated with CUL3–RBX1 (12 µM) in 25 mM HEPES, 150 mM NaCl, and 1 mM TCEP, pH 7.5 at 4 ºC for 30 min prior to analysis by size-exclusion chromatography (Superose 6, Cytiva). Samples were resolved on 4–22% SDS-PAGE gels, and gels were stained with Sypro Ruby (Sigma-Aldrich, S4942) prior to analysis by in-gel fluorescence. Analytical size-exclusion chromatograms and corresponding SDS-PAGE analyses are shown in Extended Data Figure 3k.

**Bio-layer interferometry**

Bio-layer interferometry measurements were performed on an Octet K8 system (Sartorius) at 30 ºC with shaking at 1,000 RPM. Concentrated proteins were diluted into BLI buffer (25 mM HEPES, 150 mM NaCl, 0.5 mg/mL BSA, and 1 mM TCEP, pH 7.5). Biotinylated cullin-NTDs were immobilized on streptavidin sensors (Sartorius, 18-5019). ZSWIM8 served as the analyte and was measured in seven dilutions from 650 nM to 0.9 nM. Raw data were processed in Octet Data Analysis HT software. As no dissociation was detected, the $K_D$ was estimated through fitting $R_{max}$ of association. Processed data were plotted and fit using a single-site binding model in GraphPad Prism (v10.4.0).

**Sequence alignments of protein homologs**

Protein sequences were downloaded from UniProt[20]. Multiple-sequence alignment was conducted with the MUSCLE algorithm[21]. Maximum-likelihood phylogeny was calculated using FastTree 2.2[22].

**Cryo-EM**

**Sample preparation.** All concentrations stated refer to the final concentrations used. AGO2–miR-7 (7 µM) was mixed with a 120-nt fragment of the CYRANO trigger (8 µM) (Supplementary Table 1) and incubated at room temperature for 15 min. In the meantime, ZSWIM8–ELOB/C (6 µM) was mixed with CUL3(NTD) (6 µM) in 25 mM HEPES, 50 mM NaCl, and 1 mM TCEP, pH 7.8. AGO2–miR-7–CYRANO was added and incubated on ice until plunge freezing. Cu holey-carbon grids (R1.2/1.3, 200 mesh, Quantfoil, Q250-CR1.3) were freshly glow-discharged. 3.5 µL of complex was applied, incubated for 5 s and blotted with a blot force of 2 and blot time of 2 s using a Vitrobot Mark IV (4 ºC, 95% humidity) and plunge-frozen in liquid ethane.

**Electron microscopy.** A screening data set was recorded on a Glacios cryo-transmission electron microscope at 200 kV using a Gatan Vista Alpine detector in counting mode and SerialEM software. The dataset consisted of 3,998 movies with a pixel size of 1.871 Å. Total exposure was set to 60 electrons $Å^{-2}$ (40 frames), and a defocus range between –0.5 and –2.5 µm. A high-resolution dataset was recorded on a Titan Krios transmission electron microscope at 300 kV using a Gatan K3 detector in counting mode and Serial EM software. The dataset (26,355 micrographs) was recorded at a magnification of 105,000, pixel size 0.8512 Å, total exposure 58 electrons $Å^{-1}$, and a defocus range –0.5 and –2.0 µm. A representative micrograph can be found in Supplementary Figure 2.

**Data processing.** All datasets were processed in CryoSparc (v4.6.2)[23] (Supplementary Table 2). Raw movie files were motion-corrected and dose-weighted, followed by estimation of contrast-transfer functions. Exposures were manually curated to remove micrographs with damage or ice contamination. Particles were picked in CryoSparc without a template and with a template generated from the screening dataset. Heterogeneous refinements, 3D classification, global and local refinements, particle subtraction, and post-processing were performed in CryoSparc. For all heterogeneous refinements, one decoy class was included. All final maps were post-processed using manual sharpening or DeepEMhancer. The processing scheme is shown in Supplementary Figure 2 and 3. 3D variability analysis was performed in CryoSparc with six modes and a filter resolution of 10 Å.

**Model building.** The high-resolution map of CUL3(NTD)–ZSWIM8–ELOB/C–AGO2–miR-7–CYRANO (map A, sharpened with a B factor of 89.8) was further refined using three focused-refined maps showing high-resolution features (maps B, C, and D). A composite map was constructed based on focused-refined and DeepEMhancer-sharpened maps (map E). Each focused map has been deposited, along with a composite map. Maps A, B, C, and E were used to build $ZSWIM8^{NPAZ}$ and $ZSWIM8^{MID}$ interactions with AGO2, RNA, and C-terminal HEAT repeats. Maps A, C, D, and E were used to build CUL3(NTD), ELOB, ELOC, and the ZSWIM8 E3 superdomain. Structurally flexible RNA-binding elements (RBEs) were modelled using a combination of unsharpened maps (map F) and low-pass filtered maps (map G). An initial model was generated by docking *de novo*-folded ZSWIM8 (AlphaFold3), CUL3 (PDB 5NLB), ELOB/C (PDB 1LM8), and AGO2 (PDB 6NIT) into the composite map. Iterative manual model building in COOT and real-space refinement using Phenix.refine were performed until a satisfactory map-to-model correlation was achieved[24,25]. Due to the structural heterogeneity of CUL3, only the first 150 residues were modelled. For ZSWIM8, side chains were built for residues with interpretable density. For CUL3 and ELOB/C, side-chain residues were included based on input models. Three long unstructured regions in ZSWIM8 were not modelled due to the absence of any interpretable density. The TDMD sensor domain of $ZSWIM8^{NPAZ}$ includes a disconnected density that was modelled to contain ZSWIM8 residues Q39–R45 based on AlphaFold3 predictions. RBEs 1–3 on $ZSWIM8^{NPAZ}$ were modelled with side chains based on sharpened and unsharpened maps. RBEs 1–3 on $ZSWIM8^{MID}$ were modelled without side chains due to inferior densities. C-terminal HEAT repeats were modelled based on AlphaFold3 predictions without side chains due to a lack of interpretable side-chain densities. Trigger RNA base U44 was poorly resolved in sharpened maps; its flipped-out position was modeled based on unsharpened maps. The density for trigger RNA base A43 indicates a syn conformation and was modelled as such. All figures were generated using ChimeraX (v1.8–v.1.9)[26]. All densities shown in figures were prepared using the composite map, unless otherwise stated.

**Cellular assays**

**Cell lines**

For generating clonal *ZSWIM8*-knockout K562s, parental K562 cells harboring a miR-7-sensitive GFP reporter[1] were transiently transfected with a plasmid derived from PX458 (Addgene #48138) expressing pSpCas9-2a-BFP and either one or two sgRNAs targeting the *ZSWIM8* gene (Supplementary Table 5), followed by single-cell sorting of BFP-positive cells. Clones were screened for disruption of all *ZSWIM8* alleles by amplification of the locus followed by nanopore sequencing of the PCR amplicon.

When generating clonal *Zswim8*-knockout MEFs, we observed substantial heterogeneity in clonal MEF populations derived from polyclonal parental populations, which was consistent with previous observations[27]. To minimize this clone-to-clone variability, we first generated a clonal parental population of wild-type MEFs by single-cell sorting of GFP-positive cells from MEFs[28] that had been transiently transfected with a plasmid expressing pSpCas9-2a-GFP and a non-targeting sgRNA (Addgene #48138, Supplementary Table 5). After isolating clonal lines, these parental cells were transiently transfected with a plasmid expressing pSpCas9-2a-GFP and an sgRNA targeting the *Zswim8* gene (Addgene #48138, Supplementary Table 5) followed by single-cell sorting of GFP-positive cells. Clones were screened for disruption of all *Zswim8* alleles by amplification of the locus followed by Sanger sequencing of the PCR amplicon. Clonal wild-type MEFs were generated in parallel from the same parental clonal lines, except using a plasmid expressing pSpCas9-2a-GFP and a non-targeting sgRNA (Supplementary Table 5).

Polyclonal knock-in K562 lines stably expressing doxycycline-inducible *ZSWIM8* transgenes were generated by PiggyBac transposition into clonal *ZSWIM8*-knockout K562 cells containing a miR-7-sensitive GFP reporter, described above. The wild-type ZSWIM8 CDS was cloned into PB-TetON (Addgene #97421), and ZSWIM8 variants were generated from this wild-type plasmid. For each well in a 6-well plate, ~600,000 cells were transfected with 1 µg of the PiggyBac transposon plasmid and 0.4 µg of the Super PiggyBac transposase plasmid (a gift from Anders Hansen) using Lipofectamine 2000 and Opti-MEM according to the manufacturer's instructions. After 48 h, cells were passaged 1:2 into fresh media containing 3 µg/mL puromycin (Gibco, A1113803). Subsequently, selection media was refreshed every three days as cells were expanded up to 10-cm plates.

**ZSWIM8 intracellular rescue assay**

Following generation of *ZSWIM8*-knockout reporter K562 cells stably expressing doxycycline-inducible *ZSWIM8* transgenes, as described above, ~400,000 cells were plated per well in a 6-well plate with media containing 3 µg/mL puromycin and 33.3 ng/mL doxycycline (Takara Bio, 631311). After 48 h, cells were expanded to 10-cm plates and media containing 3 µg/mL puromycin and 33.3 ng/mL doxycycline was refreshed. After another 48 hr, ~75% of the culture was set aside for northern analysis and ~12.5% was set aside for immunoblotting. The remaining ~12.5% was concentrated to 2 million cells/mL in media and subjected to flow cytometry. For each sample, ~20,000 live cells were analyzed using a BD LSR Fortessa instrument (BD Biosciences). Subsequent analysis was performed using FlowJo (v10.10.0). Gating strategies for flow cytometry analysis are shown in Supplementary Figure 5. Statistical analyses were performed in GraphPad Prism (v10.6.1), using an ordinary one-way ANOVA with Dunnett's multiple comparisons tests to compare each ZSWIM8 variant to WT ZSWIM8 (*$P < 0.05$, ***$P < 0.0001$; n.s., not significant). Quantified results of the ZSWIM8 intracellular rescue assay are shown in Figure 3c; 4c; Extended Data Figure 3f, n; 4b, j. Immunoblots of ZSWIM8 protein variants are shown in Supplementary Figure 6.

**RNA isolation**

Total RNA was extracted with TRI Reagent (Invitrogen, AM9738) according to the manufacturer's instructions. K562 cells were pelleted by centrifugation, washed with PBS, and resuspended in 1 mL of TRI Reagent. GlycoBlue (Invitrogen, AM9516) was added to a final 25 µg/mL to facilitate precipitation and visualization of RNA pellets. After isopropanol precipitation and one 75% ethanol wash, pellets were resuspended in water and quantified by nanodrop.

**Northern blots**

Unless otherwise specified, 10 µg of total RNA was resolved on a denaturing 15% polyacrylamide gel and transferred onto a Hybond-N+ membrane (Cytiva, RPN303B) using a semi-dry transfer apparatus (Bio-Rad). The membrane was then incubated at 65 °C for 1–2 h with EDC (N-(3-dimethylaminopropyl)-N′-ethylcarbodiimide; Thermo Scientific, 22980) diluted in 1-methylimidazole, which chemically crosslinked 5′ phosphates to the membrane. Blots were hybridized to radiolabeled oligonucleotide probes (DNA or LNA) (Supplementary Table 5). Stripping of probes prior to re-probing was performed by incubation in boiling 0.04% (v/v) SDS with shaking. A detailed protocol for northern-blot analysis of small RNAs is available at http://bartellab.wi.mit.edu/protocols.html. Results were analyzed on a Typhoon phosphorimager, using ImageQuant TL (v10.2) for quantification of band intensities.

**Immunoblots**

K562 cells were pelleted by centrifugation, washed with PBS, and lysed using Pierce IP Lysis Buffer containing 1 tablet of cOmplete Mini EDTA-free Protease Inhibitor Cocktail per 10 mL buffer. Following 15 min of incubation in lysis buffer at 4 °C, lysates were clarified by centrifugation at 17,000*g* for 15 min at 4 °C and quantified by Bradford assay. Clarified lysates were incubated at 65 °C for 10 min in the presence of 1X NuPAGE LDS sample buffer (Invitrogen, NP0007) and 2.5% (v/v) BME. 8–12 µg of lysate was resolved on NuPAGE 4–12% Bis-Tris protein gels (Invitrogen, WG1403) with 1X MES SDS running buffer (Invitrogen, NP0002) at 180 V for 50 min, and then transferred onto 0.45-µm PVDF membranes (Invitrogen, STM2006) at 25 V for 1 h on ice using a Midi Gel Tank (Invitrogen, STM1001) and Midi Blot Module (Invitrogen, STM2001). Membranes were then blocked with 5% (w./v.) BSA (RPI, A30075-100.0) dissolved in PBS containing 0.1% (v/v) Tween 20 (Sigma-Aldrich, P1379) (PBS-T) for 1 h at room temperature, and then incubated with primary antibodies overnight at 4 °C. The following day, membranes were washed with 0.1% PBS-T at room temperature and incubated with near-infrared fluorescent secondary antibodies (LI-COR) for 1 h at room temperature, followed by washes and detection on an Odyssey CLx imager (LI-COR). Primary antibodies and their dilutions included: rabbit polyclonal anti-ZSWIM8 (1:400; Invitrogen, PA5-59492), rabbit monoclonal anti-HA (1:5,000; Cell Signaling Technology, C29F4, 3724), and mouse monoclonal anti-GAPDH (1:2,000; Invitrogen, GA1R, MA5-15738). Near-infrared fluorescent secondary antibodies were used at 1:10,000 (LI-COR). Protein levels were quantified in ImageStudio (LI-COR, v6.1.0.79).

**Lentiviral transduction**

Lentiviral production and transduction were performed as described[29]. Briefly, transfer plasmids expressing wild-type AGO2 as well as R40 and $R_{C5+6}$ variants (Supplementary Table 3) were prepared as described[29]. Transfer plasmids expressing new AGO2 variants were generated from the wild-type pLJM1-3xHA-AGO2 plasmid and prepared from NEB Stable Competent *E. coli* (NEB, C3040). For each well in a 6-well plate, ~170,000 HEK293FT cells/cm$^2$ were reverse-transfected with 1.4 µg of transfer plasmid, 0.94 µg of pCMV-dR8.91 packaging plasmid (a gift from Jonathan Weissmann), and 0.47 µg of pMD2.G envelope plasmid (Addgene #12259) using Lipofectamine 2000 and Opti-MEM. After 16 h, media was refreshed. After another 48 h, media was collected and centrifuged at 500*g* for 10 min to pellet debris. To infect MEFs, 500 µL of virus-containing supernatant (~40% of the total) was combined with 1 mL media and 12 µg/mL polybrene (Santa Cruz Biotechnology, SC-134220), and then added to one well of a 6-well plate containing MEFs that were plated 24 h before at ~15,600 cells/cm$^2$, for a final 8 µg/mL polybrene. Plates were centrifuged at 1,200*g* for 1.5 h at ~30 °C, and then returned to the 37 °C incubator. The following day, cells were expanded to 10-cm plates with media containing 4 µg/mL puromycin. Puromycin selection continued for another six days, refreshing selection media every two days. Cells typically reached confluency after three days of selection and were left confluent for an additional three days.

**AGO2 intracellular co-IP assays**

Experiments testing the effect of AGO2 mutations on intracellular levels of associated miRNAs were performed as described[29]. Briefly, lentiviral production and transduction of MEFs with pLJM1-3xHA-AGO2 constructs was performed as described above. Following six days of puromycin selection (which included three days of contact inhibition), approximately 5 million cells per sample were washed with PBS, dissociated using 1X TrypLE Express Enzyme (Gibco, 12604039), washed with PBS again, and lysed with 600 µL Pierce IP Lysis Buffer (Thermo Scientific, 87788) containing 1 tablet of cOmplete Mini EDTA-free Protease Inhibitor Cocktail per 10 mL buffer. Following 15 min of end-over-end rotation in lysis buffer at 4 °C, lysate was clarified by centrifugation at 17,000*g* for 15 min at 4 °C. After setting aside a portion for immunoblotting, each clarified lysate was incubated with 50 µL Pierce anti-HA magnetic beads with end-over-end rotation at 4 °C for 2 h. Beads were then washed three times with 250 µL lysis buffer, and co-immunoprecipitating RNA was extracted with TRI Reagent added directly to the beads. Following RNA isolation, one-third of the total volume was used for northern analysis. Statistical analyses were performed in GraphPad Prism (v10.6.1), using an ordinary one-way ANOVA with Dunnett's multiple comparisons tests to compare each AGO2 variant to WT AGO2 (*$P < 0.05$, ***$P < 0.0001$; n.s., not significant). Representative northern blots are shown in Extended Data Figure 7b, and the corresponding quantification is shown in Extended Data Figure 7c. Immunoblots of AGO2 protein variants are shown in Supplementary Figure 9.

**sRNA-seq**

AGO protein, along with bound miRNAs, was isolated from MEFs and S2 cells using a FLAG-tagged T6B peptide[17], as described[18], except that a 3xFLAG-SUMO-tagged peptide (hereafter referred to as T6B-3xFLAG peptide) was used instead of the original 1xFLAG-GST. Its purification is described above. *Zswim8*-knockout and control MEF cell lines, generated as described above, were allowed to reach confluency, at which point media was refreshed and cells were left confluent for 5 days prior to harvesting. *Zswim8*-knockout and control S2 cells[29] were harvested once they reached confluency. Both MEFs and S2 cells were flash-frozen as cell pellets prior to lysis. Anti-FLAG M2 magnetic beads were pre-coupled to T6B-3xFLAG peptide, washed to remove unbound peptide, and then incubated with clarified lysate from MEFs or S2 cells that were lysed in 50 mM Tris pH 8.0, 150 mM NaCl, 5 mM EDTA, 10% (v/v) glycerol, and 0.5% (v/v) NP-40, with 1 tablet of cOmplete Mini EDTA-free Protease Inhibitor Cocktail per 10 mL buffer. Beads were washed three times with lysis buffer and once with PBS, and bound RNA was extracted directly from beads with TRI Reagent, as described above.

sRNA-seq libraries were prepared as described (http://bartellab.wi.mit.edu/protocols.html), with the following changes. sRNA-seq libraries were generated using 40% of the RNA isolated from a T6B pull-down. Instead of an 18-nt size marker, a 12-nt size marker (Supplementary Table 5) was used in all size-selection steps to capture shorter miRNA isoforms. For libraries generated from S2 cells, 2S rRNA was depleted from samples after the initial size selection using subtractive hybridization with a biotinylated antisense oligo, as described[18]. Libraries were sequenced on the Illumina NovaSeq platform with 100-nt single-end reads.

Processing of sequencing reads was done by trimming adapters with cutadapt v4.8[30], filtering out reads with a quality score of 30 or below with FASTX Toolkit v0.0.14, and then string-matching the first 13 nt of each read against a dictionary of miRNA names and sequences derived primarily from TargetScanFly7 and TargetScanMouse7[31]. For miRNAs whose sequences in the TargetScan annotations differed from those in miRGeneDB 3.0 annotations[32], which was most common for lowly expressed and/or non-conserved miRNAs, sequences from miRGeneDB were added to the mapping dictionaries as processing variants (e.g. mmu-let-7g-3p.1 and mmu-let-7g-3p.2). MicroRNAs sharing the same first 13 nucleotides were collapsed into a single entry, and the full names of the collapsed species are provided in Supplementary Table 6. The length of each mapped miRNA was recorded, generating a table of read counts for each miRNA at each size, ranging from 13 to 31 nt.

To identify miRNAs accumulating in *Zswim8*-knockout lines compared to control lines, reads associated with all isoforms of a given miRNA were summed, filtered based on a read cutoff of greater than five reads per library for at least two of the knockout lines and 30 reads across all six lines (three knockouts and three controls), and normalized for sequencing depth using the estimateSizeFactors function of DESeq2 v1.38.3[33]. Fold-changes between *Zswim8*-knockout and control lines were then estimated using summed count values with DESeq2 v1.38.3 without use of the lfcShrink function[33]. The mean of normalized abundances in the control lines was taken as the expression in WT cells. To identify miRNAs that accumulated significantly upon loss of ZSWIM8, we applied a modified BBUM model (FDR-adjusted $P$-value < 0.05)[34].

Three approaches were used to identify miRNAs that were not classified as significantly up-regulated but nonetheless were potentially ZSWIM8-sensitive, with the goal of stringently removing from subsequent analyses any miRNAs whose dominant length isoform (typically 22 nt) could be subject to TDMD. First, using the fold-changes calculated by summing the counts associated with all isoforms of a given miRNA, miRNAs with $P_{adj}$ less than 0.05, where the $P$-values were adjusted by the Benjamini-Hochberg method by DESeq2, and fold-changes ($\log_2$) greater than zero were considered potentially ZSWIM8-sensitive. Second, the fold-change of each miRNA was compared to that of the co-produced passenger miRNA derived from the same precursor hairpin, as described[35]. MicroRNAs were classified as potentially ZSWIM8-sensitive if their $\log_2$ fold-changes and those of their passenger strands were separated beyond their respective standard errors, as calculated by DESeq2, as would be expected if ZSWIM8 altered the degradation rate of the miRNA and not the production rate. For miRNAs with more than one potential passenger strand, read counts attributed to each passenger were summed, and the $\log_2$ fold-change of the summed counts was used in the comparison. For this analysis, miRNAs were not subject to the read cutoff described previously, as many passenger miRNAs were lowly expressed. Third, miRNAs in MEFs were annotated as potentially ZSWIM8-sensitive if they were previously called ZSWIM8-sensitive in mouse tissues[35], and miRNAs in S2 cells were annotated as potentially ZSWIM8-sensitive if they were called sensitive in S2 cells[29] or Drosophila embryos[36]. The isoforms of miRNAs meeting any of these three conditions were excluded from the analyses in Extended Data Figure 8.

Analogous analyses for these miRNAs whose levels are significantly or potentially affected by ZSWIM8 are shown in Supplementary Figure 13.

To investigate the relationship between miRNA length and ZSWIM8-dependent degradation, fold-changes observed upon loss of ZSWIM8 were calculated for individual length isoforms of the remaining miRNAs that had not been excluded due to potential ZSWIM8 sensitivity of their major isoforms. To this end, read counts corresponding to each isoform were first normalized using the size factors previously computed based on the summed counts, and fold-changes ($\log_2$) were calculated with DESeq2. Finally, the median $\log_2$ fold-change of all 22-nt species was calculated and subtracted from the $\log_2$ fold-changes of all species. This normalization centers the distribution of $\log_2$ fold-changes for the 22-nt isoforms at zero, as would be expected if it were not sensitive to loss of ZSWIM8.
