## Supplementary_Table_2 for "The E3 ubiquitin ligase mechanism specifying target-directed microRNA degradation"

### Cryo-EM data collection, refinement and validation statistics

| | Map A ZSWIM8-CUL3 complex bound to AGO2-miR-7-CYRANO, | Map B Locally refined interactions of ZSWIM8-CUL3 complex bound to AGO2-miR-7-CYRANO | Map C focused map of ZSWIM8-CUL3 complex bound to AGO2-miR-7-CYRANO | Map D Locally refined map of ZSWIM8-CUL3 complex bound to AGO2-miR-7-CYRANO | Map E Composite map of ZSWIM8-CUL3 complex bound to AGO2-miR-7-CYRANO |
|---|---|---|---|---|---|
| **Data collection and processing** | | | | | |
| Magnification | 105,000 | 105,000 | 105,000 | 105,000 | 105,000 |
| Voltage (kV) | 300 | 300 | 300 | 300 | 300 |
| Electron exposure (e–/Å$^2$) | 58 | 58 | 58 | 58 | 58 |
| Defocus range (μm) | -0.5 – -2.0 | -0.5 – -2.0 | -0.5 – -2.0 | -0.5 – -2.0 | -0.5 – -2.0 |
| Pixel size (Å) | 0.8512 | 0.8512 | 0.8512 | 0.8512 | 0.8512 |
| Symmetry imposed | C1 | C1 | C1 | C1 | C1 |
| Initial particle images (no.) | 5,812,715 | 5,812,715 | 5,812,715 | 5,812,715 | 5,812,715 |
| Final particle images (no.) | 234,181 | 234,181 | 234,181 | 234,181 | 234,181 |
| Map resolution (Å) | 3.1 | 3.1 | 3.2 | 3.2 | 3.1 |
| FSC threshold | | | | | |
| Map resolution range (Å) | 2.8-3.3 | 2.7-3.2 | 2.8-3.3 | 2.8-3.4 | 2.8-3.3 |
| **Refinement** | | | | | |
| Initial model used (PDB code) | | | | | Alphafold3, AGO2 (6NIT), CUL3 (5NLB), ELOB/C (1LM8) |
| Model resolution (Å) | | | | | 3.1 |
| FSC threshold | | | | | |
| Model resolution range (Å) | | | | | |
| Model composition | | | | | |
| Non-hydrogen atoms | | | | | 28,597 |
| Protein residues | | | | | 3,609 |
| Nucleotides | | | | | 50 |
| Ligands | | | | | 2 Zn |
| *B* factors (Å$^2$) | | | | | |
| Protein | | | | | 87.42 |
| Nucleotides | | | | | 103.12 |

| | |
|---|---|
| Ligand | 97.48 |
| R.m.s. deviations | |
| Bond lengths (Å) | 0.007 |
| Bond angles (°) | 0.603 |
| Validation | |
| MolProbity score | 2.02 |
| Clashscore | 5.75 |
| Poor rotamers (%) | 2.10 |
| Ramachandran plot | |
| Favored (%) | 92.77 |
| Allowed (%) | 7.14 |
| Disallowed (%) | 0.08 |
